# Systematic characterization of the ovarian landscape across mouse menopause models

**DOI:** 10.1101/2025.06.10.658920

**Authors:** Minhoo Kim, Rajyk Bhala, Justin Wang, Xucheng Liang, Mingfei Chen, Ryan J. Lu, Evelyn H. Lee, Julio L. Alvarenga, Rapheal G. Williams, Bérénice A. Benayoun

## Abstract

Menopause affects not only fertility but also systemic health. Yet mechanisms underlying this coupling are still poorly understood, partly due to the absence of robust, age-relevant preclinical models with comprehensive molecular and phenotypic characterization. To address this, we systematically compared three candidate mouse models of menopause: (1) intact aging, (2) chemical ovarian follicle depletion using 4-vinylcyclohexene diepoxide (VCD) administered at multiple ages, and (3) *Foxl2* haploinsufficiency, a genetic model based on a gene linked to human premature ovarian failure. Through histology, serum hormone profiling, single-cell transcriptomics and machine-learning approaches, we identify both shared and model-specific features of follicle loss, endocrine disruption, and transcriptional remodeling. Our comparative framework enables informed selection of context-appropriate preclinical rodent models to study menopause and the broader physiological consequences of ovarian aging.

**Teaser:** This systematic characterization of mouse menopause models lays the groundwork for future preclinical work on the systemic impacts of menopause.

## Introduction

Women are born with a finite oocyte reserve, and its progressive depletion with age leads to the irreversible loss of ovarian function, known as menopause (*1–3*). Epidemiological studies link later age-at-menopause with increased lifespan and reduced risk of age-related diseases (*e.g*. cardiovascular disease, osteoporosis, and neurodegeneration) (*4–6*). Conversely, menopause is accompanied by a sharp rise in age-associated pathologies, underscoring widespread consequences of ovarian aging (*7*). Despite its public health relevance, how ovarian decline contributes to systemic aging is still poorly understood, in part due to the dearth of well-characterized, aging-relevant preclinical models.

Menopause is characterized by endocrine reprogramming that extends beyond the reproductive system (*8*). As follicular reserves decline, reduced steroidogenic activity disrupts feedback regulation of the hypothalamic-pituitary-gonadal (HPG) axis, leading to sustained hormonal shifts (*8*). Circulating estrogen and anti-Müllerian hormone (AMH) fall to near-undetectable levels, while follicle-stimulating hormone (FSH) and inhibin A (INHBA) rise (*9–11*). These changes influence multiple somatic systems, including metabolic, skeletal, and central nervous systems (*12, 13*). However, dissecting the mechanisms linking ovarian dysfunction to systemic physiology requires robust preclinical models recapitulating key biological features of menopause.

The most frequently used rodent models capture menopausal physiology only imperfectly (*14*). Although intact aging mice are widely used, they do not undergo true menopause; instead, they enter a state of “estropause,” retaining low but detectable estrogen levels and lacking the postmenopausal endocrine milieu observed in postmenopausal women (*15, 16*). Although ovariectomy (OVX) achieves rapid estrogen depletion, it lack a gradual endocrine transition, and it depletes residual post-reproductive ovarian tissue, which is known to exert androgenic and paracrine effects (*17*). Thus, while OVX models acute hormone loss, it incompletely models the progressive ovarian decline characteristic of menopause. Repeated injections with 4-vinylcyclohexene diepoxide (VCD) leads to dramatic follicle depletion through selective targeting of small, pre-antral follicles, inducing progressive ovarian failure while preserving a post-reproductive ovarian tissue (*15*). This approach produces estrous acyclicity, estrogen deficiency, and compensatory FSH elevation, resembling several endocrine features of human menopause (*18, 19*). However, prior studies have primarily applied VCD to young adult mice (∼2-3 months), limiting insight into how the timing of ovarian failure intersects with organismal aging (*20, 21*). Because aging alters metabolic, immune, and endocrine states in profound ways (*22–25*), understanding menopausal biology requires evaluating ovarian decline in the context of middle-aged biology.

Genetic models informed by human physiology can also provide useful tools to understand ovarian decline and its systemic impacts. Haploinsufficiency of *Foxl2*, a gene encoding a transcription factor essential for granulosa cell identity and ovarian maintenance (*26–29*), is causally linked to premature ovarian insufficiency in humans (*30–32*). In adult mice, *Foxl2* expression is primarily restricted to ovarian granulosa cells and pituitary gonadotroph-lineage cells (*33*). While complete *Foxl2* knockout leads to severe developmental defects and perinatal lethality, heterozygous carriers have been incidentally reported to be subfertile, suggestive of milder ovarian dysfunction that may better recapitulate menopausal transition (*28, 29*). Unlike surgical or chemical models, *Foxl2* haploinsufficiency represents a physiologically relevant genetic perturbation that enables investigation of early and potentially subclinical features of ovarian decline without exogenous insult. Rather than modeling overt menopausal hormone depletion alone, this model may provide insight into early trajectories of ovarian dysfunction that precede systemic endocrine collapse.

Collectively, these models could help recapitulate distinct mechanisms of ovarian decline in a preclinical setting. However, differential impact of VCD with age-at-exposure and *Foxl2* haploinsufficiency have not been systematically characterized and benchmarked within a unified framework. Such comparative characterization is essential for interpreting prior literature and for selecting appropriate models tailored to understanding systemic impacts of menopause-like ovarian failure.

Here, we present a comprehensive resource evaluating three complementary mouse models of ovarian aging: (1) intact aging, (2) VCD-induced follicle depletion initiated at multiple adult ages, and (3) *Foxl2* haploinsufficiency. Our objective is to systematically benchmark shared and divergent features across multiple biological scales. We integrate histological profiling, endocrine measurements, and single-cell transcriptomics to characterize ovarian structural, functional, and molecular trajectories. Additionally, we develop hormone-based and transcriptome-based ovarian aging clocks to provide quantitative measures of reproductive aging across models. This comparative framework establishes a foundational resource to guide selection and interpretation of menopause models and to support future studies investigating how ovarian dysfunction influences systemic aging trajectories.

## Results

### A systematic framework for modeling menopause in mice

To systematically assess and compare physiologically relevant menopause models, we characterized three rodent models that represent distinct mechanisms of ovarian dysfunction: (1) the intact aging model (“Aging model”), (2) a chemically-induced model using repeated intraperitoneal injections of 4-vinylcyclohexene diepoxide (VCD, “VCD model”) at various age-of-exposure (*15, 34*), and (3) a genetic model of *Foxl2* haploinsufficiency (“*Foxl2* +/− model”) (*26*). Our aim was to establish a comparative framework to evaluate these models in terms of histological, hormonal, and integrated ovarian health outcomes to evaluate their suitability for modeling key aspects of ovarian aging (Fig. 1A). The Aging model involved comparison of young adult (4 months) and old (20 months) female C57BL/6JNia mice. We selected these timepoints to capture the transition from reproductive to firmly post-estropausal states, as C57BL/6 females typically enter estropause ∼12 months of age and show pronounced ovarian aging phenotypes by 20 months (*35, 36*). The VCD model included C57BL/6J female mice injected with vehicle control (safflower oil, hereafter referred to as “CTL”) or VCD starting at 3, 6, 8, or 10 months of age. This scheme allowed us to examine how the timing of ovarian insult (potentially modeling varying “age-at-menopause”) could impact ovarian physiology, particularly when VCD exposure occurs closer to natural estropause (*35*). Importantly, incorporating multiple age-at-injection paradigms should help disentangle ovarian-intrinsic failure mechanisms and how it interacts with the aging milieu. The *Foxl2* haploinsufficiency model compared *Foxl2^+/+^* and *Foxl2^+/−^* female littermate mice from our transgenic colony at young (3-4 months), early middle-age (8-10 months) and later middle-age (12-13 months) to test whether partial loss of *Foxl2* leads to anticipated occurrence of age-associated ovarian phenotypes compared to wild-type control mice. We selected the heterozygous model because complete *Foxl2* loss is associated with severe developmental defects and high perinatal lethality (*28*), and heterozygous mutations of FOXL2 are sufficient to drive both syndromic and isolate premature ovarian failure in humans patients.

**Fig. 1.**
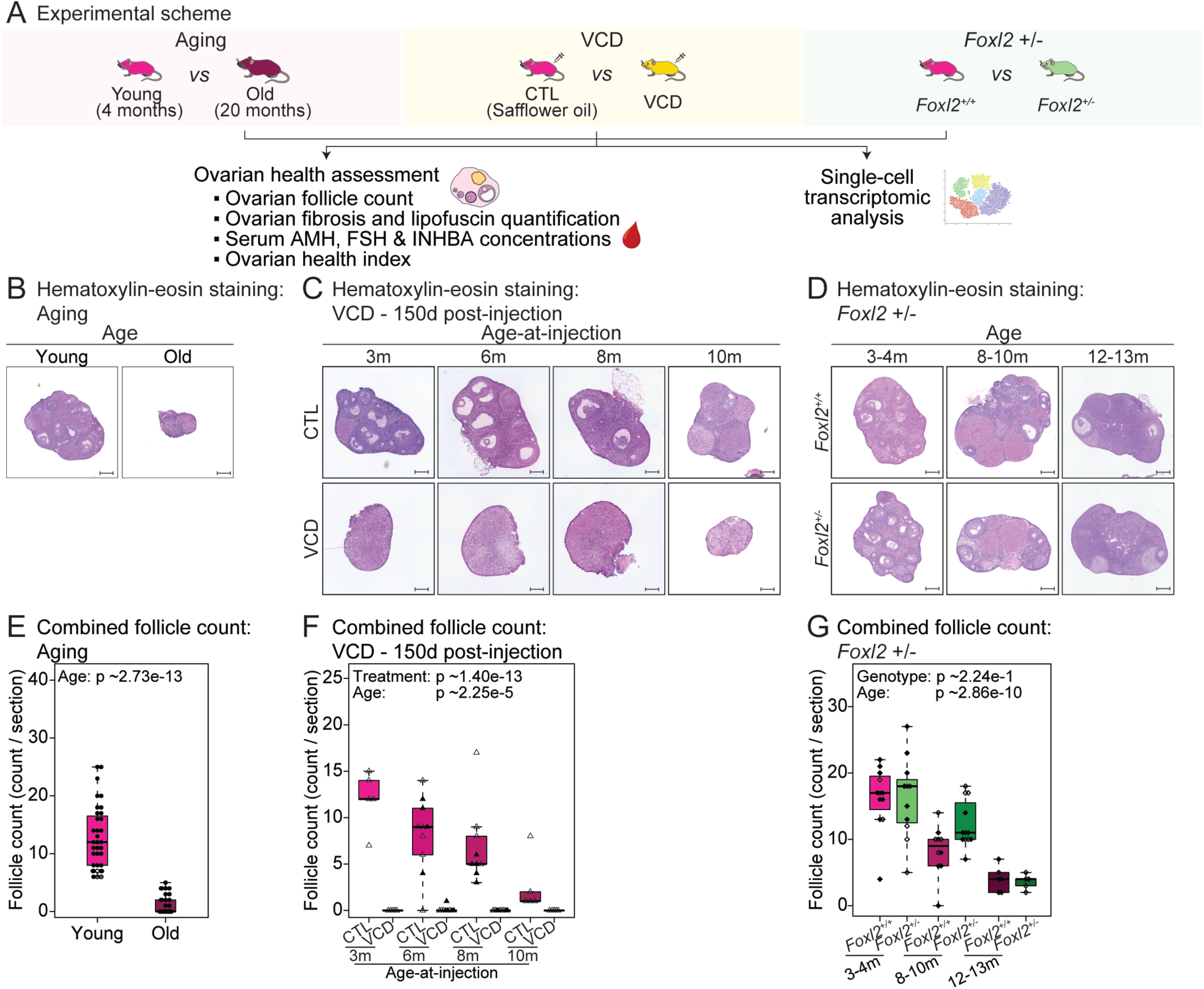
Ovarian follicle count quantification of animals from Aging, VCD and *Foxl2* haploinsufficiency models. **(A)** Schematic of the experimental design. **(B-D)** Representative hematoxylin-eosin staining images of ovarian tissues from Aging, VCD, and *Foxl2* haploinsufficiency model mice. For the VCD model, ovarian tissues were analyzed 5 months post-injection. Scale bar: 250 µm. **(E-G)** Combined follicle counts (primordial, primary, secondary and antral follicles, and corpus luteum) from Aging (n = 25 per group), VCD (n = 5-10 per group), and *Foxl2* haploinsufficiency (n = 11-24 per group) model mice. Statistical significance was assessed using Wilcoxon test for the Aging model and aligned rank transform ANOVA to test effects for the VCD and *Foxl2* haploinsufficiency models; p-values are shown. Open data points indicate data used for ovarian health index calculation.

To evaluate the impact of *Foxl2* haploinsufficiency on ovarian aging phenotypes, we generated a knock-out *Foxl2* allele using mice carrying a regenerated floxed *Foxl2* locus (*26*), which allowed us to generate mice carrying a constitutive deletion of *Foxl2*’s single exon (fig. S1A; see Materials and Methods). Importantly, RT-qPCR analysis confirmed that *Foxl2* expression was significantly reduced in *Foxl2^+/−^*ovaries compared to *Foxl2^+/+^* controls (genotype effect, p-value ∼0.0206; fig. S1B). Additionally, fertility outcomes for *Foxl2^+/+^* and *Foxl2^+/−^*females from our breeding colony confirmed previous reports of decreased fertility of *Foxl2* haploinsufficient mice (figs. S1C,D and table S1; see Materials and Methods). Specifically, adult *Foxl2^+/−^* females showed significantly smaller first litters (p-value ∼0.0161; fig. S1C), and increased latency to first time to delivery after male pairing compared to matched *Foxl2^+/+^* female controls (p-value ∼0.0260; fig. S1D), suggesting impaired reproductive potential consistent with ovarian dysfunction or premature ovarian aging.

### Histological and tissue-level remodeling across mouse menopause models

A decline in ovarian follicle numbers is a well-established hallmark of ovarian aging and menopause (*37*). Thus, we performed histological analysis of ovarian follicle populations using hematoxylin and eosin (H&E) staining (Figs. 1B-G and fig. S2A). As expected, the Aging model showed a significant reduction in follicles at all developmental stages in old females, including primordial, primary, secondary, and antral follicles, as well as corpora lutea (Figs. 1B,E and fig. S2B), consistent with the established progressive follicular depletion that accompanies natural ovarian aging (*38, 39*). At 5 months post-injections, VCD-treated animals also showed significant depletion of follicle numbers, regardless of age-at-injection (Figs. 1C,F and fig. S2C). To note, due to the terminal nature of histological analysis, ovaries were collected for H&E staining at a single 5 months post-injection time point. This endpoint was selected to align with the conclusion of longitudinal serum hormone profiling (see below). In contrast, we did not observe any significant consistent differences in combined follicle counts between *Foxl2^+/+^* and *Foxl2^+/−^* ovaries (Figs. 1D,G and fig. S2D).

Next, we leveraged Picrosirius Red and Sudan Black B staining of ovarian tissue to quantify fibrosis and lipofuscin accumulation, respectively (Fig. 2A). Previous studies have demonstrated increased fibrosis and lipofuscin deposition in aged ovaries, reflecting progressive remodeling with ovarian aging (*40–42*). Consistently, ovaries from the Aging model exhibited significant increases in both fibrosis and lipofuscin accumulation in old females compared to young controls (p ∼0.0190 and p ∼0.00433, respectively; Figs. 2B,E,H,K). VCD-treated animals similarly showed significant increases in both fibrosis and lipofuscin staining relative to vehicle-treated controls, indicating that chemical follicle depletion is accompanied by stromal remodeling and accumulation in a way consistent with accelerated ovarian aging (treatment effect, p ∼0.0242 and p ∼0.00776, respectively; Figs. 2C,F,I,L). In the *Foxl2* haploinsufficiency model, despite the absence of overt follicle depletion, genotype was significantly associated with increased fibrosis and lipofuscin accumulation (genotype effect, p ∼0.00279 and p ∼0.0213, respectively; Figs. 2D,G,J,M), consistent with premature aging of the ovarian stroma. These findings indicate that structural remodeling features can emerge independently of substantial reductions in follicle number, highlighting both shared and model-specific aspects of ovarian aging phenotypes.

**Fig. 2.**
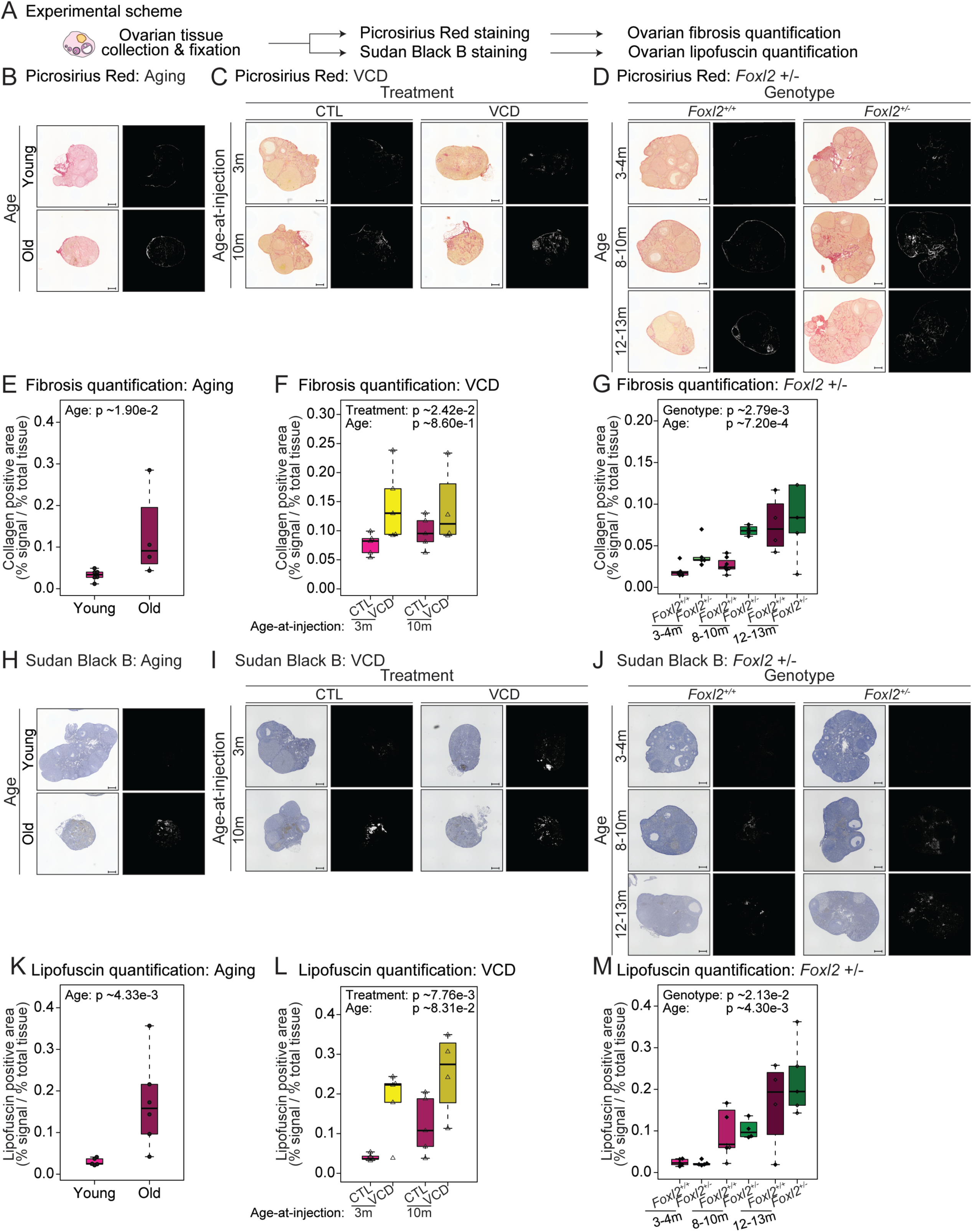
Quantification of ovarian fibrosis and lipofuscin accumulation across Aging, VCD, and *Foxl2* haploinsufficiency models. **(A)** Schematic of the experimental design. Picrosirius Red and Sudan Black B staining were performed and quantified to assess fibrosis and lipofuscin accumulation in the ovaries. **(B-D)** Representative Picrosirius Red staining images of ovarian tissues from Aging, VCD, and *Foxl2* haploinsufficiency model mice. Original and processed image are shown. **(E-G)** Quantification of percent fibrotic area across Aging (n = 6 and 4 for Young and Old, respectively), VCD (n = 4-5 per group), and *Foxl2* haploinsufficiency (n = 4-7 per group) model mice. **(H-J)** Representative Sudan Black B staining images showing lipofuscin accumulation in ovarian tissues from Aging, VCD, and *Foxl2* haploinsufficiency model mice. Original and processed image are shown. **(K-M)** Quantification of percent lipofuscin-positive area across Aging (n = 5 and 6 for Young and Old, respectively), VCD (n=4-5 per group), and *Foxl2* haploinsufficiency (n = 4-7 per group) model mice. For the VCD model, ovarian tissues were analyzed 5 months post-injection. Scale bar: 200 µm. For (**E-G** and **K-M**), statistical significance was assessed using the Wilcoxon test for the Aging model and aligned rank transform ANOVA to test effects for the VCD and *Foxl2* haploinsufficiency models; p-values are shown.

### Comparative endocrine alterations across mouse menopause models

To evaluate how each model perturbs endocrine regulation, we measured serum levels of AMH, FSH and INHBA, key markers of ovarian reserve and HPG axis regulation (*43, 44*) (Fig. 3A and table S2). Estradiol was not quantified due to known technical limitations affecting assay reliability in rodent samples (see Discussion). To note, due to a change of FSH quantitation assays offered at the UVA ligand core over the course of our study, FSH measurements were harmonized across methods using a polynomial correction model (fig. S3A; see Materials and Methods).

**Fig. 3.**
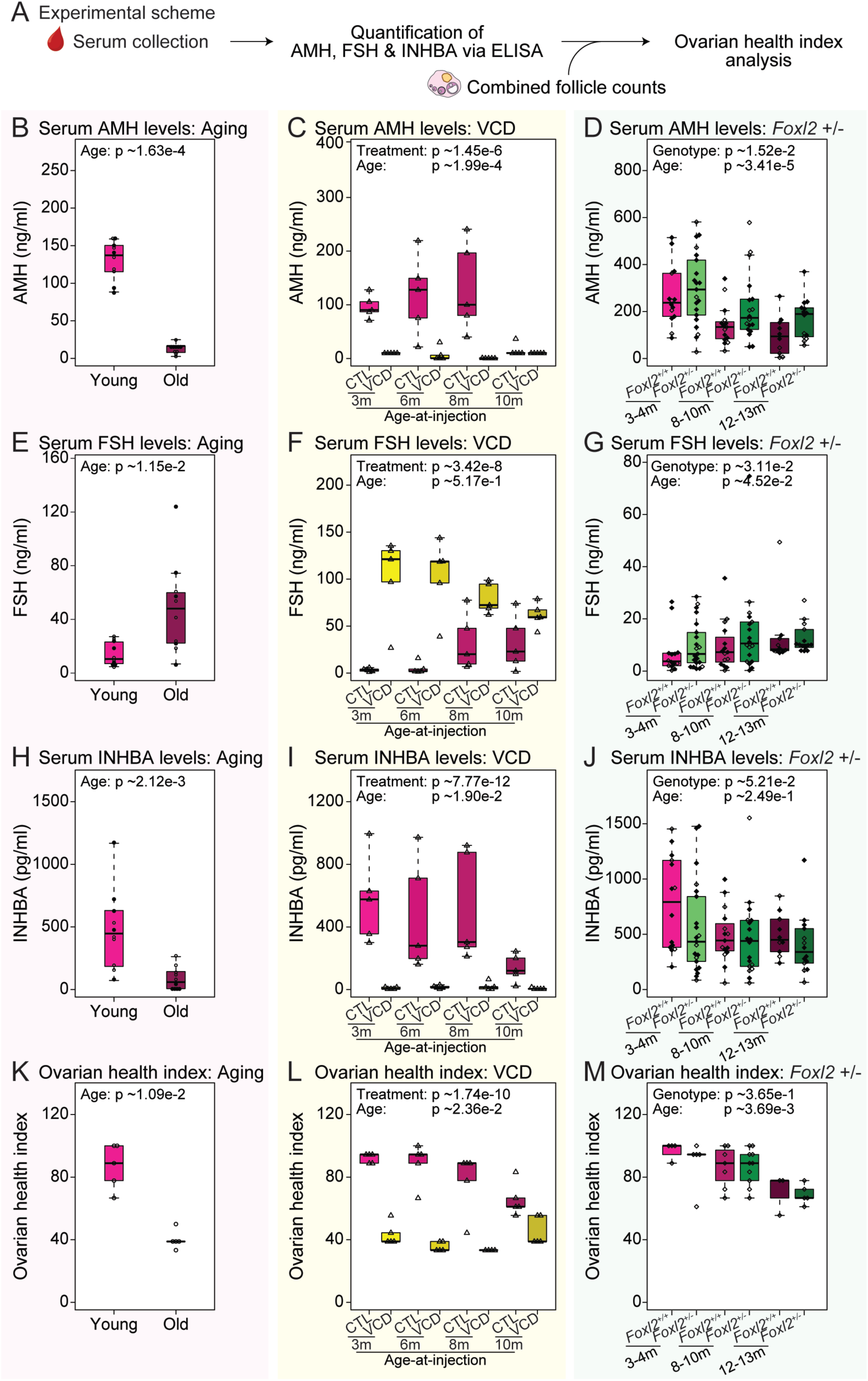
Ovarian hormone marker quantification of animals from Aging, VCD and *Foxl2* haploinsufficiency models. **(A)** Schematic of the experimental design. Serum levels of AMH, FSH and INHBA were quantified via ELISA. Serum hormone levels and combined follicle counts were integrated to calculate the ovarian health index. **(B-D)** Serum AMH level quantification from Aging (n = 10 per group), VCD (n = 5 per group), and *Foxl2* haploinsufficiency (n = 10-21 per group) model mice. **(E-G)** Serum FSH level quantification from Aging (n = 10 per group), VCD (n = 5 per group), and *Foxl2* haploinsufficiency (n = 10-21 per group) model mice. **(H-J)** Serum INHBA level quantification from Aging (n = 10 per group), VCD (n = 5 per group), and *Foxl2* haploinsufficiency (n = 10-21 per group) model mice. **(K-M)** Ovarian health index from Aging (n = 5 per group), VCD (n = 4-5 per group), and *Foxl2* haploinsufficiency (n = 4-10 per group) model mice. Statistical significance was assessed using Wilcoxon test for the Aging model and aligned rank transform ANOVA to test effects for the VCD and *Foxl2* haploinsufficiency models; p-values are shown. Open data points indicate data used for ovarian health index calculation.

In the Aging model, old females showed expected decreases in circulating AMH and INHBA and increased circulating FSH compared to young females, consistent with diminished ovarian reserve (*44, 45*) (Figs. 3B,E,H). A similar pattern was observed in the VCD model, whereby AMH and INHBA were decreased and FSH was increased compared to the vehicle control groups at 5 months post-injection timepoint, regardless of age-at-injection (Figs. 3C,F,I). To enable a comprehensive analysis of endocrine responses across reproductive life stages, we also performed longitudinal hormone profiling over a five-month period following VCD exposure at various ages (3, 6, 8 and 10 months; figs. S3B-D). Interestingly, VCD injections at older ages resulted in attenuated hormonal shifts compared to younger counterparts, suggesting that ovarian or systemic factors at midlife modulate sensitivity to VCD-induced insult to ovarian reserve. In the *Foxl2^+/−^* mice, we observed conflicting hormonal trends: AMH levels were increased (consistent with younger follicle pools), whereas FSH levels were elevated and INHBA levels were reduced (consistent with older follicle pools) compared to wild-type controls (Figs. 3D,G,J and fig. S3A). To note, *FOXL2* has been shown to interact with *AMH* to modulate follicle recruitment in humans (*46*). Thus, the elevated AMH levels in the *Foxl2^+/−^*animals likely reflect altered gene regulatory networks driven by *Foxl2* deficiency, rather than preservation or rejuvenation of ovarian reserve (see Discussion).

Next, we applied our previously described composite ovarian health index, which integrates information from follicle counts and serum hormone levels to yield a holistic measure of ovarian health (*36*) (Figs. 3K-M). As expected, the index was significantly lower in old *vs*. young animals and VCD-treated animals vs. CTL (regardless of age-at-injection), reflecting impaired ovarian health (Figs. 3K,L). In contrast, *Foxl2^+/−^* mice did not differ significantly from wild-type animals (Fig. 3M), likely due to elevated AMH levels.

Together, these results provide a comprehensive characterization of histological and endocrine features of three mechanistically distinct mouse menopause models. Both the Aging and VCD models exhibit robust phenotypic hallmarks of ovarian decline, including depletion of follicular reserves, tissue remodeling, and disrupted endocrine profiles. In contrast, the *Foxl2* haploinsufficiency model displays a distinct phenotype characterized by preserved follicle counts and elevated AMH levels, but with anticipated aged-like stromal remodeling and reproductive impairment. These findings indicate that the three models capture overlapping yet non-identical dimensions of ovarian aging biology, underscoring the importance of selecting models based on specific structural, endocrine, or regulatory processes under investigation.

### A hormone-based clock, “OvAge”, for non-terminal prediction of ovarian aging

Although the ovarian health index is a useful metric of ovarian health, its partial reliance on post-mortem histological measurements limits its applicability in longitudinal studies. Thus, we aimed at developing a complementary, non-terminal metric of ovarian aging, using a hormone-based predictive machine-learning model, which we termed “OvAge” (Fig. 4A). By integrating information on AMH, FSH, and INHBA levels, we sought to develop a model that provides a more comprehensive assessment of endocrine function rather than evaluating each hormone individually. To increase the generalizability of the clock, we trained a random forest regression model using serum hormone data (AMH, FSH, and INHBA) from animals with intact ovarian function and no experimental perturbation, including animals from the Aging model, vehicle control (CTL) groups from the VCD model, *Foxl2^+/+^* animals from the *Foxl2* haploinsufficiency model. In addition, to further increase statistical power, we also took advantage of similar data from *Fshr^+/+^* animals derived from our previously published study of *Fshr* haploinsufficient mice (*47*). This hormone level data was then randomly partitioned into distinct training (n = 212) and testing (n = 68) sets to enable robust evaluation of model performance.

**Fig. 4.**
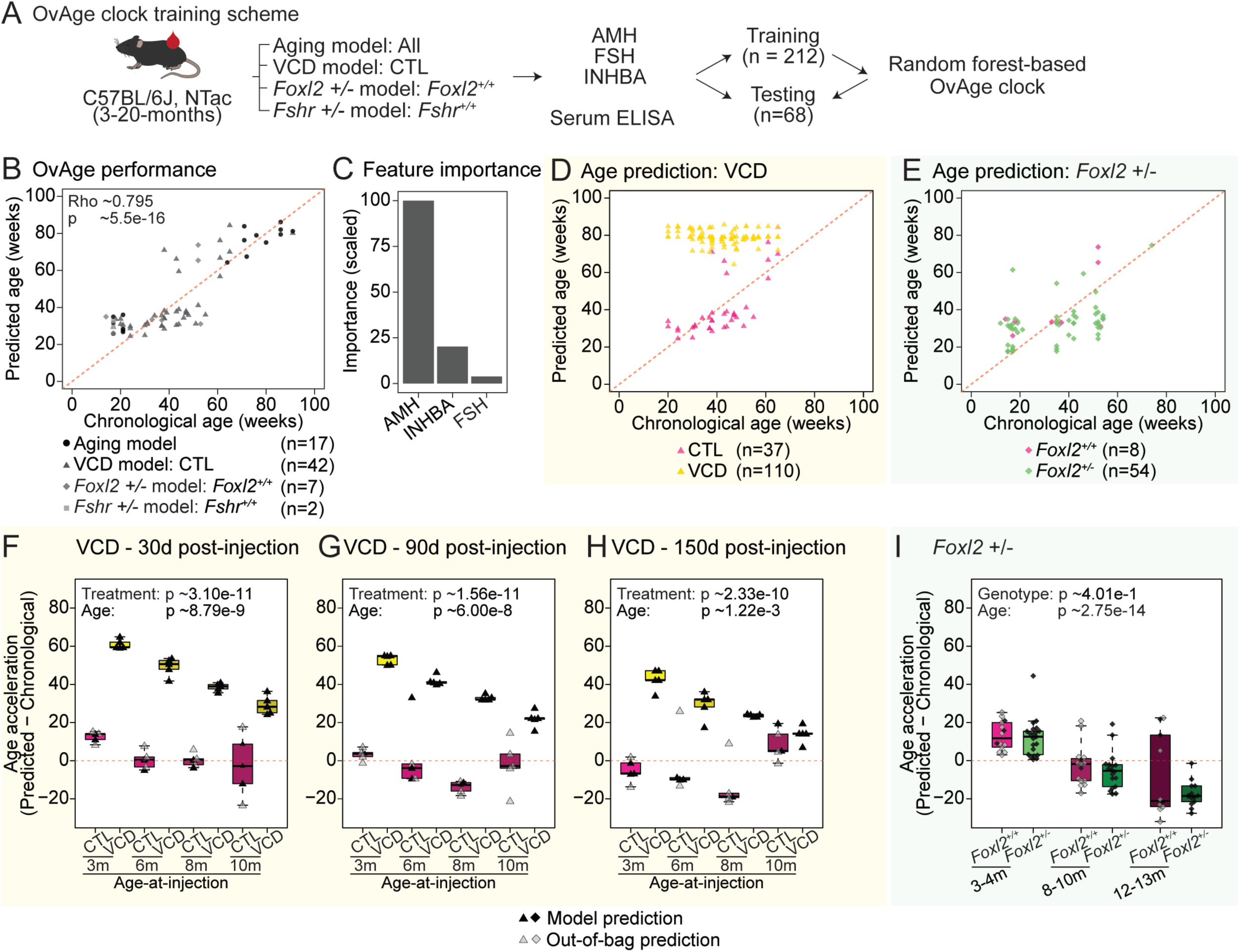
Development and application of OvAge, a hormone-based ovarian aging clock. **(A)** Schematic representation of the OvAge clock development pipeline. Data from all animals from the Aging model, vehicle control (CTL) from the VCD model, *Foxl2^+/+^* from the *Foxl2* haploinsufficiency model, and *Fshr^+/+^* from the *Fshr* haploinsufficiency model mice were used to train and test OvAge. **(B)** Scatter plot comparing OvAge-predicted and chronological ages, reported in weeks, in the test dataset (n = 68). **(C)** Bar plot of scaled feature importance for OvAge model. **(D,E)** Scatter plots comparing OvAge-predicted and chronological ages for VCD (n = 147) and *Foxl2* haploinsufficiency (n = 62) model mice. **(F-H)** Age acceleration analysis for VCD model animals at 30-, 90- and 150-days post-injection (n = 4-5 per group). **(I)** Age acceleration analysis for *Foxl2* haploinsufficiency animals (n = 10-21 per group). Age acceleration was measured as the difference between predicted OvAge and chronological age. Open data points indicate data from out-of-bag predictions. For **(F-I)**, statistical significance was assessed using Wilcoxon test for the Aging model and aligned rank transform ANOVA to test effects for the VCD and *Foxl2* haploinsufficiency models; p-values are shown.

Model performance evaluation showed strong agreement between predicted OvAge and chronological age (Spearman’s Rho ∼0.795; p-value ∼5.5 × 10^−16^), confirming successful model construction (Fig. 4B). Additionally, feature importance analysis revealed that AMH contributed most strongly to age prediction, followed by INHBA and FSH (Fig. 4C). We then calculated OvAge using hormonal measurements from VCD-injected and *Foxl2^+/−^* animals (Figs. 4D,E). In the VCD model, predicted ovarian age was consistently higher than chronological age across all age-at-injection groups (Fig. 4D), suggestive of accelerated endocrine aging. In contrast, predicted ages of *Foxl2^+/−^* animals did not substantially differ from their wild-type counterparts (Fig. 4E), likely due to the importance of AMH levels in OvAge calculation.

To quantify the degree of divergence between predicted and chronological age, we computed hormonal age acceleration, defined as the difference between OvAge-predicted and chronological age (Figs. 4F-I). For the VCD model, we analyzed animals at 30-, 90-, and 150-days post-injection. In the VCD model, predicted ovarian age exceeded chronological age following VCD exposure (Figs. 4F-H). Analysis revealed significant treatment effects on age acceleration at 30, 90, and 150 days post-injection. Post hoc analyses confirmed significant increases in age acceleration at most timepoints; however, animals injected at 10 months of age did not exhibit a significant difference at 150 days post-injection (adjusted p-value ∼0.142; table S3). This pattern may reflect convergence of ovarian aging trajectories between control and VCD-treated animals as they approach estropausal onset (∼12-15 months in C57BL/6 mice (*35*)). In contrast, *Foxl2* haploinsufficient animals did not show a consistent increase in OvAge or age acceleration (genotype effect, p-value ∼0.401; Fig. 4I). This lack of hormonal age acceleration, despite evidence of stromal remodeling and reproductive impairment, suggests that *Foxl2* haploinsufficiency does not recapitulate the endocrine aging trajectory captured by OvAge, but instead may reflect a distinct regulatory perturbation of ovarian physiology.

Together, our OvAge analysis captures divergent ovarian aging trajectories in response to distinct biological perturbations. The clock recapitulates expected aging patterns in naturally aged animals, detects accelerated ovarian aging following VCD exposure, and reveals a distinct endocrine trajectory in the *Foxl2* haploinsufficiency model. These findings support the utility of OvAge as a non-terminal framework for assessing ovarian aging and highlight model-specific differences that may inform the study of reproductive senescence. To facilitate broader use of this framework, we developed an interactive online R Shiny application that enables researchers to estimate ovarian age using serum hormone measurements (https://minhooki.shinyapps.io/OvAge_Predictor/).

Importantly, although the *Foxl2* haploinsufficiency model did not exhibit overt ovarian dysfunction based on ovarian health index or OvAge (Figs. 3M, 4I), the absence of endocrine age acceleration does not exclude underlying molecular perturbations. Indeed, *Foxl2^+/−^* animals displayed significant accumulation of ovarian fibrosis and lipofuscin (Figs. 2D,G,J,M), histological hallmarks associated with tissue aging and cell stress. Together with the established role of *FOXL2* in granulosa cell identity and ovarian maintenance, its association with premature menopause in humans (*30, 31, 48*), and the reproductive impairment phenotypes observed in our cohort (Fig. S1C, D), our findings suggest that *Foxl2* haploinsufficiency likely disrupts ovarian tissue homeostasis prior to overt depletion of follicular pools, deserving further investigation of the accompanying transcriptional landscape across ovarian cell types.

### Single-cell transcriptomic profiling of ovarian cell types across menopause models

To characterize the ovarian transcriptional landscapes across candidate mouse menopause models, we performed single-cell RNA-sequencing (scRNA-seq) on ovaries collected from the Aging, VCD, and *Foxl2* haploinsufficiency models (Fig. 5A). For the Aging model, samples were collected from young (4 months) and old, estropausal (20 months) animals to capture transcriptional changes associated with intact aging (Fig. 5B). For the VCD model, we focused on two age-at-injection groups, 3 and 10 months, to capture the extremes of VCD responsiveness observed in prior analyses, as well as two post-injection time points, 30 and 90 days, based on differential effects on age acceleration at these stages (Fig. 5B). For the *Foxl2* haploinsufficiency model, ovaries were collected from young adult (4 months), middle-age (9 months) and old (17 months) *Foxl2^+/+^* and *Foxl2^+/−^*animals, enabling the assessment of gene expression changes driven by partial loss of *Foxl2* function across adulthood (Fig. 5B). Indeed, we reasoned that inclusion of a later time point for the *Foxl2* haploinsufficiency model should help amplify regulatory impact and thus help reveal clearer trends in transcriptional remodeling. The resulting datasets yielded 15,800, 47,386, and 39,110 high-quality single cells from the Aging, VCD, and *Foxl2* haploinsufficiency models, respectively (table S4). Ovarian cell populations were defined through dimensionality reduction, unsupervised clustering, and annotation using both established marker genes and automated reference mapping to publicly available single-cell datasets (Figs. 5C-H and table S5; see Materials and Methods).

**Fig 5.**
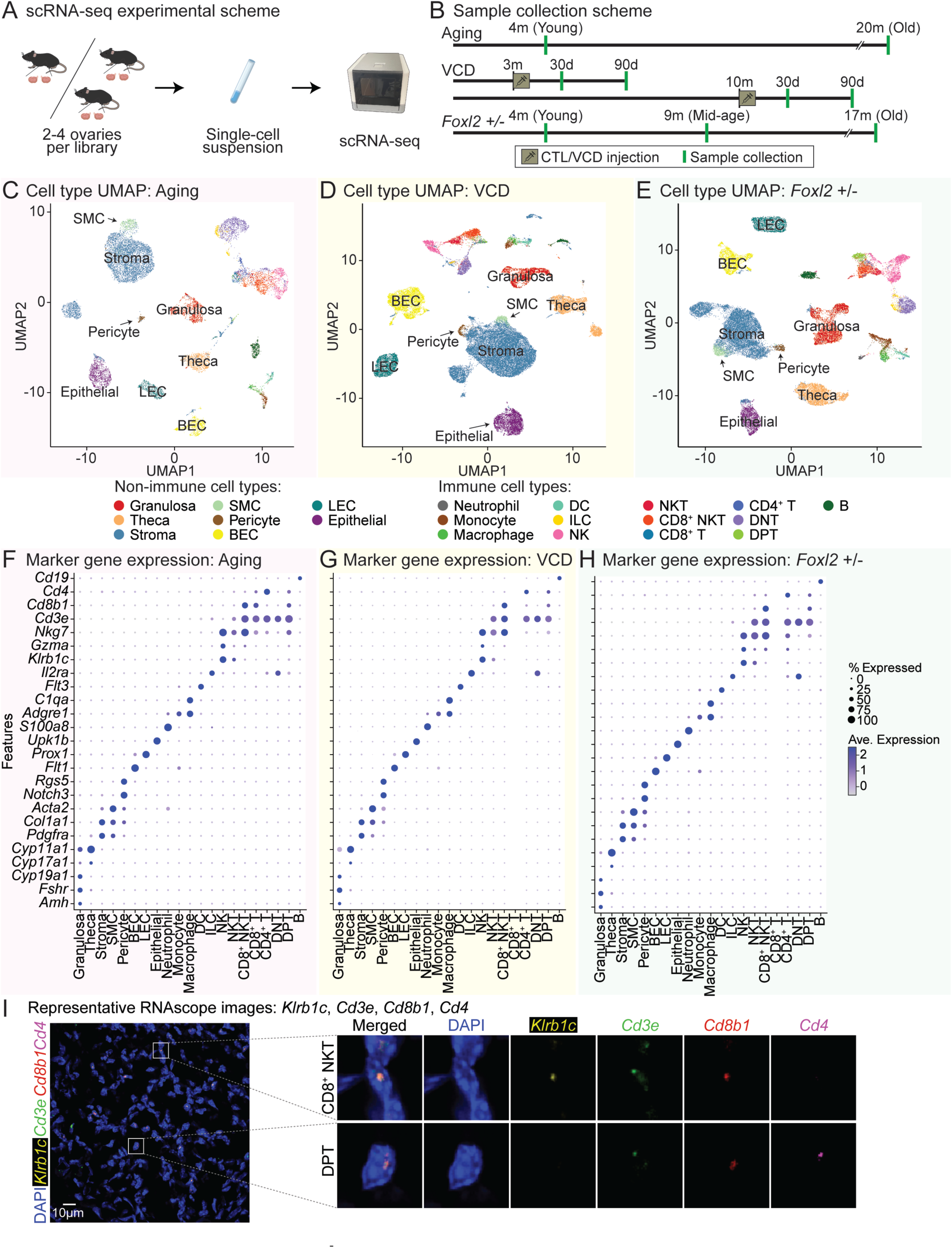
scRNA-seq profiling of ovaries from Aging, VCD, and *Foxl2* haploinsufficiency model mice. **(A)** Schematic of the experimental design. **(B)** Schematic of sample collection timepoints. **(C-E)** UMAP plots of scRNA-seq datasets from Aging, VCD and *Foxl2* haploinsufficiency models. **(F-H)** Dotplot representation of expression of marker genes of cell types detected in scRNA-seq datasets from Aging, VCD and *Foxl2* haploinsufficiency models. **(I)** Representative RNAscope images of DAPI, *Klrb1c*, *Cd3e*, *Cd8b1* and *Cd4* probes. Images were enhanced for visualization. The shown image is from a middle-aged *Foxl2^+/+^*animal.

Importantly, we identified all major ovarian cell types in our datasets across all models (Figs. 5C-H). Among the abundant non-immune populations, granulosa cells (*Amh, Fshr, Cyp19a1*), theca cells (*Cyp17a1, Cyp11a1*), stromal cells (*Pdgfra, Col1a1*), smooth muscle cells (SMCs; *Acta2*), pericytes (*Notch3*, *Rgs5*), blood endothelial cells (BECs; *Flt1*), lymphatic endothelial cells (LECs; *Prox1*), and epithelial cells (*Upk1b*) were identified. The immune compartment included neutrophils (*S100a8*), monocytes (*Adgre1*), macrophages (*C1qa*), dendritic cells (*Flt3*), innate lymphoid cells (ILCs; *Il2ra*), natural killer (NK) cells (*Klrb1c, Gzma, Nkg7*), NKT cells (*Cd3e, Klrb1c, Gzma, Nkg7*), CD8^+^ NKT cells (*Cd3e, Klrb1c, Gzma, Nkg7, Cd8b1*), CD8^+^ T cells (*Cd3e , Cd8b1*), CD4^+^ T cells (*Cd3e, Cd4*), double-negative T (DNT) cells (*Cd3e*), double-positive T (DPT) cells (*Cd3e, Cd8b1, Cd4*), and B cells (*Cd19*) (Figs. 5F-H).

As an important control, given known ovotoxic effects of VCD and the absence of follicles 150 days post treatment (Figs. 1C,F and fig. S2C), we performed additional histological analyses on condition-matched ovaries (matched for age-at-injection and time post-injection) to confirm the presence of follicular structures in animals processed for scRNA-seq (fig. S4). H&E staining of condition-matched samples revealed that, despite extensive follicle depletion, residual follicles were still present in most VCD-treated animals at these time points, consistent with the presence of follicle-associated cell types in our scRNA-seq data (Figs. 5D,G). We also performed Picrosirius Red and Sudan Black B staining to quantify fibrosis and lipofuscin accumulation, respectively (fig. S5). Broadly, VCD-treated ovaries exhibited increased fibrosis and lipofuscin deposition relative to matched control samples at the 30- and 90-day time points (fig. S5), consistent with our previous observations at 150 days (Figs. 2C,F,I,L).

### RNAscope validation of ovarian cell type identities

To validate cell type assignments, we performed RNAscope, a high-sensitivity *in situ* hybridization assay for both well-characterized and less frequently profiled ovarian cell populations (Fig. 5I and figs. S6A-C). Granulosa cells (*Fshr*), theca cells (*Cyp11a1*), stromal cells (*Pdgfra*) and SMCs (*Acta2*) were validated using established marker probes, confirming their presence in the dataset (fig. S6A). Epithelial cells (*Upk1b*), BECs (*Flt1*) and LECs (*Prox1*) were similarly detected, supporting the robustness of annotations (fig. S6B). Importantly, we validated the presence of various immune cell subsets, including NK (*Klrb1c*), NKT (*Klrb1c*, *Cd3e*), CD8^+^ T (*Cd3e*, *Cd8b1*), CD4^+^ T cells (*Cd3e, Cd4*), DNT (*Cd3e,* with no Cd8b1/Cd4 signal) and B (*Cd19*) cells (fig. S6C). We also confirmed presence of less commonly reported ovarian cell types, including CD8+ NKT (*Klrb1c, Cd3e, Cd8b1*) and DPT (*Cd3e, Cd8b1, Cd4*) cells (Fig. 5I). These results further support the accuracy of the single-cell transcriptomic annotations and highlight the diverse somatic and immune microenvironments present across the aging models.

Together, a total of 19 cell types, 8 non-immune and 11 immune types, were shared across all three models (fig. S6D). Some cell types were not shared across models, potentially due to technical factors (e.g., capture efficiency and sequencing depth), or underlying biological variation inherent to each mouse model. To assist researchers in navigating the data efficiently, we developed an interactive R shiny app that makes the annotated datasets directly accessible and explorable through a graphical interface: https://minhooki.shinyapps.io/shinyappmulti/.

### scRNA-seq analysis reveals shifts in ovarian cell composition in menopause models

Next, we examined changes in cell type proportions across our datasets (Fig. 6A and table S6). We first assessed whether the proportion of immune cells shifted with ovarian aging by analyzing the expression of *Ptprc*, which encodes Cd45, a pan-immune marker (Figs. 6B-D). In the Aging model, we observed a significant increase in *Ptprc*^+^ cells, indicating an expansion of the immune cell population with age (Fig. 6B). This is consistent with previously published scRNA-seq studies in intactly aged mouse ovaries, which reported an increase in immune cell abundance at even younger ages (9-15.5 months) (*41, 49*).

**Fig 6.**
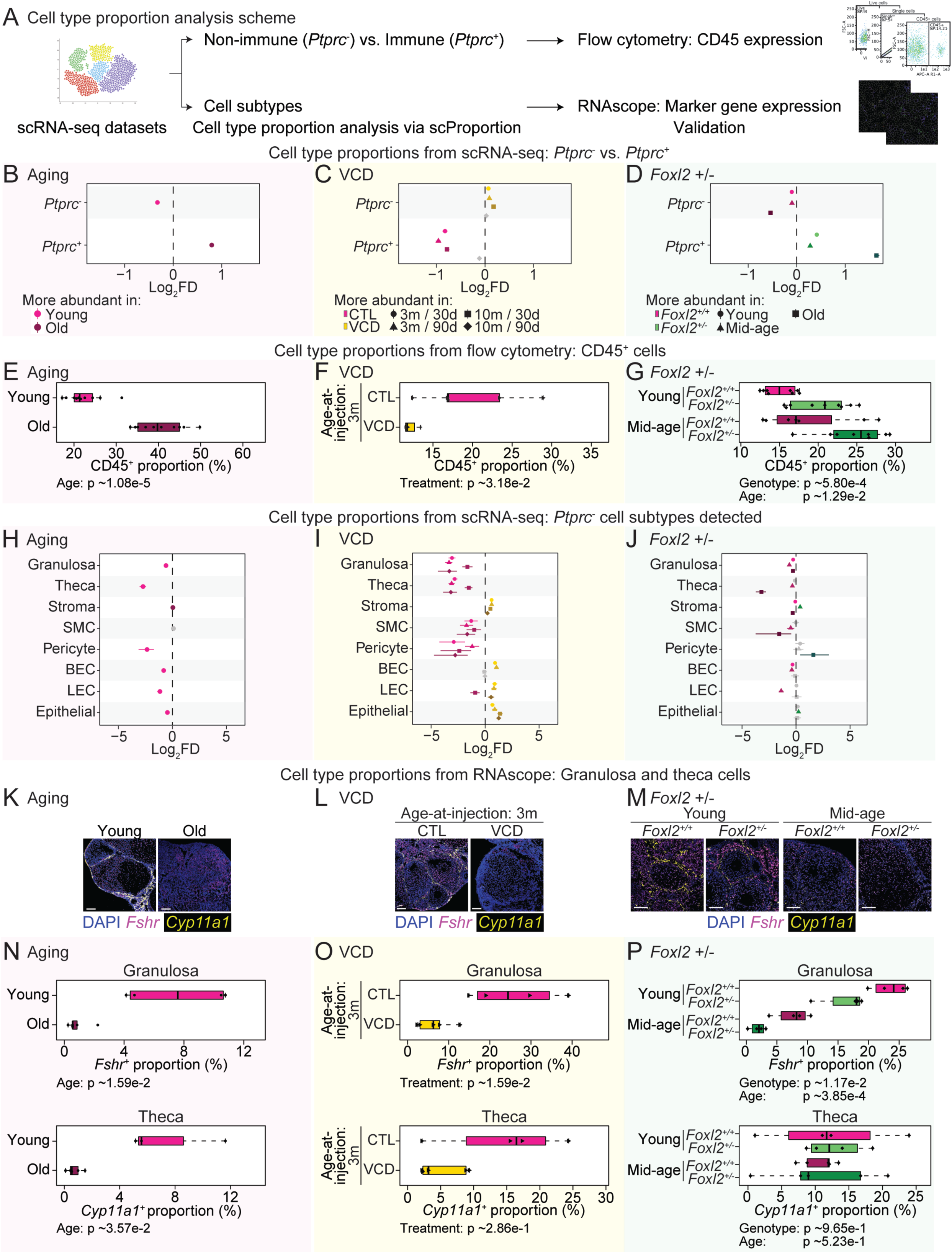
Characterization of cell type proportions in Aging, VCD, and *Foxl2* haploinsufficiency model scRNA-seq datasets. **(A)** Schematic of the experimental design. **(B-D)** Proportion differences of *Ptprc*^−^and *Ptprc*^+^ cells between young vs. old in Aging model, CTL (vehicle control) vs. VCD in VCD model, and *Foxl2^+/+^* vs. *Foxl2^+/−^*in the *Foxl2* haploinsufficiency model datasets. Colored points indicate statistically significant differences (p-value < 0.05) assessed using scProportionTest, whereas gray points denote non-significant comparisons. **(E-G)** Flow cytometry analysis of CD45^+^ cell proportions in ovaries of Aging, VCD, and *Foxl2* haploinsufficiency model mice. **(H-J)** Proportion differences of non-immune cell types detected in Aging, VCD, and *Foxl2* haploinsufficiency model scRNA-seq datasets. Color coding as in panels (**B-D)**. **(K-M)** Representative RNAscope images of *Fshr* (granulosa) and *Cyp11a1* (theca) probes from Aging, VCD, and *Foxl2* haploinsufficiency model mice. Scale bar: 100µm. **(N-P)** Cell abundance quantification data from RNAscope image analysis of *Fshr*^+^ and *Cyp11a1*^+^ cells from Aging, VCD, and *Foxl2* haploinsufficiency model mice. For panels **(E-G)** and **(N-P)**, statistical significance was assessed using Wilcoxon test for the Aging model and aligned rank transform ANOVA to test effects for the VCD and *Foxl2* haploinsufficiency models; p-values are shown.

VCD-treated mice exhibited a drastic reduction in *Ptprc*^+^ cells across all conditions, indicative of a depletion of immune cells in ovarian tissue (Fig. 6C). This reduction may result from the loss of follicular structures, alterations in immune recruitment, or uncharacterized systemic effects of VCD on immune homeostasis. Additionally, broader tissue remodeling, including fibrosis, may contribute to the altered immune cell landscape observed in VCD-treated ovaries. Thus, the VCD model does not recapitulate immune features of intact aging ovaries in mice, regardless of age-at-injection. In contrast, we detected a significant increase in *Ptprc*^+^ cells in *Foxl2^+/−^* animals, regardless of age (Fig. 6D). These findings were further validated by flow cytometric quantification of CD45^+^ cells (Figs. 6E-G and fig. S7A). Intriguingly, this suggests that the *Foxl2* haploinsufficiency model better recapitulates age-related shifts in ovarian immune cells compared to the VCD model.

Then, we conducted a comprehensive analysis of all ovarian cell populations detected in the datasets to identify shifts in cellular composition in response to intact aging, VCD exposure, and *Foxl2* haploinsufficiency (Figs. 6H-J and figs. S7B-D). In the Aging model, nearly all non-immune cell populations showed a decline in proportion with age, except for stromal cells and SMCs, which remained stable (Fig. 6H). In contrast, several immune cell populations increased with age, including neutrophils, ILCs, CD8^+^ NKT, CD4^+^ T, DNT, and DPT cells (fig. S7B). Notably, DNT cells have been reported to increase in aging mouse ovaries, and a potential role for these cells in regulating ovarian aging has been proposed (*41, 50*). Conversely, monocytes, NK, NKT, CD8^+^ T, and B cells exhibited a significant decrease with age (fig. S7B).

In the VCD model, cell proportion shifts were largely consistent within the treatment group, regardless of age-at-injection or time post-injection (Fig. 6I and fig. S7C). The VCD-treated group showed an increasing trend in stromal cells, BECs, LECs, epithelial cells, and CD4^+^ T cells. In contrast, granulosa cells, theca cells, SMC, and pericytes were significantly reduced in proportion (Fig. 6I). Among immune cells, neutrophils, monocytes, macrophages, DCs, NKT, DPT, and B cells were all decreased following VCD treatment (fig. S7C).

In the *Foxl2* haploinsufficiency model, granulosa cell proportions were consistently reduced in *Foxl2^+/−^* animals across all ages (Fig. 6J). In the *Foxl2* haploinsufficiency model, granulosa cell proportions were consistently reduced in *Foxl2^+/−^*animals across all ages (Fig. 6J). Notably, this reduction in granulosa cell representation was not accompanied by a significant decrease in total follicle number at the histological level (Fig. 1G and fig. S2D), suggesting that *Foxl2* haploinsufficiency may alter follicular composition (*e.g*. promoting more immature states, that hold lower granulosa cell number) or granulosa cell state (*e.g.* altered differentiation status, impaired maturation, or partial loss of canonical granulosa cell identity) rather than simply causing overt follicle loss. Given the established role of *FOXL2* in granulosa cell differentiation, maintenance (*51*), and AMH-associated regulation of follicle recruitment (*46*), one possible explanation is that partial *Foxl2* loss shifts follicles toward altered maturation states or changes granulosa cell transcriptional identity, thereby reducing their representation in scRNA-seq without necessarily reducing total follicle counts. Theca and SMCs also showed an overall decrease, whereas other non-immune populations showed relatively mild shifts (Fig. 6J). In contrast, immune cell populations generally increased in *Foxl2^+/−^* ovaries, although macrophages, dendritic cells, and DNT cells showed less consistent trends across ages (fig. S7D). Together, these results indicate reduced representation of key somatic ovarian cell types alongside a relative enrichment of immune populations.

To validate the cell abundance patterns identified by scRNA-seq, we performed RNAscope *in situ* hybridization for representative non-immune and immune populations (Figs. 6K-P and figs. S8, S9). We examined granulosa cells, theca cells, stromal cells, SMCs, BECs, LECs, and epithelial cells, along with NK, NKT, CD8⁺ NKT, CD8⁺ T, CD4⁺ T, DNT, DPT, and B cells. In the Aging model, RNAscope confirmed decreases in granulosa and theca cells with age (Figs. 6K,N). In old animals, residual granulosa signals were largely non-specific and did not colocalize with DAPI (Fig. 6K, right panel). BECs also showed a significant decline (fig. S8A). In contrast, stromal cells, SMCs, LECs, and epithelial cells did not show significant changes in abundance (fig. S8A). Thus, RNAscope validated several of the major somatic shifts detected by scRNA-seq while showing more limited changes for some populations. For immune populations, RNAscope largely recapitulated the scRNA-seq trends, showing decreases in NK, NKT, and B cells and increases in CD8⁺ NKT, CD4⁺ T, DNT, and DPT cells (figs. S7B, S9A). CD8⁺ T cells did not show a significant change by RNAscope despite a decrease with age in scRNA-seq, potentially reflecting differences in detection sensitivity or spatial distribution.

In the VCD model, we focused RNAscope analysis on the 3-month age-at-injection and 30-day post-injection condition, given consistent trends across groups (Figs. 6I,L,O and figs. S7C, S8B, S9B). RNAscope confirmed a reduction in granulosa cells and an increase in stromal cells following VCD exposure. Theca cells showed a decreasing trend but did not reach significance. Other non-immune populations did not show significant differences, in contrast to scRNA-seq findings, likely reflecting probe sensitivity or marker expression limitations. Among immune populations, RNAscope detected decreases in NK, NKT, CD8⁺ NKT, CD8⁺ T, and DNT cells, broadly consistent with the scRNA-seq results (fig. S9B). CD4⁺ T, DPT and B cells did not reach statistical significance.

For the *Foxl2* haploinsufficiency model, we focused RNAscope analysis on young and middle-age groups (Fig. 6J,M,P). Our analysis confirmed reduced granulosa cell proportions in *Foxl2^+/−^*animals across both age groups (Fig. 6J,M,P). Other somatic populations showed modest or variable changes. Immune populations showed patterns broadly consistent with the scRNA-seq analysis, including increased NK, NKT cells and CD8⁺ NKT cells (fig. S9C). Although some populations did not reach statistical significance, likely due to lower detection sensitivity, the overall trends supported immune expansion in *Foxl2^+/−^*ovaries. Together, these results validate the major cell composition shifts detected by scRNA-seq and support model-specific remodeling of ovarian somatic and immune populations.

### Global transcriptional perturbations of ovarian cell types across menopause models

To systematically evaluate how different menopause models influence global transcriptional responses in the ovary, we applied “Augur” (*52*), a computational framework that quantifies cell type-specific separability across experimental conditions based on single-cell level gene expression profiles (fig. S10A). Augur scores (derived from underlying machine-learning algorithm performance, reported as Area Under the Curve [AUC] values) represent the ability within each cell type to distinguish between biological groups within a given model based on overall transcriptional landscapes (*52*). In the Aging model, granulosa cells exhibited the highest AUC score (AUC ∼0.721) among non-immune cells, reinforcing their role as key transcriptional responders to chronological aging (figs. S10B,C). In the immune compartment, NKT and ILCs ranked highest (fig. S10C). Interestingly, DNT cells, despite their expansion in the aging ovary, showed the lowest AUC score (AUC ∼0.519), indicating limited transcriptional remodeling with age (fig. S10C).

In the VCD model, we observed marked transcriptional perturbations in theca, stromal, BEC, and LEC cells, especially at 90 days post-injection, with these effects largely consistent across both 3-month and 10-month age-at-injection groups (figs. S10D-G). However, epithelial and granulosa cells displayed more pronounced age-at-injection-dependent differences: epithelial cells were more responsive in younger animals, while granulosa cells were notably absent in the 10-month cohort, likely reflecting more severe follicular depletion (fig. S10G, left panel). Among immune cell types, NK, DNT, and CD8^+^ NKT cells showed higher transcriptional sensitivity in younger animals (fig. S10G, right panel). At 30 days post-injection, transcriptional divergence was generally more divergent in the VCD model compared to its 90-day counterpart (figs. S10D,E). For example, LECs in the 10-month group displayed strong divergence (AUC ∼0.81), suggesting age-related vascular remodeling may emerge early after VCD exposure.

In the *Foxl2* haploinsufficiency model, transcriptional divergence was more pronounced in old animals, suggesting that the effects of *Foxl2* loss may not fully manifest in the younger ovaries (fig. S11). Among non-immune populations, granulosa, theca and epithelial cells showed the highest AUC scores in older age groups (figs. S11C,E, left panels). In immune populations ILC, NK and monocytes exhibited the strongest divergence, with all showing elevated AUC scores in the older age groups, indicating potential immune activation or heightened sensitivity of specific immune compartments to *Foxl2* insufficiency (figs. S11C,E, right panels).

Together, these results reveal both common and model-specific overall transcriptional trajectories across ovarian aging paradigms. Granulosa cells consistently emerge as sensitive indicators of ovarian dysfunction across all three models. Immune cells show particularly pronounced divergence in *Foxl2^+/−^*animals, suggesting that immune alterations may precede, accompany, or even drive ovarian changes in this model. Notably, because *Foxl2* expression in adults is largely restricted to granulosa cells in the ovary and gonadotroph-lineage cells in the pituitary and it is not expressed at meaningful levels in ovarian immune cells (*33*), the immune-related phenotypes observed here are unlikely to be a cell autonomous results of decreased *Foxl2* expression within immune cell types. These comparisons underscore the importance of contextualizing cell type-specific transcriptional changes within each model’s mechanistic framework and highlight how distinct perturbations, such as chronological aging, chemical ablation, or genetic loss, can produce divergent molecular responses across ovarian cell types.

### Transcriptional and pathway-level signatures across menopause models

To investigate transcriptional remodeling across menopause models, we performed differential expression and pathway enrichment analyses across cell types and models (Fig. 7A and table S7). Importantly, we used a robust pseudobulking approach per annotated cell type in each individual scRNA-seq library, via “muscat” (*53*), so as to limit false positive rate. Six cell types, granulosa, theca, stromal, BEC, epithelial, and DNT cells, were consistently detected across all three models and thus included in downstream analyses (fig. S12A). “DESeq2” (*54*) was used to identify differentially expressed genes (DEGs) for each model: young vs. old in the Aging model; CTL vs. VCD in the VCD model (using age-at-injection and time post-injection as modeling covariates); and *Foxl2^+/+^* vs. *Foxl2^+/−^*in the *Foxl2* haploinsufficiency model (using age as a modeling covariate). As expected, we confirmed reduced *Foxl2* expression in *Foxl2^+/−^*animals in our scRNAseq dataset (fig. S12B).

**Fig 7.**
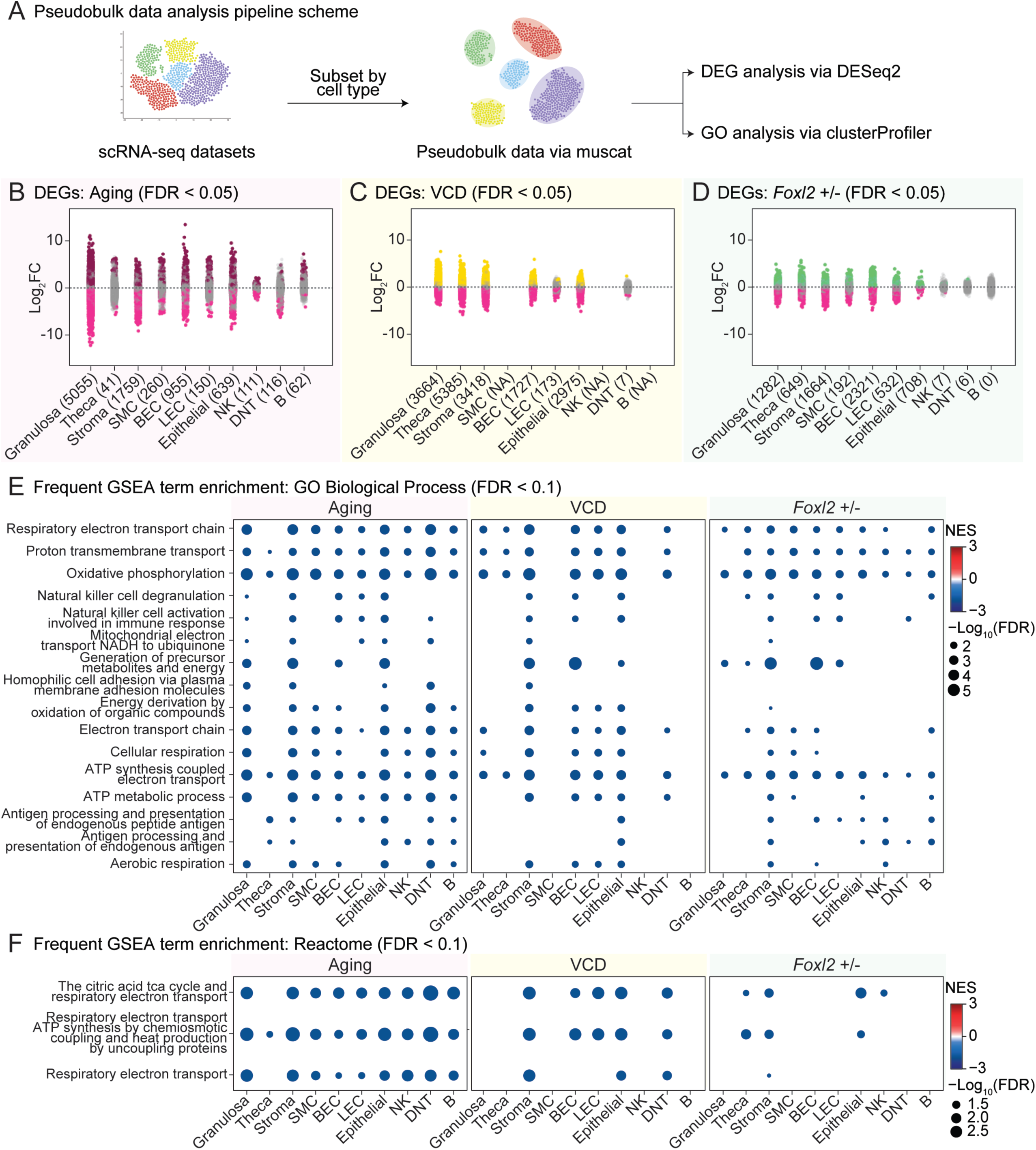
Pseudobulk analysis of differential gene expression and gene ontology analysis across Aging, VCD, and *Foxl2* haploinsufficiency models. **(A)** Schematic of pseudobulk analysis pipeline. **(B-D)** Strip plots of differentially expressed genes (DEGs) from Aging, VCD and *Foxl2* haploinsufficiency models. DEGs were identified using DESeq2, comparing young vs. old (Aging model), CTL vs. VCD (VCD model), and *Foxl2^+/+^* vs. *Foxl2^+/−^* (*Foxl2* haploinsufficiency model). Genes that passed the FDR < 0.05 threshold are colored and the numbers in parentheses following each cell type indicate the total number of DEGs that met the significance threshold. **(E-F)** Frequent Gene Set Enrichment Analysis (GSEA) term enrichment for GO Biological Process (GO BP, FDR < 0.1) and Reactome (FDR < 0.1) pathways across Aging, VCD, and *Foxl2* haploinsufficiency models.

Using an FDR cutoff of 0.05, we observed substantial transcriptional remodeling across models and cell types (Figs. 7B-D). Importantly, we asked such transcriptional changes may be unduly influenced by underlying differences in estrous cyclicity of mice. For this purpose, we took advantage of a publicly available scRNA-seq dataset profiling ovaries from young adult mice across the four stages of the estrous cycle (fig. S13A) (*55*). After processing the data using our standardized pseudobulk workflow, we focused on granulosa cells, which were among the most strongly perturbed populations across estrous cycle stages (*55*). We first compared multidimensional scaling (MDS) structures across the estrous-cycle dataset and our Aging, VCD, and *Foxl2* haploinsufficiency datasets (fig. S13B). Whereas our datasets showed sample separation according to the primary biological variables of interest (*i.e*. age, treatment, genotype), samples from the estrous-cycle dataset revealed substantial overlap in MDS space, suggesting smaller or less consistent transcriptional remodeling. Consistently, differential expression analysis identified only modest transcriptional differences between estrous stages, even in the proestrus-versus-estrus comparison, which has previously been reported to show the strongest stage-dependent effects (fig. S13C) (*55*). These findings indicate that the transcriptional changes observed across our menopause models are substantially larger than those attributable to estrous cycle-effects and are therefore unlikely to be driven primarily by stage differences.

Next, we performed gene set enrichment analysis (GSEA) to identify gene ontology (GO) biological process terms recurrently enriched across models (Fig. 7E and table S8). Mitochondrial-related gene sets were consistently downregulated across models and cell types, including “Respiratory electron transport chain,” “Oxidative phosphorylation” and “ATP synthesis coupled electron transport” (FDR < 0.1; Fig. 7F). Immune-related processes, such as “Antigen processing and presentation of endogenous peptide antigen,” were also frequently identified as downregulated. Using independent gene sets from Reactome (*56*) revealed consistent trends of mitochondria-related gene sets being downregulated (FDR < 0.1; Fig. 7F), suggesting convergent dysregulation of metabolic function during ovarian aging and in menopause models. Notably, immune-related gene sets were more prominently altered in the Aging and *Foxl2* haploinsufficiency models compared to the VCD model, consistent with the immune cell shifts observed in these datasets and suggesting that the *Foxl2* haploinsufficiency model could be particularly informative for investigating immune dysregulation associated with ovarian aging.

### Convergent and divergent aging-associated features are observed across menopause models

To identify transcriptional features shared or divergent across menopause models, we derived age-associated gene signatures from the Aging model and used them as reference cell type-specific “aging” gene sets (fig. S14A). The VCD model showed broad concordance, with consistent upregulation of aging-associated genes across cell types. In contrast, the *Foxl2* haploinsufficiency model displayed partial divergence. Granulosa and stromal cells showed reduced expression of genes typically upregulated with age, while BECs exhibited inverse trends for both age-associated up- and downregulated gene sets (fig. S14A). These cell type-specific deviations suggest that partial *Foxl2* loss may engage distinct molecular programs relative to physiological aging. We also evaluated enrichment of the published, validated SenMayo senescence-associated gene set (*57*) as a proxy for senescence burden (fig. S14B). As expected, most cell types in the Aging model showed increased SenMayo expression. The VCD model partially recapitulated this pattern, whereas the *Foxl2* haploinsufficiency model showed mixed, cell type-dependent effects, with several non-immune cell populations exhibiting inverse trends. These differences indicate that VCD and *Foxl2* haploinsufficiency models capture distinct cell-specific aspects of accelerated ovarian aging.

Increased transcriptional variability is a characteristic of aging and has been observed across multiple tissues, including the ovary (*49*) and hypothalamus (*58*) in mice, and is thought to reflect a loss of transcriptional regulation/precision with aging. Thus, we next evaluated gene expression variability in each cell type and condition using the coefficient of variation (CV) (fig. S14C). As expected, transcriptional variability broadly increased with aging across cell types (fig. S14C). The VCD model showed elevated variability primarily in theca cells, while other populations exhibited heterogeneous responses. In the *Foxl2* haploinsufficiency model, variability increased predominantly at middle-age, although inconsistently across cell types. These mixed patterns suggest that intervention models may involve compensatory or cell type-specific adaptations rather than uniform aging-like deregulation.

Finally, we examined transposable element (TE) transcription using “scTE” (*59*) (fig. S14D). TE transcription is known to be derepressed with aging across tissues, driven in part by age-associated chromatin remodeling, reduced heterochromatin integrity and impaired epigenetic repression (*60, 61*). While aging was associated with increased TE expression in several somatic and immune populations, intervention models showed more heterogeneous responses. VCD exposure led to increased TE expression in stromal and vascular compartments but reductions in steroidogenic cells, whereas *Foxl2* haploinsufficiency showed cell type-specific increases and decreases across ovarian lineages. These findings indicate that TE regulation varies depending on the mechanism of ovarian perturbation rather than uniformly mirroring chronological aging.

### Age-associated gene expression trajectories reveal model-specific amplification patterns

To better characterize how VCD exposure and *Foxl2* haploinsufficiency modulate age-associated transcriptional trajectories, we performed likelihood ratio test (LRT) analysis for the VCD and *Foxl2* haploinsufficiency models (Fig. 8A). Significant genes were subsequently grouped by unsupervised clustering to identify coherent gene expression patterns. Due to their relevance to ovarian function as key steroidogenic cells, we focused our analyses on granulosa and theca cells.

**Fig. 8.**
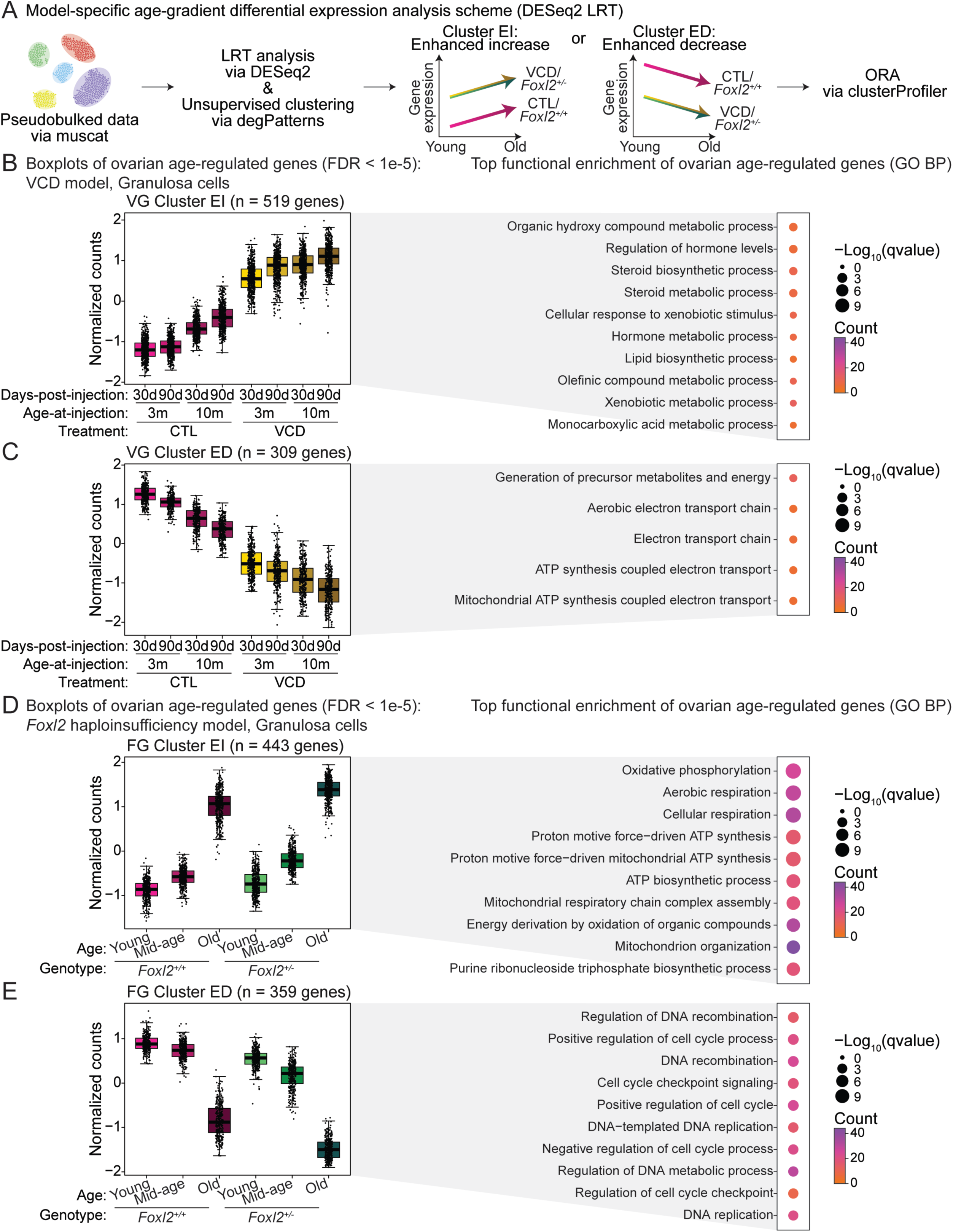
Age-associated gene expression trajectory analysis reveals model-specific amplification patterns in granulosa cells. **(A)** Schematic illustrating analytical workflow. Likelihood ratio tests (LRT) and unsupervised clustering of genes were performed using DESeq2 and degPatterns, respectively, to identify genes with significant expression changes across experimental groups in the VCD and *Foxl2* haploinsufficiency models. Clusters were designated as enhanced increase (EI) or enhanced decrease (ED) when perturbations amplified age-associated expression changes. Over-representation analysis (ORA) was performed for each cluster via clusterProfiler. **(B,C)** Boxplots showing normalized gene expression patterns of genes from granulosa EI and ED clusters in the VCD model. Bubble plots display ORA results for enriched pathways for each cluster. **(D,E)** Boxplots showing normalized gene expression patterns of representative genes from granulosa EI and ED clusters in the *Foxl2* haploinsufficiency model. Bubble plots show ORA enrichment results.

To interpret cluster behavior relative to physiological aging, we first examined expression patterns in corresponding control groups (CTL for the VCD model; *Foxl2^+/+^* for the *Foxl2* haploinsufficiency model) to establish baseline age-associated trajectories. Clusters were then classified based on how the perturbation modified these trajectories. When perturbations exaggerated age-associated increases or decreases, we termed these “enhanced increase” (EI) or “enhanced decrease” (ED) patterns, respectively (Fig. 8A).

In granulosa cells, both VCD exposure and *Foxl2* haploinsufficiency modified age-associated transcriptional trajectories in distinct, but partially overlapping ways (Figs. 8B-E). In the VCD model, EI genes were enriched for hormone metabolism pathways, whereas ED genes were associated with mitochondrial respiration (Figs. 8B,C). In the *Foxl2* haploinsufficiency model, EI genes were enriched for respiration-related pathways, while ED genes were linked to housekeeping processes, including cell cycle regulation, DNA replication, and recombination (Figs. 8D,E). Notably, mitochondrial and respiration-related pathways came up in both models, appearing in VCD ED clusters and *Foxl2* haploinsufficiency EI clusters, indicating that mitochondrial programs are broadly perturbed across models, albeit in opposite directions. Although additional significant gene regulatory patterns were also observed (fig. S15), these did not show consistent aging-associated trajectories and we did not perform further analyses on them.

In theca cells, we also observed broadly similar amplification patterns (figs. S16, S17). VCD-associated EI genes were enriched for metabolic pathways, whereas ED genes were enriched for processes linked to actin organization and cell signaling (figs. S16A-C). In the *Foxl2* haploinsufficiency model, ED genes were enriched for extracellular matrix and cell signaling pathways, suggesting altered stromal and intercellular communication (fig. S16D). Similar to the granulosa analysis, we also detected additional significant trajectory patterns in *Foxl2* haploinsufficient theca cells, but since these did not demonstrate clear aging-related behavior or significant functional enrichment, suggestive of model-specific effects (fig. S17), they were not explored further.

Together, these trajectory analyses reveal that both VCD exposure and *Foxl2* haploinsufficiency modulate age-associated transcriptional programs but do so in distinct ways. VCD primarily amplifies metabolic and mitochondrial changes in steroidogenic cells, whereas *Foxl2* haploinsufficiency preferentially affects pathways related to respiration, cell cycle regulation, and extracellular matrix organization. These results highlight how different menopause models reshape age-dependent transcriptional trajectories across ovarian cell types.

### Gene co-expression network and transcription factor activity across menopause models

To identify gene regulatory programs associated with ovarian aging, we performed weighted gene co-expression network analysis (WGCNA) (*62*) and transcription factor (TF) activity inference using “decoupleR” (*63*) (fig. S18A). As a reference framework, WGCNA was first applied to pseudobulked gene expression data from the Aging model, which captures physiological ovarian decline. Gene modules significantly correlated with age (FDR < 0.1) were retained for comparative analyses (figs. S18B, S19). As expected, these modules showed strong enrichment within the Aging dataset, providing a benchmark for assessing transcriptional convergence and divergence across intervention models (fig. S20A).

Module behavior in the VCD model largely paralleled that of the Aging model, indicating that VCD-induced ovarian failure recapitulates major regulatory programs associated with chronological aging (fig. S20A). In contrast, the *Foxl2* haploinsufficiency model displayed more divergent module dynamics. Notably, stromal cell modules frequently showed opposite expression patterns relative to the Aging model, suggesting that partial *Foxl2* loss alters regulatory programs governing stromal remodeling and tissue organization (fig. S20A). These comparisons highlight both shared aging-related regulatory circuits and model-specific deviations shaped by the underlying mechanism of ovarian dysfunction.

To investigate the functional basis of these modules, we performed Gene Ontology over-representation analysis (ORA; figs. S20B-D). In granulosa cells, modules were enriched for pathways related to mitochondrial metabolism and ribosomal function, consistent with high energetic and translational demands in steroidogenic cells (fig. S20B) (*64, 65*). Stromal cell modules were enriched for mitochondrial processes as well as mitotic machinery and chromosome segregation pathways, suggesting coordinated changes in cellular proliferation and tissue remodeling during aging (fig. S20C) (*41, 66*). In BECs, enriched terms included immune-related pathways, chromosome segregation, and cytoskeletal organization, pointing to coupled regulation of vascular integrity, immune interactions, and structural dynamics (fig. S20D) (*67*).

To complement co-expression network-based analyses, we also inferred transcription factor (TF) activity across cell types using decoupleR (fig. S21A and table S10). Across datasets, we detected shared changed TF activity signatures in multiple cell types (fig. S21B). Across datasets, shared TF activity changes were observed in multiple cell types (fig. S21B). In granulosa cells, we consistently detected reduced predicted activity of cell cycle-associated regulators (E2f1, E2f2, E2f3, E2f4, and Myc; fig. S21C) (*68, 69*). In contrast, predicted activity increased for steroidogenic regulators (Nr5a1, Nr5a2) (*70, 71*), metabolic regulators (Srebf1) (*72*), and stress- and hormone-responsive factors (Crem, Foxo1, Jun, and Sp1) (*73–76*), suggesting coordinated remodeling of endocrine function, metabolic homeostasis, and stress-adaptive programs in granulosa cells (fig. S21C).

Beyond granulosa cells, TF activity changes revealed distinct cell type-specific regulatory patterns (fig. S21C). Theca cells showed reduced activity of Ar, Foxo3, Nr3c1, and Stat3, consistent with attenuated steroidogenic signaling and stress-response pathways (*77, 78*). Stromal cells exhibited broad decreases in TFs associated with developmental signaling, tissue remodeling, and cellular plasticity, alongside increased immune function-linked regulatory activity, suggesting altered stromal homeostasis with enhanced immune involvement (*79–81*). Among vascular endothelial populations, BECs displayed reduced cell cycle-related TF activity (*82*), whereas LECs showed increased activity of immune and inflammatory regulators (*83, 84*), indicating divergent adaptations between blood and lymphatic vasculature. Epithelial cells exhibited reduced Sp1 activity, suggesting dampened transcriptional programs linked to cellular maintenance and metabolic regulation (*85*).

Collectively, these findings reveal both conserved and model-specific regulatory programs underlying ovarian aging, with coordinated transcription factor activity changes highlighting cell type-specific shifts in endocrine function, metabolic regulation, tissue remodeling, and immune adaptation.

### Transcriptome-based aging clocks capture evidence of accelerated ovarian aging across models

Finally, to obtain an integrative view of ovarian aging, we developed transcriptome-based clocks that predict ovarian age from gene expression profiles (Fig. 9A and fig. S22A), similar to approaches used in other tissues (*86, 87*). Given the central role of granulosa cells in ovarian function, we first trained a lasso regression model using granulosa cell transcriptomes (Fig. 9A). In parallel, a comparable model was constructed using theca cell transcriptomes to assess the fidelity of our menopause models in another key steroidogenic population (fig. S22A).

**Fig 9.**
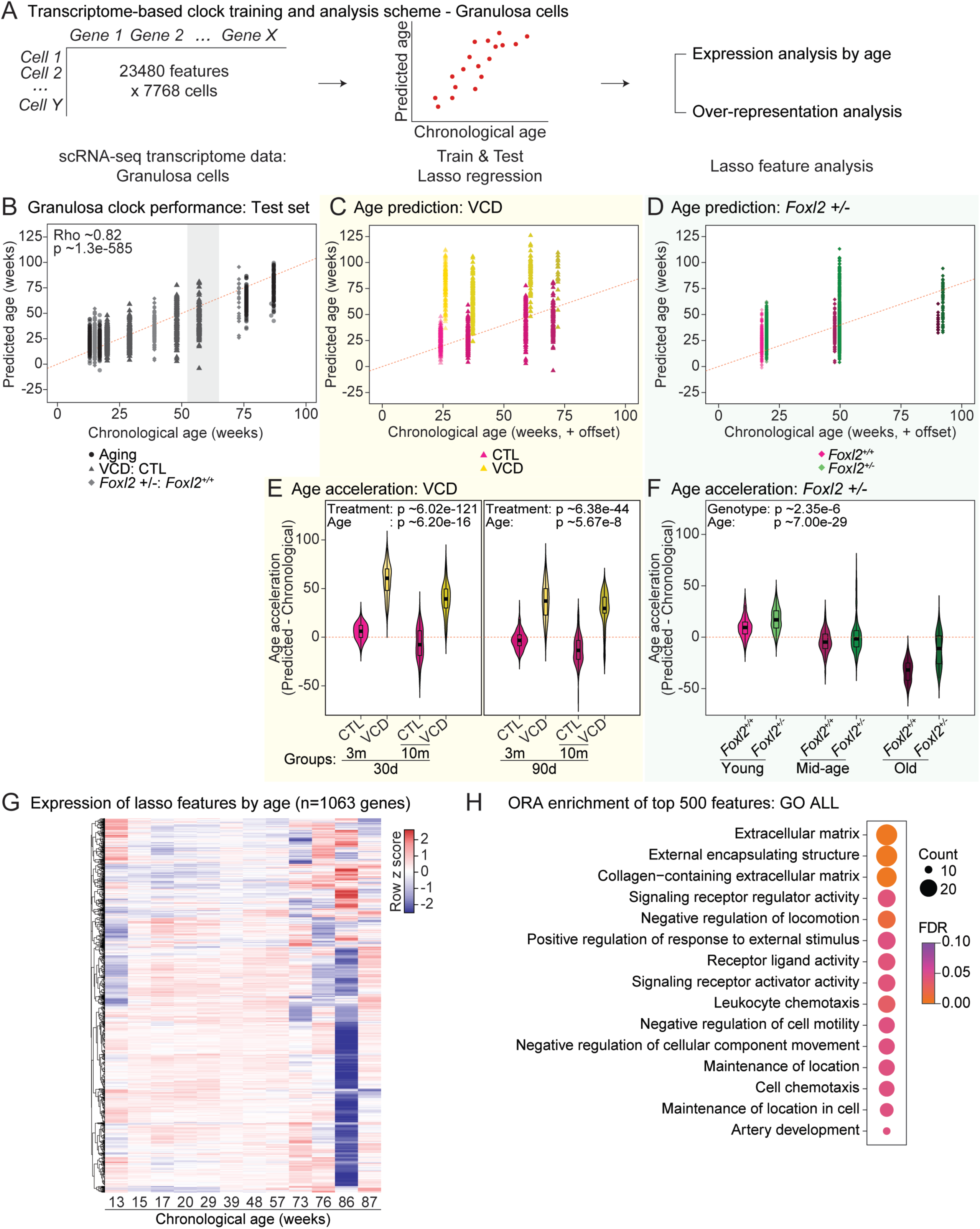
Development of a transcriptome-based aging clock for granulosa cells. **(A)** Schematic of analysis and training workflow for the lasso regression-based clock. **(B)** Scatter plot of predicted age *vs*. chronological age for the test set, with Spearman’s correlation and p-value displayed. **(C,D)** Age prediction for VCD and *Foxl2* haploinsufficiency datasets, comparing predicted age to chronological age in weeks. Offsets were added to improve visualization. **(E,F)** Age acceleration analysis for VCD and *Foxl2* haploinsufficiency datasets, calculated as the difference between predicted and chronological age. Statistical significance was assessed using aligned rank transform ANOVA to test effects. **(G)** Heatmap of expression lasso features from pseudobulked expression data. **(H)** Over-representation analysis of GO ALL terms of the top 500 lasso features.

Performance evaluation on held-out data demonstrated strong predictive accuracy for the granulosa-based model (Spearman’s Rho ∼0.82, p-value ∼1.3 × 10^−585^; Fig. 9B). The theca-based model also performed well (Spearman’s Rho ∼0.76, p-value ∼3.6 × 10^−447^; fig. S22B), although accuracy declined more sharply at older ages. Notably, prediction accuracy decreased in samples derived from mice aged ≥12 months, coinciding with the reported onset of estropause in C57BL/6 mice (*35*) (fig. S22B; expected age-at-estropause highlighted in gray).

We next applied the granulosa-based clock to VCD and *Foxl2* haploinsufficiency datasets (Figs. 9C,D). VCD-treated animals exhibited consistently elevated predicted transcriptomic ages relative to chronological age, indicating molecular acceleration of ovarian aging. *Foxl2^+/−^* animals similarly showed increased predicted ages compared to wild-type controls. To further quantify these differences, we next calculated transcriptomic age acceleration (predicted minus chronological age; Figs. 9E,F). In the VCD model, transcriptional age acceleration decreased with increasing age at exposure, mirroring trends observed in the hormone-based OvAge model, consistent with reduced responsiveness of older ovaries to chemical insult (Figs. 4F-H, 9E). In contrast, *Foxl2^+/−^* granulosa cells showed increased transcriptional age acceleration relative to controls, consistent with acceleration of ovarian aging and contrasting with our previous OvAge predictions (Figs. 4I, 9F). This discrepancy suggests that transcriptome-based clocks may capture earlier, distinct physiological alterations preceding systemic hormonal changes.

In parallel, we also applied the theca cell-based clock to the VCD and *Foxl2* haploinsufficiency models (figs. S22C,D). Despite its nominally lower accuracy, the theca clock recapitulated key aging patterns observed in granulosa cells. VCD-treated animals again showed elevated predicted transcriptomic ages. In the *Foxl2* haploinsufficiency model, genotype-dependent effects varied with age: *Foxl2^+/−^* animals exhibited increased predicted ages at young and middle-age but reduced age acceleration in the older cohort relative to wild-type animals. Age acceleration patterns were broadly consistent with results from granulosa-based clock in the VCD model, while *Foxl2* haploinsufficiency showed age-dependent genotype effects (figs. S22E,F). Analysis confirmed a significant overall genotype effect on age acceleration, supporting the sensitivity of the theca-based clock in detecting molecular ovarian aging across models. Together, these results demonstrate that transcriptome-based aging clocks capture molecular features of accelerated ovarian aging and reveal both convergent and cell type-specific responses to chemical and genetic perturbations.

Finally, to identify transcriptional features driving age prediction, genes with non-zero β coefficients from the trained lasso model were extracted and their expression trajectories examined across age in the pseudobulked granulosa cells (Fig. 9G). These genes displayed heterogeneous temporal patterns, indicating involvement in diverse age-associated regulatory programs. To further assess biological relevance, the top 500 genes based on absolute β coefficient were subjected to ORA (Fig. 9H). Enriched terms were predominantly related to extracellular matrix organization, cell signaling, and immune function, including “extracellular matrix,” “signaling receptor regulator activity,” and “leukocyte chemotaxis.” These results suggest that early alterations in tissue remodeling, intercellular communication, and immune regulation contribute substantially to ovarian transcriptomic aging and underscore the sensitivity of the transcriptome-based clock in detecting early molecular changes.

A similar approach was applied to the theca cell-based model (figs. S22G,H). Genes with non-zero β coefficients also exhibited heterogeneous age-dependent expression patterns, suggesting involvement in diverse regulatory programs during theca cell aging (fig. S22G). To evaluate their biological relevance, the top 500 genes based on absolute β coefficient were subjected to ORA (fig. S22H). Enriched terms were predominantly associated with reproductive and developmental processes, including “sex differentiation” and “gonad development.” Additional enrichment was observed for structural and translational processes such as “actin cytoskeleton reorganization” and components of the cytosolic ribosome, including “small ribosomal subunit.” These findings suggest that transcriptional aging in theca cells is characterized by coordinated remodeling of reproductive developmental programs alongside with shifts in cytoskeletal organization and protein synthesis capacity.

Together, these data demonstrate the utility of transcriptome-based models for detecting molecular aging trajectories in the mouse ovary. These predictive tools offer a valuable framework for assessing molecular age across experimental models and for uncovering early transcriptomic alterations that precede overt phenotypic decline.

## Discussion

Understanding how ovarian decline contributes to systemic aging requires robust experimental models that accurately recapitulate the physiological features of menopause. In this study, we systematically evaluated three mechanistically distinct mouse models of menopause: an intact aging model, a chemical follicle depletion model using VCD exposure, and a genetic model of *Foxl2* haploinsufficiency. By integrating histological, hormonal, and transcriptomic profiling across these models, we provide a comparative framework for understanding shared and model-specific trajectories of ovarian aging. Findings from our study provide a critical comparative resource for identifying conserved and divergent features of ovarian aging and evaluating the suitability of each model, with the goal of advancing both mechanistic insights and translational strategies in menopause research.

The intact Aging model showed expected hallmarks of physiological ovarian decline, including depletion of follicles across developmental stages, endocrine disruption, increased fibrosis and lipofuscin accumulation, and broad transcriptomic remodeling. At the cellular level, aging was associated with expansion of the immune compartment, reduction of most non-immune populations, and strong transcriptional perturbation in granulosa cells. These findings are consistent with prior scRNA-seq studies of mouse ovarian aging and reinforce the view that granulosa cells are among the most transcriptionally responsive somatic populations during ovarian decline (*41, 49*). Aging was also accompanied by increased transcriptional variability, altered TE expression, and coordinated pathway-level changes involving mitochondrial dysfunction and immune remodeling, indicating loss of tissue homeostasis across multiple biological scales.

The VCD model led to the appearance of several key features of accelerated ovarian failure. VCD-treated animals exhibited marked follicle depletion, endocrine disruption, increased fibrosis and lipofuscin accumulation, and elevated ovarian age as measured by both OvAge and transcriptome-based clocks. Importantly, by varying the age at VCD exposure, we demonstrate that ovarian failure can be induced across different stages of the reproductive lifespan, including near the estropausal transition. This establishes the VCD paradigm as a flexible system for modeling age-at-menopause, enabling the generation of ovarian failure within an age-relevant systemic context. Additionally, we observed that the magnitude of endocrine and transcriptomic acceleration was greater when VCD exposure occurred at younger ages and attenuated in animals treated closer to the estropausal transition. This finding indicates that the physiological consequences of follicle depletion are shaped by the baseline ovarian and systemic state at the time of insult. At the transcriptional level, VCD broadly recapitulated many features of the Aging model, including module behavior, age-associated gene signature enrichment, and transcriptome-based age acceleration. However, it also diverged from intact aging in important ways. Most notably, VCD-treated ovaries showed reduced immune cell proportions, in contrast to the immune expansion observed in naturally aged ovaries. This suggests that chemically induced follicle depletion captures aspects of ovarian failure and accelerated endocrine aging, but does not fully model the immune remodeling that accompanies physiological ovarian aging. Thus, the VCD model appears particularly useful for studying acute follicular depletion and endocrine collapse, but less suitable for interrogating immune-related features of intact ovarian aging.

The *Foxl2* haploinsufficiency model showed a more nuanced phenotype. Unlike the Aging and VCD models, *Foxl2^+/−^* animals did not exhibit overt reductions in combined follicle counts or consistent endocrine age acceleration by ovarian health index or OvAge. Nonetheless, this model displayed significant fibrosis and lipofuscin accumulation, impaired fertility, increased immune cell abundance, and marked transcriptional perturbation across several ovarian compartments. Granulosa cell proportions were consistently reduced, and transcriptome-based clocks detected elevated molecular age in granulosa cells despite the absence of a clear endocrine aging phenotype. Together, these findings indicate that *Foxl2* haploinsufficiency does not simply phenocopy overt menopause, but instead captures an alternative trajectory of ovarian dysfunction in which structural and molecular perturbations emerge before strong endocrine collapse. *FOXL2* interacts with *AMH* to regulate follicle recruitment in humans (*46*), raising the possibility that the observed histological, endocrine, and transcriptional features reflect enhanced AMH-mediated preservation of ovarian reserve in *Foxl2^+/−^* animals. Despite relatively preserved endocrine output, the heightened transcriptional sensitivity of steroidogenic and immune compartments positions this model as a valuable system for studying early or subclinical stages of ovarian aging.

This distinction is further supported by the transcriptomic analyses. Compared to the Aging and VCD models, *Foxl2* haploinsufficiency showed more divergent behavior in aging-associated gene signatures, senescence-associated programs, and co-expression modules, particularly in stromal cells. At the same time, immune-related signals were more prominent in *Foxl2^+/−^*ovaries, consistent with the increase in *Ptprc^+^* cells and the stronger transcriptional responsiveness of several immune populations. These findings suggest that partial loss of *Foxl2* may create an immune-primed ovarian environment and reshape transcriptional programs in ways that only partially overlap with chronological aging. Accordingly, the *Foxl2* haploinsufficiency model may be particularly informative for studying early or subclinical stages of ovarian dysfunction, especially those involving altered granulosa identity, tissue remodeling, and immune dysregulation.

We want to acknowledge that our *Foxl2* haploinsufficiency model uses a constitutive null allele rather than an ovary/granulosa-specific one (see Materials and Methods). However, this is not really a concern in the effort to model reproductive senescence due to the known adult expression pattern of *Foxl2*. Indeed, adult expression of *Foxl2* is largely restricted to ovarian granulosa cells and pituitary gonadotroph-lineage cells, with no evidence of expression in (ovarian) immune cell populations or other tissues (*33*). Thus, the immune phenotypes observed in *Foxl2^+/−^*ovaries are unlikely to arise from cell-autonomous *Foxl2* deficiency within immune cells, and instead more likely reflect secondary consequences of altered ovarian or neuroendocrine homeostasis. At the same time, because decreased *Foxl2* pituitary expression is likely to contribute to aspects of the reproductive phenotype (as they likely do in human *FOXL2* mutation carriers with premature ovarian failure), future studies could also evaluate how the hypothalamus/pituitary axis is impacted in the context of reproductive aging models, with relevance to human health. Notably, *FOXL2* has been shown to directly regulate expression of the common glycoprotein hormone α-subunit (Cga), a shared component of both FSH and LH, in pituitary gonadotrophs. Given that reproductive aging and menopause are characterized by elevated circulating FSH levels, and that this feature is recapitulated across our three models, partial *Foxl2* loss may contribute to dysregulation of gonadotropin production, thereby influencing ovarian function in a non-cell-autonomous manner (*88*).

Across models, several shared molecular themes emerged. Mitochondrial and respiration-related pathways were consistently altered at the pseudobulk level, and trajectory analyses in granulosa cells further highlighted mitochondrial programs as recurrent targets of remodeling in both VCD and *Foxl2* haploinsufficiency, albeit in opposite transcriptional directions. Gene regulatory network analyses also converged on processes related to mitochondrial metabolism, ribosomal function, cell cycle control, extracellular matrix organization, and immune adaptation. Transcriptome-based aging clocks reinforced these observations by identifying predictive features linked to extracellular matrix remodeling, signaling, immune regulation, and developmental programs. Together, these analyses point to common axes of ovarian aging biology while also showing that the direction, magnitude, and cellular context of these changes differ by model.

Our study also highlights the importance of cell type-specific readouts in evaluating ovarian aging. Granulosa cells consistently emerged as sensitive indicators across models, showing strong perturbation in Augur analyses, high DEG burden, and robust performance in transcriptome-based aging clocks. Theca cells similarly supported molecular age prediction and revealed complementary trajectory patterns, although the *Foxl2* model showed age-dependent divergence between granulosa- and theca-based clocks. These findings emphasize that ovarian aging is not uniform across cell types and that distinct compartments encode partially overlapping aspects of functional decline.

We would like to discuss several potential limitations of this work. First, this study was designed as a comparative resource and not to directly resolve how ovarian aging drives systemic aging phenotypes. Second, circulating estradiol was not quantified because technical constraints limit reliable measurement in mouse blood samples, including assay cross-reactivity with dietary phytoestrogens (including equol), extensive protein binding requiring complex extraction procedures, and limited sample volumes (*14*). Third, although the VCD age-at-injection paradigm enabled examination of age-dependent responsiveness to chemical follicle depletion, it does not directly model the aged systemic milieu. Finally, although cross-species parallels support the relevance of several pathways identified here, further work is needed to determine which model most faithfully captures specific aspects of human menopause biology.

In summary, our comparative analysis demonstrates that ovarian aging comprises both shared and model-specific structural, endocrine, and molecular features. The intact Aging model captures gradual physiological decline and immune remodeling; the VCD model induces robust follicle depletion and accelerated endocrine aging; and the *Foxl2* haploinsufficiency model reveals early molecular and tissue perturbations that may precede overt functional decline. By integrating histological, hormonal, and single-cell transcriptomic data, we provide a multidimensional framework for selecting and interpreting preclinical models of ovarian aging. This resource should support future studies aimed at identifying mechanisms, biomarkers, and interventions relevant to reproductive senescence and menopause-associated health decline.

## Materials and methods

### Experimental Design

This study was designed as a comparative resource to systematically evaluate mouse models of menopause and ovarian aging. Our objective was to determine how distinct mechanisms of ovarian dysfunction (physiological aging, chemically induced follicle depletion, and genetic disruption of granulosa cell identity), shape structural, endocrine, and molecular features of ovarian aging. To this end, we performed integrated histological, hormonal, and single-cell transcriptomic profiling across three models: intact aging, VCD-induced ovarian failure, and *Foxl2* haploinsufficiency.

Prespecified components included standardized histological assessment of follicle abundance and tissue remodeling, endocrine profiling of ovarian reserve markers, single-cell RNA sequencing to characterize cellular composition and transcriptional states, and integrative computational analyses to quantify molecular aging trajectories and regulatory programs. Harmonized preprocessing and analytical pipelines were applied across models to enable direct comparisons.

### Mouse husbandry

All animals were treated and housed in accordance with the Guide for Care and Use of Laboratory Animals. All experimental procedures were approved by the USC’s Institutional Animal Care and Use Committee (IACUC) and are in accordance with institutional and national guidelines. Samples were derived from animals on approved IACUC protocol numbers 21155 and 21454. All animals were acclimated in the specific-pathogen-free animal facility at USC for two weeks prior to any experimental procedures. Mice were provided PicoLab Rodent Diet 20 (LabDiet, 5053) *ad libitum*. The facility was maintained on a 12-hour light/dark cycle, with housing rooms set to 72°F and 30% humidity.

For the Aging model, female C57BL/6JNia mice (4- and 20-month-old) were obtained from the National Institute on Aging (NIA) colony at Charles River Laboratories.

For the VCD model, female C57BL/6J mice (2.5-, 5.5-, 7.5-, and 9.5-month-old) were purchased from Jackson Laboratory. Animals (strain # 000664) received daily intraperitoneal injections of either vehicle (safflower oil; Sigma S8281) or VCD (160 mg/kg/day; Sigma 94956) for 15 consecutive days. All injections were administered between 8:00 and 10:00 AM to minimize variability due to circadian influences.

*Foxl2* floxed (*Foxl2^lox/lox^*) mice were generated by Cyagen Biosciences Inc. using a previously described targeting strategy (*28*). Briefly, a targeting construct was designed to replace a 2.177 kb fragment containing the entire *Foxl2* coding region with a 1.157 kb *Neo* gene cassette (fig. S1A). The construct was linearized and introduced into C57BL/6NTac embryonic stem (ES) cells via electroporation. Candidate clones were assessed via PCR and Southern blot analysis. A targeted ES cell clone was selected and injected into C57BL/6NTac albino embryos, which were then implanted into pseudo-pregnant CD-1 females. Founder animals were identified based on coat color, and germline transmission was confirmed through breeding with C57BL/6J females followed by genotyping of the offspring. Mice carrying the desired floxed *Foxl2* allele were sent to USC for experiments and downstream phenotyping.

*Foxl2* haploinsufficiency mice (*Foxl2^+/−^*) were generated by crossing homozygous *Foxl2^lox/lox^* C57BL/6NTac mice with heterozygous B6.C-Tg(CMV-cre)1Cgn/J (JAX, stock #006054) mice. Offspring were genotyped for both the *Foxl2* alleles and the CMV-Cre transgene. To eliminate the CMV-Cre transgene, *Foxl2^+/−^* mice then were backcrossed to wild-type C57BL/6J mice. Progeny were selected to carry the deleted *Foxl2* allele (*Foxl2^+/−^*), but not the CMV-Cre transgene. Afterwards, *Foxl2^+/−^* mice were maintained as a stable heterozygous knock-out colony using allele transmission through parents of either sex with *Foxl2^+/+^ by Foxl2^+/−^* crossing to avoid generating full knockout animals. Routine genotyping was performed using genomic DNA extracted from tail biopsies using specific PCR primers, as listed in table S11A.

### RT-qPCR analysis of *Foxl2* expression in *Foxl2* haploinsufficiency model mouse ovaries

Flash-frozen ovaries collected from mice were used for RNA extraction, cDNA synthesis and quantitative RT-PCR. Ovaries were resuspended in 600 µL of TRIzol reagent (Thermo-Fisher, 15596018) and lysed using the BeadBug Homogenizer (Benchmark Scientific, D1036). Homogenization was performed at 3,500 rpm in 30-second intervals, repeated for a total of nine cycles. Total RNA was then purified using the Direct-Zol RNA Miniprep kit (Zymo Research, R2052), following the manufacturer’s protocol. RNA was eluted in nuclease-free water and immediately quantified using the NanoDrop spectrophotometer (Thermo Scientific). The purity of the RNA was assessed by measuring the 260/280nm and 260/230nm absorbance ratios. Samples with ratios close to ∼2.0 were considered suitable for downstream applications.

Total RNA was reverse transcribed into cDNA, using the Thermo Scientific™ Maxima H Minus First Strand cDNA Synthesis Kit (Thermo Scientific, K1682), following the manufacturer’s protocol. Quantitative PCR was performed using SensiFAST SYBR no-ROX kit (Bioline, BIO-98020) and the MIC Tubes and Caps kit (Bio Molecular Systems, 71e101) on the Magnetic Induction Cycler (MIC) machine (Bio Molecular Systems, MIC-2). micPCR v2.12.6 was used to capture and quantify Ct values. Gene expression levels were calculated using the ΔΔCt method, normalized to the geometric mean of two housekeeping genes: *Ubc* and *Hprt*. Primers and sequences used for RT-qPCR reactions can be found in table S11B.

### Fertility assessment of *Foxl2* haploinsufficiency model mice

Fertility data were obtained from our ongoing breeding colony, where *Foxl2^+/+^* and *Foxl2^+/−^*females were paired with *Foxl2^+/−^*and *Foxl2^+/+^* males of our colony, respectively. Importantly, data was included in the fertility assessment only if (i) *Foxl2^+/+^*and *Foxl2^+/−^* female pairings were initiated on the same day, to avoid confounds from potential housing-related or handling-related stress events, and (ii) females were aged 2.5 and 3.5 months at the time of pairing, to avoid biases related to hyperfertility of younger females and to allow sufficient time for *Foxl2* haploinsufficiency–associated phenotypes to emerge.

Fertility was assessed based on the pup count of the first litter and the latency to the first litter (*i.e*. the time between pairing and birth of the first litter). Differences in litter size between *Foxl2^+/+^* and *Foxl2^+/−^*females were assessed using the aligned rank transform ANOVA to test effects. One outlier from the *Foxl2^+/+^* group was removed based on a significant Grubbs’ test (p-value < 0.05), using the “outliers” package (v.0.15) (*89*) in R (v. 4.1.2). Latency to the first litter was analyzed using a log-rank test to compare survival curves, using the “survival” package (v. 3.6-4) (*90*) in R. Raw fertility data can be found in table S1A.

### Hematoxylin-eosin staining of mouse ovarian tissues

Ovaries were fixed in Bouin’s solution (Sigma, HT10132) for 24 hours at room temperature before being transferred to 70% ethanol for storage. Paraffin embedding, tissue sectioning, and hematoxylin and eosin (H&E) staining were carried out by the USC Norris Comprehensive Cancer Center Translational Pathology Core Facility and Immunohistochemistry Laboratory at USC Labs. H&E-stained slides were imaged using the Keyence BZ-X All-in-One Fluorescence Microscope platform using 20X objective and automated stitching using default parameters.

### Ovarian follicle counts

Follicle counts, including primordial, primary, secondary, and antral follicles, and corpus luteum, were performed on three sections per ovary by three blinded observers (fig. S2A). Median values from the three observers were used for data analysis. Statistical significance was assessed using the Wilcoxon test for the Aging model and aligned rank transform ANOVA to test effects for the VCD and *Foxl2* haploinsufficiency models.

### Ovarian fibrosis and lipofuscin histological staining and analysis

Ovaries were fixed in Bouin’s solution (Sigma, HT10132) for 24 h at room temperature and transferred to 70% ethanol for storage. Paraffin embedding, tissue sectioning, and hematoxylin and eosin (H&E) staining were performed by the USC Immunohistochemistry Laboratory. Histological staining was conducted using the following kits according to the manufacturers’ instructions: Picrosirius Red Stain Kit (Abcam, ab150681) for fibrosis detection and Sudan Black B Histochemical Stain Kit (StatLab, KTSBB) for lipofuscin detection. Stained slides were imaged using a Keyence BZ-X All-in-One Fluorescence Microscope with a 20x objective and automated stitching under default settings. For each ovary, three 8-µm sections were imaged and analyzed. Mean values across sections were used for downstream analysis.

Image quantification was performed using “ImageJ2” (v. 2.14.0/1.54f). Non-ovarian regions were manually traced and excluded from analysis. Color deconvolution was performed using the Colour Deconvolution 2 plugin with the FastRed FastBlue DAB vector. For Picrosirius Red images, the Colour_1 channel was used to quantify fibrotic signal and the Colour_2 channel to define total ovarian area. For Sudan Black B images, the Colour_2 channel was used to quantify lipofuscin signal and the Colour_1 channel to define total ovarian area. Deconvoluted grayscale images were thresholded to identify positive signal. Background thresholds (entire ovary) were set between 210–230, and positive signal thresholds were set between 150–170 depending on staining batch. Threshold parameters were determined separately for each batch based on positive and negative control sections. Signal intensity was calculated as the percentage of positive area relative to total ovarian area. All measurements were performed in a blinded manner.

Statistical significance was assessed using the non-parametric Wilcoxon rank-sum test for the Aging model. For the VCD and *Foxl2* haploinsufficiency models, factorial effects were evaluated using aligned rank transform ANOVA.

### Quantification of serum AMH, FSH and INHBA concentrations

Blood was collected either immediately following euthanasia via cardiac puncture or from live animals via a facial vein blood draw. Blood was allowed to clot at room temperature for one hour. Serum was then separated using the MiniCollect® Serum Tube (Greiner, 450472) and stored at -80°C until further analysis. The University of Virginia Center for Research in Reproduction Ligand Assay and Analysis Core quantified serum levels of AMH (Rat and Mouse Anti-Müllerian Hormone (AMH) ELISA kit, Ansh Labs, AL-113), FSH (Millipore Pituitary Panel Multiplex kit, RPT86K or Ultra-Sensitive Mouse & Rat FSH, UVA Ligand Core, in-house (*91*)), and INHBA (Inhibin A ELISA kit, Ansh Labs, AL-161), providing standardized normalized values. Statistical significance was assessed using the Wilcoxon test for the Aging model and aligned rank transform ANOVA to test effects for the VCD and *Foxl2* haploinsufficiency models. Raw serum hormone quantification data can be found in table S2.

Two different FSH assay kits were used due to changes in offered FSH quantitation assays at the UVA core during the time of our study. To ensure compatibility between measurements made using the different kits, we applied a correction procedure as previously described (*36*). Briefly, matched serum samples were analyzed using both kits, and a polynomial regression model was trained to capture the relationship between the two outputs. This model was then applied to adjust FSH values obtained using the ultra-sensitive assay, enabling direct computation of ovarian health index across datasets. The scripts used for ovarian health index calculation are available on the Benayoun lab GitHub repository: https://github.com/BenayounLaboratory/Ovarian_Aging_Microbiome/tree/main/1_Ovarian_health_index_calculation.

### Ovarian health index calculation

The ovarian health index was calculated as previously described (*36*). The scoring framework and reference thresholds were derived from the original cohort reported in that study. The index integrates two components: ovarian hormone levels (AMH, FSH, and INHBA) and follicle counts, which included the combined counts of primordial, primary, secondary, and antral follicles, as well as corpora lutea. Each parameter was assigned a score based on a three-tier system. Values exceeding the median of the young female group were assigned a score of 3, values positioned between the medians of the young and old groups were assigned a score of 2, and values below the median of the old group were assigned a score of 1. The hormone score was calculated as the mean of the individual scores for AMH, FSH, and INHBA. This hormone score was then combined with the follicle score in a 1:1 ratio to generate the overall ovarian health index, which was subsequently scaled to a 0-100 range for standardization. Statistical significance was assessed using the Wilcoxon test for the Aging model and aligned rank transform ANOVA to test effects for the VCD and *Foxl2* haploinsufficiency models.

### OvAge clock model training

The OvAge clock was trained using serum hormone quantification data from multiple model and cohorts, including animals from aging model, vehicle control animals from VCD model, wild-type animals from *Foxl2* haploinsufficiency model (*Foxl2*^+/+^) and wild-type animals from a previously published *Fshr* haploinsufficiency model (*Fshr*^+/+^) (*47*). To enhance the applicability of the model across different genetic and experimental backgrounds, animals from multiple models were included in the training dataset. Hormone data included serum levels of AMH, FSH, and INHBA, with corresponding age at blood collection recorded in weeks.

The dataset was partitioned into training (75%) and testing (25%) subsets using stratified sampling based on age, using the createDataPartition function from R package “caret” (v. 6.0-91) (*92*). A random forest (RF) regression model was trained using the training set, with out-of-bag (OOB) error correction applied for internal validation, using caret (v. 6.0-91) (*92*) and “randomForest” packages (v. 4.7-1) (*93*). The model was optimized by tuning the mtry parameter (best-performing value = 3), yielding an OOB root mean square error (RMSE) of 13.13 weeks and an OOB R² of 0.65. A total of 500 trees were used in the final model. Model performance was evaluated using Spearman correlation between chronological and predicted ages in the test dataset. The RF model achieved a Spearman correlation coefficient of 0.795 with a p-value of 5.5 × 10^−16^. Age acceleration was calculated as difference between predicted and chronological age. The final OvAge clock R object containing the trained RF model is available on the Benayoun lab GitHub repository: https://github.com/BenayounLaboratory/Mouse_Menopause_Models. We have also developed an interactive R shiny app to enable users to input hormone values and estimate predicted ovarian age using the OvAge clock: https://minhooki.shinyapps.io/OvAge_Predictor/.

### Single-cell RNAseq sample and library preparation

Ovaries from two animals (four ovaries total) were used for most single-cell RNA sequencing (scRNA-seq) sample and library preparations to optimize cellular viability and yield. In one instance (*Foxl2^+/+^ and Foxl^+/−^* old group pair), a single animal (two ovaries) was used due to sample availability.

Single-cell suspensions were generated using Miltenyi’s Multi Tissue Dissociation Kit (Miltenyi Biotec, 130-110-201) with the gentleMACS Octo Dissociator with Heaters (Miltenyi Biotec, 130-096-427), running the 37_m_LDK1 program. To maximize the recovery of viable single cells, the program was deliberately aborted 10 sec after start of the highspeed phase, approximately 20 sec before program completion. After dissociation, red blood cells were removed using Red Blood Cell Lysis Buffer (Miltenyi Biotec, 130-094-183) according to the manufacturer’s instructions. Dead cells were subsequently eliminated using the EasySep Dead Cell Removal (Annexin V) Kit (STEMCELL Technologies, 17899).

Cell count and viability were assessed via Viobility™ 405/452 Fixable Dye (Miltenyi Biotec, J66584-AB) staining and flow cytometry using a MACSQuant Analyzer 10 (Miltenyi Biotec, 130-130-420). Flow cytometry data were analyzed with FlowLogic v8 (Inivai Technologies). Cells were gated based on forward scatter area (FSC-A) vs. side scatter area (SSC-A), and singlets were identified by evaluating the linear grouping of cells in FSC-A vs. FSC-H plots (fig. S8A).

Single-cell libraries were prepared using the Chromium Next GEM Single Cell 3[ GEM, Library & Gel Bead Kit v3.1 (10X Genomics, PN-1000121) according to the manufacturer’s instructions. Based on flow cytometry estimates, cell suspensions were loaded to achieve a targeted recovery of 6,000 cells per sample. Samples were processed on a Chromium Next GEM Chip G (10X Genomics, 2000177) as per the manufacturer’s protocol.

Completed single-cell libraries were assessed for quality using the 4200 TapeStation system (Agilent Technologies, G2991A) with High Sensitivity D1000 DNA ScreenTape (Agilent Technologies, 50675584). Libraries were sequenced on an Illumina NovaSeq 6000, generating 150 bp paired-end reads at Novogene USA. Raw FASTQ reads have been deposited in the Sequence Read Archive under accession PRJNA863443.

### Single-cell RNAseq data analysis

#### Data processing

Raw sequencing reads were processed using “Cell Ranger” (v. 7.1.0) (*94*) from 10x Genomics (table S4). Reads were aligned to the mm10 reference genome. To remove contamination from ambient RNA, decontamination was performed using DecontX from the “celda” (v. 1.14.2) (*95*) package. Raw and filtered feature-barcode matrices were used to estimate ambient RNA levels, and corrected expression matrices were generated for each sample. Doublet removal was performed using a two-step approach. First, “DoubletFinder” (v. 2.0.3) (*96*) was applied to each sample independently, estimating doublet rates based on the expected multiplet rates from 10x Genomics. The optimal pK parameter was determined using a parameter sweep, and predicted doublets were labeled. In parallel, a second doublet detection method, “scds” (v. 1.10.0) (*97*), was applied using the cxds_bcds_hybrid function. Cells identified as doublets by either method were excluded from further analysis.

The filtered single-cell expression matrices were loaded into “Seurat” (v. 4.3.0) (*98*), and initial quality control (QC) steps were applied. Cells with fewer than 500 detected features, a total RNA count outside the range of 500-100,000, mitochondrial RNA content exceeding 15%, or a DecontX contamination score above 0.25 were removed. Following QC, datasets were normalized using SCTransform, regressing out the number of detected features, mitochondrial content, DecontX contamination score and batch effects.

#### Data integration

To correct for batch effects, integration was performed using “Harmony” (v. 1.0) (*99*). First, datasets were processed separately by normalizing gene expression using SCTransform, regressing out number of detected features (nFeature_RNA), mitochondrial percentage (percent.mito), ambient RNA contamination (decontX_contamination), and sequencing library (Library). Highly variable genes were selected using vst (nfeatures=2000 genes). The preprocessed datasets were merged using Seurat (v. 4.3.0) (*98*), ensuring shared variable features were maintained. Principal component analysis (PCA) was performed on the merged dataset, and batch effects were assessed by visualizing PCA embeddings colored by batch identity. Batch correction was applied using Harmony, where batch information was explicitly modeled as a covariate. The corrected embeddings were used for downstream analysis, including UMAP visualization, clustering, and differential expression analysis.

#### Cell type annotation

Cell type annotation was conducted using a multi-step approach that combined automated reference-based methods with manual marker-based annotation. First, *Ptprc*⁺ and *Ptprc*⁻ cell populations were identified using “scGate” (v. 1.0.1) (*100*). Cells expressing *Ptprc* (Cd45) were classified as immune cells, while those lacking *Ptprc* expression were designated as non-immune cells. The two subsets were then processed separately for further annotation. Each subset underwent independent cell type annotation using three complementary computational methods: “SingleR” (v. 1.8.1) (*101*), “scSorter” (v. 0.0.2) (*102*), and “scType” (v. 1.0) (*103*). The ImmGen dataset (*104*) was used as a reference for *Ptprc*⁺ cells, while two publicly available ovarian single-cell RNA-seq datasets (Open Science Framework ID 924fz (*55*) and NCBI GEO accession no. GSE232309 (*41*)) were used for *Ptprc*⁻ cells. To supplement these automated methods, manual annotation was conducted based on the expression of canonical marker genes specific to ovarian cell types (table S5). The final annotation was determined using a majority voting approach - cell type labels from SingleR, scSorter, scType, and manual annotation were compared, and the most frequently assigned label was selected as the final classification. All analyses were performed using Seurat (v. 4.3.0) (*98*) within R.

#### Cell type proportion and Augur analysis

Cell type proportions were quantified using “scProportionTest” (v. 0.0.0.9000) (*105*) within R. This method performs a permutation-based test to compare cell-type abundances between two groups, estimating the relative change in proportions along with a confidence interval for each cell type. The analysis was conducted at two levels: the broader classification of *Ptprc*⁺ vs. *Ptprc*⁻ populations and the more detailed sub-cell type level within each category. Relative proportion of detected cell types can be found in table S6.

We used “Augur” (v. 1.0.3) (*52*) to identify cell types that exhibited the most transcriptional changes associated with aging. Pairwise comparisons were conducted for each experimental condition, assessing differences between (1) young and old females, (2) vehicle control (CTL) and VCD-treated groups, and (3) *Foxl2^+/+^* and *Foxl2^+/−^* genotypes.

#### Pseudobulk analysis for differential gene expression

Pseudobulk differential gene expression analysis was conducted separately for three datasets: the Aging, VCD, and *Foxl2* haploinsufficiency models. Single-cell transcriptomic data were aggregated at the cell type level using “muscat” (v. 1.18.0) (*53*) to generate pseudobulk expression profiles. To ensure robust analysis, cell types were filtered based on dataset-specific criteria. In the Aging model, cell types with at least 25 cells per sample across all samples were retained. For the VCD model, cell types were included if they contained at least 25 cells in at least 12 samples, representing 75% of the dataset. In the *Foxl2* haploinsufficiency model, the threshold was set at a minimum of 25 cells in at least 8 samples, or 80% of the dataset.

Differential gene expression analysis was performed using “DESeq2” (v. 1.44.0) (*54*), with comparisons made between young and old females in the Aging model, vehicle control (CTL) and VCD-treated samples in the VCD model, and *Foxl2^+/+^* and *Foxl2^+/−^*samples in the *Foxl2* haploinsufficiency model. Batch effects in the VCD and *Foxl2* haploinsufficiency models were assessed using “sva” (v. 3.52.0) (*106*) to determine the number of surrogate variables (SVs), and batch correction was applied using the removeBatchEffect() function from “limma” (v. 3.60.6) (*107*). The adjusted counts were subsequently used for DESeq2 analysis, incorporating age-at-injection and time post-injection as covariates for the VCD model and age as a covariate for the *Foxl2* haploinsufficiency model. Variance-stabilized transformed counts were computed using getVarianceStabilizedData(), and differentially expressed genes (FDR < 0.05) were visualized using strip plots. The complete pseudobulk DESeq2 output, including log fold changes, adjusted p-values, and base mean expression values for each dataset and cell type, can be found in table S7.

#### GO analysis for differential gene expression

Gene set enrichment analysis (GSEA) was performed to identify Gene Ontology Biological Process (GO BP) and Reactome terms enriched in differentially expressed genes across cell types in the Aging, VCD, and *Foxl2* haploinsufficiency models. The analysis was conducted using the “clusterProfiler” package (v. 4.2.2) (*108*) and the Molecular Signatures Database (MSigDB) gene sets retrieved via “msigdbr” (v. 7.4.1) (*109*). Differential expression results from pseudobulk DESeq2 analysis were ranked by t-statistic and used as input for GSEA. Enrichment analysis was performed separately for each cell type in each dataset, with an FDR cutoff of 0.1 applied to identify significant terms. To identify recurrent biological processes across datasets, GO-BP terms enriched in at least four cell types within each dataset were extracted. The list of significantly enriched GO-BP terms (FDR < 0.1) for all datasets can be found in table S8.

#### Estrous cycle transcriptomic variability analysis

To evaluate potential confounding effects of estrous stage, we reanalyzed a publicly available mouse ovary scRNA-seq dataset spanning the four estrous stages (*55*). Count matrices were obtained from Open Science Framework (accession 924fz), and stage annotations were incorporated as metadata. Granulosa cells were aggregated into pseudobulk profiles using muscat (v. 1.18.0) (*53*) and analyzed using DESeq2 (v. 1.44.0) (*54*) following the same pipeline described above. Variance-stabilized counts were used for principal component analysis (PCA; prcomp) and multidimensional scaling (MDS; Spearman distance, cmdscale). Differential expression analysis was performed in DESeq2, focusing on the proestrus versus estrus contrast.

#### GSEA of aging-associated transcriptional signatures across models

GSEA was performed using differentially expressed genes (DEGs) from the Aging model dataset as input gene sets with the clusterProfiler (v4.12.6) (*108*) package. Upregulated and downregulated gene sets (FDR < 0.05) were extracted from the Aging model dataset and tested for enrichment in the Aging, VCD and *Foxl2* haploinsufficiency model datasets. Genes were ranked based on the DESeq2-derived t-statistic for analysis. Additionally, GSEA was conducted to assess the enrichment of the SenMayo (*57*) gene set within the dataset. Differential expression results from the Aging, VCD, and *Foxl2* haploinsufficiency datasets were used for GSEA, with genes ranked using the DESeq2-derived t-statistic.

#### Coefficient of variation analysis

Gene expression variability across cell types was assessed by calculating the coefficient of variation (CV) for each gene in the Aging, VCD, and *Foxl2* haploinsufficiency models. CV was computed as the ratio of the standard deviation to the mean expression level per gene within each condition (young vs. old for Aging, CTL vs. VCD for VCD, and *Foxl2^+/+^* vs. *Foxl2^+/−^* for *Foxl2* haploinsufficiency model) using variance-stabilized transformed counts obtained from DESeq2 (v1.44.0) (*54*) pseudobulk analysis. For the *Foxl2* haploinsufficiency model, the old-age group was excluded from CV analysis due to insufficient biological replicates. Statistical differences in CV between groups were evaluated using the Wilcoxon rank-sum test.

#### Transposable element (TE) analysis

Transposable element (TE) expression was quantified using “scTE” (v1.0) (*59*) with default parameters. BAM files generated from Cell Ranger (v. 7.1.0) (*94*) outputs were processed using the mm10 genome index provided by scTE. Features detected by scTE but absent in the single-cell gene expression data were extracted and merged, generating a Seurat object containing both gene and TE expression profiles.

Cells were aggregated at the cell type level using muscat (v1.18.0) (*53*), and differential expression analysis was performed using DESeq2 (v1.44.0) (*54*), following the same approach as described above.

To assess functional enrichment of TE families, GSEA was conducted on ranked gene lists derived from DESeq2 results. TE annotations were obtained from the UCSC database for mm10, the same reference used by scTE. TE subfamilies were stratified to broader categories, including LINE, SINE, LTR, and DNA transposons. TE family annotations were obtained from the *rmsk.txt* file downloaded from the UCSC genome annotation database for the mm10 reference genome. For each cell type, genes were ranked by t-statistic from differential expression analysis, and enrichment of TE families was assessed using clusterProfiler (v4.12.6) (*108*). Significant TE families (FDR < 0.1) were reported for visualization.

#### Likelihood ratio test–based trajectory analysis

Pseudobulk counts of granulosa and theca cells from each dataset were analyzed using likelihood ratio tests (LRTs) implemented in DESeq2 (v1.44.0) (*54*) to identify genes exhibiting perturbation-dependent age-associated transcriptional trajectories in the VCD and *Foxl2* haploinsufficiency models. Resulting p-values were adjusted using the Benjamini-Hochberg method, and genes with FDR < × 10^−5^ were retained. Variance-stabilized expression values of significant genes were subjected to unsupervised clustering using the degPatterns function from “DEGreport” (v 1.40.1) (*110*). Baseline age-associated trends were defined from control groups, and clusters were classified according to whether perturbations amplified or attenuated age-related transcriptional changes.

#### WGCNA and ORA enrichment analysis

Weighted gene co-expression network analysis (WGCNA) (*62*) was performed on the Aging model dataset to identify gene co-expression modules. Variance-stabilized transformed counts from DESeq2 (v1.44.0) (*54*) were used as input, with analyses conducted separately for each cell type. For each cell type, an optimal soft-thresholding power was determined using pickSoftThreshold, followed by the construction of an adjacency matrix and topological overlap matrix (TOM). Hierarchical clustering was performed to identify gene modules, which were assigned colors using cutreeDynamic, and eigengenes were computed to summarize module expression patterns. Gene sets from each module were classified into upregulated and downregulated groups based on eigengene expression differences between young and old samples. These gene sets were then subjected to GSEA via clusterProfiler (v. 4.2.2) (*108*), using differential expression results from the VCD and *Foxl2* haploinsufficiency model datasets to assess whether the identified modules were significantly enriched in these datasets. Modules significantly associated with aging (FDR < 0.1) were further analyzed through over-representation analysis using clusterProfiler (v. 4.2.2) (*108*) and the org.Mm.eg.db annotation package.

#### Transcription factor activity inference using decoupleR

Transcription factor (TF) activity inference was performed using “decoupleR” (v2.10.0) (*63*) with the CollecTRI transcriptional regulatory network for mouse. Differential expression results from the Aging, VCD, and *Foxl2* haploinsufficiency models were used as input. For each dataset, fgsea scoring was applied to rank TF activity based on DESeq2-derived t-statistics, and significant TFs were identified at a threshold of FDR < 0.05. To compare TF activity across datasets, TFs were filtered to retain those with consistent activity direction (positive or negative) in at least two datasets.

#### R shiny application generation

An interactive R shiny application of Aging, VCD and *Foxl2* haploinsufficiency scRNA-seq datasets was generated using “ShinyCell” (v.2.1.0) (*111*), and made available at https://minhooki.shinyapps.io/shinyappmulti/.

### RNAscope sample preparation, imaging and data analysis

#### *In situ* Hybridization Protocol

Frozen ovaries were mounted in Tissue-Tek O.C.T Compound (Sakura, 4583) and sliced into 8 μm slices from the same ovarian region using PTFE-coated microtome blades (Duraedge, 7223) at −20°C on a Cryostat CM1860 (Leica, 14-0491-46884). 3 ovarian sections from the same region were mounted in succession on VWR Premium Superfrost Plus Microscope slides (VWR, 48311-703). All batches of sample processing were prepared using the fresh-frozen sample preparation protocol along with negative and positive controls (Advanced Cell Diagnostics, UM 323100-USM). 12 probes were tested in 3 sets, with Set 1 containing markers for theca (*Cyp11a1*; C1), granulosa (*Fshr*; C2), smooth muscle (*Acta2*; C3), and stromal (*Pdgfra*; C4) cells. Set 2 had markers for epithelial cells (*Upk1b*; C1), lymphatic endothelial cells (*Prox1*; C2), blood endothelial cells (*Flt1*; C3), and B cells (*Cd19*; C4). Set 3 contained probes for T (*Cd3e*; C1), natural killer (*Klr1b1c*; C2), CD8+ (*Cd8b1*; C3), and CD4+ (*Cd4*; C4) cells. Per the protocol, the RNAscope^TM^ Multiplex Fluorescent Reagent Kit v2 (Advanced Cell Diagnostics, 323100) was used. The fluorescent signals TSA Vivid Fluorophore 520 (Advanced Cell Diagnostics, 323271), TSA Vivid Fluorophore 570 (Advanced Cell Diagnostics, 323272), TSA Vivid Fluorophore 650 (Advanced Cell Diagnostics, 323273), and Opal 690 (Akoya Biosciences, FP1497001KT) were used to stain the probes in each set. The slides were preserved in ProLong Gold Antifade Mountant (Thermo Fisher Scientific, P36930).

#### Imaging

Imaging was performed using a Leica Stellaris 5 confocal microscope (Leica Microsystems) equipped with a HC PL APO 20x/0.75 IMM CORR CS2 objective (Leica, 11506343) and Leica Microsystems Type F Immersion liquid oil (Leica Microsystems, 11513859). Images were acquired using Leica Application Suite X software (LAS X v4.4.0).

#### Imaging analysis

To quantify probe expression, the images were analyzed using the “QuPath” program (v0.5.1). Cell boundaries based on DAPI signal were calculated using the following settings modified from the default parameters – Requested pixel size: 0 μm, Nucleus background radius: 10 μm, Maximum area: 400 μm^2^, Threshold: 0.5, and Cell expansion: 2. Sets 1 and 2 were analyzed using subcellular detection to determine their expression levels relative to the number of cells. All thresholding values were uniform across the date of processing of each ovarian tissue. For Set 3, single measurement classifiers were created for each probe to generate colocalization data. Cells below the detection threshold were not counted for analysis.

### Cell-type proportion analysis using flow cytometry

Single-cell suspensions of ovarian cells were prepared following the same protocol used for single-cell RNA sequencing sample preparation, as described above. Cells were stained for APC-CD45 antibody (Miltenyi, 130-123-784) according to the manufacturer’s recommendations. Flow cytometry data were acquired using the MACSQuant10 flow cytometer (Miltenyi Biotec, 130-096-343) and analyzed using FlowLogic v.8 (Inivai Technologies). The raw flow cytometry dataset was deposited to Figshare (DOI: 10.6084/m9.figshare.28733222).

### Granulosa and theca transcriptional clock model training and feature assessment

Granulosa and theca cells were independently subsetted from scRNA-seq datasets encompassing multiple experimental models, including natural aging, VCD, and *Foxl2* haploinsufficiency. Additionally, an independent middle-aged female dataset (13, 76, and 86 weeks) was included to ensure age coverage for intermediate ages not well represented in the datasets.

To develop the cell transcriptional clocks, cells from aging model, middle-aged females, vehicle controls from the VCD model, and *Foxl2+/+* animals from the *Foxl2* haploinsufficiency model were extracted from the dataset. These cells were used to train and evaluate the model. Data were partitioned into training (75%) and testing (25%) sets using stratified sampling, using the createDataPartition function from the “caret” package (v. 6.0-91). Training data were used for model optimization, while the test set served as an independent validation dataset.

Predictive models were trained using L1-regularized lasso regression with the “glmnet” package (v. 4.1-7) to estimate transcriptional age (in weeks) based on gene expression profiles. Five-fold nested cross-validation was performed on the training dataset, and the optimal regularization parameter (lambda.min) was determined based on cross-validation results. Spearman correlation analysis was performed to assess the correlation between predicted and chronological age.

Transcriptional clocks were applied to experimental test groups, including VCD-treated and *Foxl2^+/−^* animals. Model predictions for test groups were compared to those of control samples to assess differences in predicted age patterns. Additionally, age acceleration was calculated as the difference between predicted and chronological age. Statistical significance of group differences in predicted age and age acceleration was assessed using the Wilcoxon rank-sum test.

To examine the age-associated expression dynamics of lasso features selected by the models, we performed pseudobulk analysis using control samples. For each cell type, raw counts were aggregated per sample using muscat (v. 1.18.0) (*53*), and DESeq2 (v. 1.44.0) (*54*) was used to compute variance-stabilized transformed expression values. Batch effects were corrected using removeBatchEffect, and expression heatmaps of lasso-selected features were generated to visualize age-dependent expression trends.

The top 500 lasso features were selected based on the absolute values of their coefficients. Over-representation analysis was performed using clusterProfiler (v. 4.2.2) and “org.Mm.eg.db” (v. 3.14.0), analyzing “ALL” Gene Ontology terms. A p-value cutoff of 0.05 was applied to identify significantly enriched pathways.

### Statistical Analysis

All statistical analysis was performed using the R software, version 4.1.2. For all boxplots, the data is shown with the median, the 25th and 75th percentile of the data, and the whiskers represent 1.5 ∗ the inter-quartile range (IQR). Individual datapoints are overlayed, when possible, for transparency and rigor. Specific statistical tests, number of biological replicates and animals used are indicated in the corresponding figure legends and associated methods.

## Acknowledgements

We thank Dr. Gilbert Garcia for guidance and feedback on confocal imaging. We are grateful to Dr. Lynae M. Brayboy for providing the ovarian single-cell dissociation protocol and for valuable advice on protocol optimization. We thank Kristen Mehalko for maintaining the Foxl2 haploinsufficiency mouse colony and assisting with tissue collection. We acknowledge Younggyun Kim, Kelly Koh, and Sanjana Paye for assistance with follicle quantification from hematoxylin and eosin-stained ovarian sections. We thank Dr. Victor Ansere, Clayton Baker, and Aaron J.J. Lemus for constructive feedback on the manuscript.

Ovarian histological processing was performed by the Translational Pathology Core at the USC Norris Comprehensive Cancer Center and Immunohistochemistry Laboratory at USC Labs, and serum hormone measurements were conducted by the University of Virginia Center for Research in Reproduction Ligand Assay and Analysis Core. Some schematic panels were generated using the Generic Diagramming Platform.

## Funding

Global Consortium for Reproductive Longevity and Equality Postdoctoral Fellowship GCRLE-2020

supported by the Bia-Echo Foundation (MK)

USC Provost’s Undergraduate Research Fellowship (JW)

Global Consortium for Reproductive Longevity and Equality Junior Scholar Award GCRLE-0520

supported by the Bia-Echo Foundation (BAB)

Pew Biomedical Scholar Award #00034120 from The Pew Charitable Trusts (BAB)

Generous gifts from Kathleen Gilmore and Dr. Eric Hennigan (BAB)

National Cancer Institute Cancer Center Support Grant P30 CA014089 supporting the Translational Pathology Core at the USC Norris Comprehensive Cancer Center

Eunice Kennedy Shriver National Institute of Child Health and Human Development grant R24HD102061

supporting the University of Virginia Center for Research in Reproduction Ligand Assay and Analysis Core

## Author contribution

Conceptualization: MK, BAB

Investigation: MK, RB, JW, XL, MC, RJL, EHL, JLA, RGW, BAB

Formal analysis: MK, RB, JW, EHL, BAB

Visualization: MK, JW, BAB

Supervision: BAB

Writing - original draft: MK, BAB

Writing - review & editing: MK, RB, JW, XL, RJL, EHL, JLA, RGW, BAB

## Competing interest

Authors declare that they have no competing interests.

## Data availability

The raw FASTQ files have been deposited in the Sequence Read Archive under accession number PRJNA863443. Raw microscopy images of ovarian hematoxylin and eosin (H&E), Picrosirius Red and Sudan Black B staining, compressed RNAscope images (due to file size constraints), and raw flow cytometry datasets are available on Figshare (DOIs: 10.6084/m9.figshare.28745765, 10.6084/m9.figshare.28745792, 10.6084/m9.figshare.28745813, 10.6084/m9.figshare.29260391 and 10.6084/m9.figshare.28733222). Any additional information required to reproduce or reanalyze the data presented in this study is available from the corresponding author upon reasonable request.

All new scripts used to analyze the datasets are available on the Benayoun lab GitHub at https://github.com/BenayounLaboratory/Mouse_Menopause_Models.

## List of Supplementary Materials

Figs. S1 to S22

Tables S1 to S11

## Supplementary figures

**Fig. S1. Characterization of the *Foxl2* haploinsufficiency model. (A)** Schematic representation of constructs for *Foxl2* haploinsufficiency mouse line generation. **(B)** *Foxl2* expression levels in *Foxl2*^+/+^ and *Foxl2*^+/−^ mice ovaries from young and mid-age groups, measured by RT-qPCR (n = 5 per group). P-values were calculated using aligned rank transform ANOVA to test effects of genotype and age. **(C,D)** Fertility assessment of *Foxl2* haploinsufficiency females, comparing first litter pup counts and latency to first pregnancy (n = 11 and 15 for *Foxl2^+/+^* and *Foxl2^+/−^*, respectively). P-values were calculated using the Wilcoxon and log-rank tests, respectively.

**Fig. S2. Follicle counts across Aging, VCD and *Foxl2* haploinsufficiency models. (A)** Follicle counting scheme. Pr: primordial follicle; P: primary follicle; S: secondary follicle; A: antral follicle; CL: corpus luteum. **(B-D)** Follicle counts for primordial, primary, secondary and antral follicles, and corpus luteum, from Aging (n = 25 per group), VCD (n = 5-10 per group), and *Foxl2* haploinsufficiency (n = 11-24 per group) model mice. For panels **(B-D)**, statistical significance was assessed using Wilcoxon test for the Aging model and aligned rank transform ANOVA to test effects for the VCD and *Foxl2* haploinsufficiency models; p-values are shown.

**Fig. S3. Harmonization of FSH results for the *Foxl2* haploinsufficiency model and longitudinal analysis of serum AMH, FSH, and Inhibin A levels in the VCD model. (A)** Polynomial regression-based correction model used to harmonize FSH measurements obtained from two different assay kits (n = 6 and 9 for *Foxl2^+/+^* and *Foxl2^+/−^*, respectively). **(B-D)** Serum levels of AMH, FSH, and INHBA measured from 0 to 5 months post-injection in VCD model mice (n = 5 per group).

**Fig. S4. Histological characterization of the VCD model mice. (A)** Representative hematoxylin and eosin staining images of ovarian tissues from condition-matched animals used in the VCD model scRNA-seq analysis. Scale bar: 250µm. **(B,C)** Follicle counts for the condition-matched animals used in the VCD model scRNA-seq analysis (n = 4-5 per group). Statistical significance was assessed using aligned rank transform ANOVA; p-values are shown.

**Fig. S5. Condition-matched analysis of ovarian fibrosis and lipofuscin accumulation following VCD exposure. (A,B)** Representative Picrosirius Red staining images of ovarian tissues collected at 30 days and 90 days post-VCD injection. **(C,D)** Quantification of percent fibrotic area at 30 and 90 days post-injection (n = 5 per group). **(E,F)** Representative Sudan Black B staining images showing lipofuscin accumulation in ovarian tissues at 30 and 90 days post-injection. **(G,H)** Quantification of percent lipofuscin-positive area at 30 and 90 days post-injection (n = 4-5 per group). Scale bar: 200 µm. For **(C,D,G,H)**, statistical significance was assessed using aligned rank transform ANOVA; p-values are shown.

**Fig. S6. Validation of detected ovarian cell types by *in situ* hybridization. (A)** Representative RNAscope images for *Fshr, Cyp11a1, Pdgfra* and *Acta2* probes. **(B)** Representative RNAscope images for *Upk1b, Cd19, Flt1* and *Prox1* probes. **(C)** Representative RNAscope images for *Klrb1c, Cd3e, Cd8b2* and *Cd4*. Images were enhanced for visualization. **(D)** Heatmap of detected cell types in the Aging, VCD and *Foxl2* haploinsufficiency scRNA-seq datasets. For panels **(A-D)**, shown images are from a middle-aged *Foxl2^+/+^* animal.

**Fig. S7. Cell type proportion analysis of ovarian cells via flow cytometry and scRNA-seq analysis. (A)** Gating strategy of CD45^+^ cells in flow cytometry analysis. **(B-D)** Proportion differences of *Ptprc*+ cell types detected in Aging, VCD and *Foxl2* haploinsufficiency model scRNA-seq datasets. Colored points indicate statistically significant differences (p-value < 0.05) assessed using scProportionTest, whereas gray points denote non-significant comparisons. Asterisks (*) indicate comparisons in which the observed log_2_ fold change is infinite due to complete absence of the cell type in one of the groups.

**Fig. S8. Cell type proportion analysis of non-immune ovarian cells via *in situ* hybridization assays. (A-C)** Cell type abundance analysis of stromal, SMC, BEC, LEC and epithelial cells via RNAscope *in situ* hybridization assays from Aging, VCD and *Foxl2* haploinsufficiency model mice (n = 4-5 per group across models). Statistical significance was assessed using Wilcoxon test for the Aging and VCD models, and aligned rank transform ANOVA for the *Foxl2* haploinsufficiency model; p-values are shown.

**Fig. S9. Cell type proportion analysis of immune ovarian cells via *in situ* hybridization assays. (A-C)** Cell type abundance analysis of NK, NKT, CD8^+^ NKT, CD8^+^ T, CD4^+^ T, DNT, DPT and B cells via RNAscope *in situ* hybridization assays from Aging, VCD and *Foxl2* haploinsufficiency model mice (n = 4-5 per group across models). Statistical significance was assessed using Wilcoxon test for the Aging and VCD models, and aligned rank transform ANOVA for the *Foxl2* haploinsufficiency model; p-values are shown.

**Fig. S10. Comparative analysis of global gene expression in the Aging and VCD models via Augur. (A)** Schematic of global gene expression analysis comparisons using Augur. **(B,C)** UMAP visualization of AUC scores and lollipop plot of AUC quantification from the Aging model. **(D-G)** UMAP visualization of AUC scores and scatter plot of AUC quantification from the VCD model, comparing CTL vs. VCD at 3m and 10m age-at-injection and 30 days vs. 90 days post-injection. For the scatter plot, Ptprc- and Ptprc+ cells were separately plotted to improve visualization. For **(E,G)**, data points with NA AUC scores (due to low cell count or QC filtering) were assigned an AUC of 0.5 and colored gray to improve visualization.

**Fig. S11. Comparative analysis of global gene expression in the *Foxl2* haploinsufficiency model via Augur. (A-E)** UMAP visualization of AUC scores and scatter plot of AUC quantification from the *Foxl2* haploinsufficiency model comparing *Foxl2^+/+^* vs. *Foxl2^+/−^* at young, mid-age and old. For the scatter plot, *Ptprc^−^* and *Ptprc^+^*cells were separately plotted to improve visualization. For **(C,E)**, data points with NA AUC scores (due to low cell count or QC filtering) were assigned an AUC of 0.5 and colored gray to improve visualization.

**Fig. S12. Cell type representation across pseudobulked scRNA-seq datasets and validation of Foxl2 expression in Foxl2 haploinsufficiency mice. (A)** Heatmap of cell types detected in pseudobulked scRNA-seq datasets. **(B)** DESeq2 normalized log_2_ counts of *Foxl2* from young, mid-age and old *Foxl2^+/+^* and *Foxl2^+/−^*mice.

**Fig. S13. Assessment of estrous cycle effects on ovarian transcriptional profiles. (A)** Overview of a publicly available single-cell RNA-seq dataset profiling mouse ovaries across the four estrous cycle stages. Granulosa cell transcriptomes were pseudobulked for downstream analyses. **(B)** Multidimensional scaling plots of pseudobulked granulosa cell transcriptomes from Cycling, Aging, VCD and *Foxl2* haploinsufficiency model data. **(C)** Differential expression analysis (FDR < 0.05) between estrous cycle dataset (estrus vs. proestrus) and menopause model dataset.

**Fig. S14. Aging-associated signatures and transposable element enrichment analysis across Aging, VCD and *Foxl2* haploinsufficiency models. (A,B)** GSEA enrichment analysis of differentially expressed genes dentified in the Aging model (FDR < 0.05) and SenMayo gene list. **(C)** Dot plot showing the coefficient of variation analysis results. **(D)** GSEA enrichment analysis of transposable element (TE) families across Aging, VCD and *Foxl2* haploinsufficiency models.

**Fig. S15. Classification of age-associated transcriptional trajectory patterns in granulosa cells. (A)** Boxplot showing normalized gene expression patterns of genes from granulosa cluster 1 in the VCD model (VG Cluster 1). Bubble plot displays ORA results for enriched pathways for the cluster. **(C)** Boxplot showing normalized gene expression patterns of genes from granulosa cluster 2 in the *Foxl2* haploinsufficiency model (FG Cluster 2). Bubble plot shows ORA enrichment results.

**Fig. S16. Age-associated transcriptional trajectory patterns and pathway enrichment in theca cells. (A-C)** Boxplots showing normalized gene expression patterns of genes from theca EI, ED 1 and ED 2 clusters in the VCD model. Bubble plots display ORA results for enriched pathways for each cluster. **(D)** Boxplot showing normalized gene expression patterns of genes from theca ED cluster in the *Foxl2* haploinsufficiency model. Bubble plot shows ORA enrichment results.

**Fig. S17. Age-associated transcriptional trajectories in *Foxl2* haploinsufficient theca cells. (A,B)** Boxplots showing normalized gene expression patterns of genes from theca clusters 1 and 2 in the *Foxl2* haploinsufficiency model. No enrichment was identified via ORA.

**Fig. S18. Module trees of WGCNA analysis. (A)** Schematic of Weighted Gene Co-expression Network Analysis (WGCNA) and transcription factor inference analysis. **(B)** WGCNA module trees identified in the Aging model for granulosa, theca, stromal, BEC, LEC, epithelial, and DNT cells.

**Fig. S19. Association analysis of WGCNA modules and ovarian aging traits from the Aging model. (A)** Heatmap of trait significance for WGCNA modules in the Aging model. Modules that passed the FDR < 0.1 threshold are shown in bold.

**Fig. S20. Preservation of co-expression modules in Aging, VCD, and *Foxl2* haploinsufficiency models. (A)** GSEA enrichment of WGCNA module genes associated with aging traits across Aging, VCD and *Foxl2* haploinsufficiency models. **(B-D)** Bubble plots of over-representation analysis (ORA) results of GO ALL terms for WGCNA module genes (FDR < 0.1).

**Fig. S21. Analysis of recurrent transcription factor activity inferred by decoupleR. (A)** Heatmap of transcription factor (TF) activity for factors detected in at least three cell types within each model with an FDR < 0.05. (B) (C) **(B)** UpSet plot of decoupleR-inferred TF activity, showing the number of TF overlap across datasets and the corresponding cell types. **(C)** Heatmap of TF activity scores of common decoupleR-inferred TFs for each cell type.

**Fig. S22. Development of a transcriptome-based aging clock for theca cells**. **(A)** Schematic of analysis and training workflow for the lasso regression-based clock. **(B)** Scatter plot of predicted age *vs*. chronological age for the test set, with Spearman’s correlation and p-value displayed. **(C,D)** Age prediction for VCD and *Foxl2* haploinsufficiency datasets, comparing predicted age to chronological age in weeks. Offsets were added for visualization purposes. **(E,F)** Age acceleration analysis for VCD and *Foxl2* haploinsufficiency datasets, calculated as the difference between predicted and chronological age. Statistical significance was assessed using aligned rank transform ANOVA to test effects. **(G)** Heatmap of expression lasso features from pseudobulked expression data. **(H)** Over-representation analysis of GO ALL terms of the top 500 lasso features.

## Supplementary tables

**Supplementary table 1. Raw fertility data from *Foxl2* haploinsufficiency model mice.**

**Supplementary table 2. Raw serum AMH, FSH and INHBA quantification data.** Raw data from **(A)** Aging, **(B)** VCD and **(C)** *Foxl2* haploinsufficiency models.

**Supplementary table 3. BH-adjusted post hoc p-values for OvAge age acceleration in VCD and Foxl2 models.**

**Supplementary table 4. 10xGenomics CellRanger QC metrix.**

**Supplementary table 5. List of canonical marker genes used to annotate cell types in scRNA-seq datasets.**

**Supplementary table 6. Relative proportion of detected cell types from scRNA-seq datasets.** Proportion data from **(A)** Aging, **(B)** VCD and **(C)** *Foxl2* haploinsufficiency models.

**Supplementary table 7. DESeq2 output from pseudobulk analysis.** DESeq2 output from **(A)** Aging, **(B)** VCD and **(C)** *Foxl2* haploinsufficiency model datasets.

**Supplementary table 8. Significant GSEA enrichment terms (FDR < 10%) for GO Biological Process and Reactome terms.** GSEA enrichment terms for Biological Process from **(A)** Aging, **(B)** VCD and **(C)** *Foxl2* haploinsufficiency model datasets. GSEA enrichment terms for Reactome from **(D)** Aging, **(E)** VCD and **(F)** *Foxl2* haploinsufficiency model datasets.

**Supplementary table 9. LRT DESeq2 output with cluster information.** LRT DESeq2 output with degPatterns cluster information from **(A)** VCD model granulosa cells, **(B)** *Foxl2* haploinsufficiency model granulosa cells, **(C)** VCD model theca cells and **(D)** *Foxl2* haploinsufficiency model theca cells.

**Supplementary table 10. List of decoupleR TF activity scores (FDR < 5%) for each cell type. Supplementary table 11. Primers used in the study. a**, Primers used for genotyping *Foxl2* haploinsufficiency model animals. **b**, Primers used for *Foxl2* expression quantification in *Foxl2* haploinsufficiency model animals via RT-qPCR.

## Notes

### Competing Interest Statement

The authors have declared no competing interest.

### Summary of Updates

- Added additional samples to scRNAseq datasets - Added histological stains for fibrosis (PSR) and lipofucin (Sudan Black) analysis - Added deeper analyses of granulosa and theca cell transcriptomes - Textual updates for clarity and precision

