## Supplementary material for "Systematic characterization of the ovarian landscape across mouse menopause models"

**Supplementary table 10. List of decoupleR TF activity scores (FDR < 5%) for each cell type.**

**Supplementary table 11. Primers used in the study.** **a**, Primers used for genotyping *Foxl2* haploinsufficiency model animals. **b**, Primers used for *Foxl2* expression quantification in *Foxl2* haploinsufficiency model animals via RT-qPCR.

Fig. S1

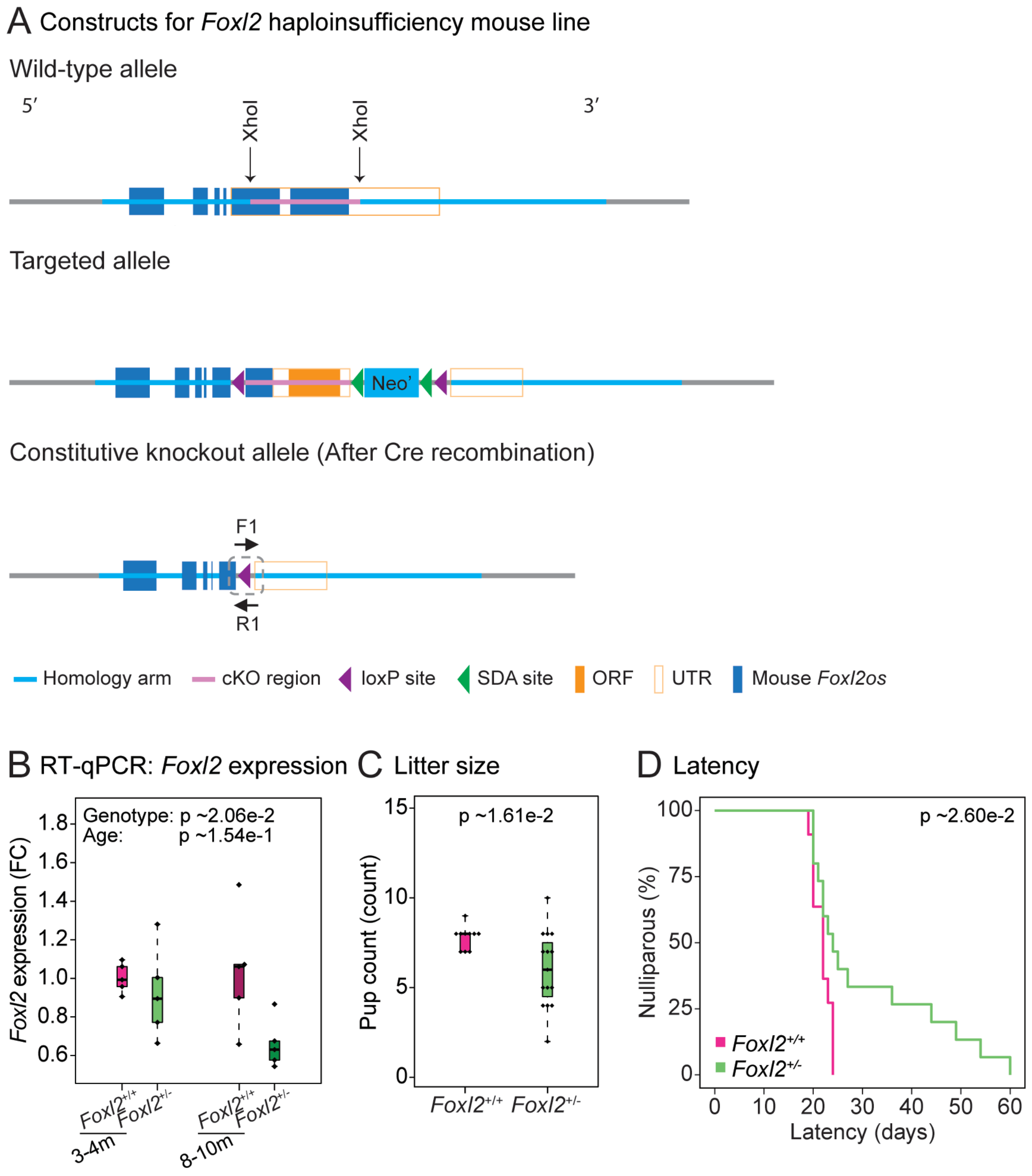
A Constructs for Foxl2 haploinsufficiency mouse line
Wild-type allele
5′
3′
XhoI
XhoI
Targeted allele
Neo'
Constitutive knockout allele (After Cre recombination)
F1
R1
Homology arm
cKO region
loxP site
SDA site
ORF
UTR
Mouse Foxl2os
B RT-qPCR: Foxl2 expression
Genotype: p ~2.06e-2
Age: p ~1.54e-1
Foxl2 expression (FC)
Foxl2+/+
Foxl2+/-
3-4m
8-10m
C Litter size
p ~1.61e-2
Pup count (count)
D Latency
p ~2.60e-2
Nulliparous (%)
Latency (days)

Fig. S2

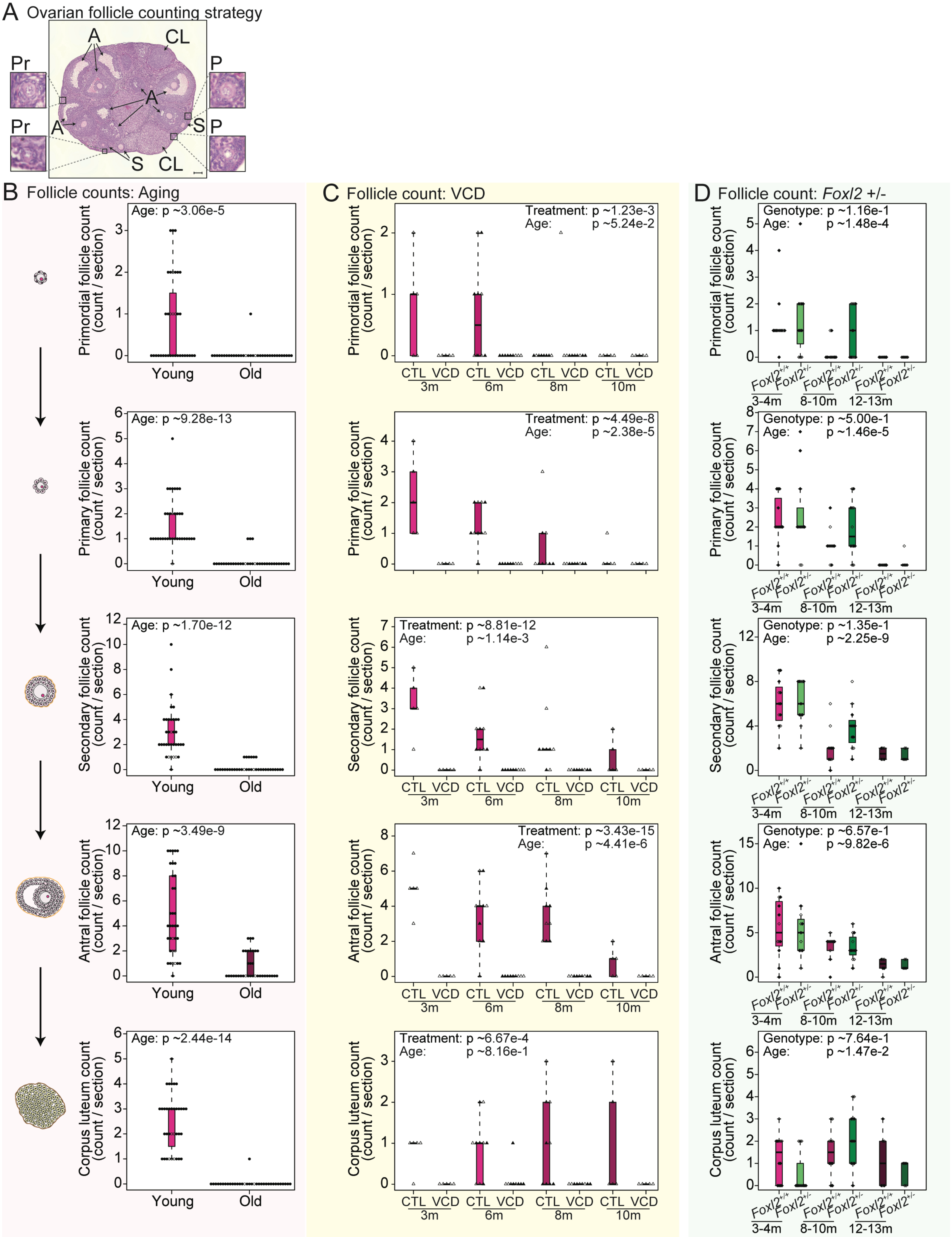
A Ovarian follicle counting strategy
A
CL
Pr
P
S
B Follicle counts: Aging
Age: p ~3.06e-5
Primordial follicle count (count / section)
Age: p ~9.28e-13
Primary follicle count (count / section)
Age: p ~1.70e-12
Secondary follicle count (count / section)
Age: p ~3.49e-9
Antral follicle count (count / section)
Age: p ~2.44e-14
Corpus luteum count (count / section)
Young
Old
C Follicle count: VCD
Treatment: p ~1.23e-3
Age: p ~5.24e-2
Treatment: p ~4.49e-8
Age: p ~2.38e-5
Treatment: p ~8.81e-12
Age: p ~1.14e-3
Treatment: p ~3.43e-15
Age: p ~4.41e-6
Treatment: p ~6.67e-4
Age: p ~8.16e-1
CTL VCD CTL VCD CTL VCD CTL VCD
3m 6m 8m 10m
D Follicle count: Foxl2 +/-
Genotype: p ~1.16e-1
Age: p ~1.48e-4
Genotype: p ~5.00e-1
Age: p ~1.46e-5
Genotype: p ~1.35e-1
Age: p ~2.25e-9
Genotype: p ~6.57e-1
Age: p ~9.82e-6
Genotype: p ~7.64e-1
Age: p ~1.47e-2
Foxl2+/+ Foxl2+/- Foxl2+/+ Foxl2+/- Foxl2+/+ Foxl2+/-
3-4m 8-10m 12-13m

Fig. S3

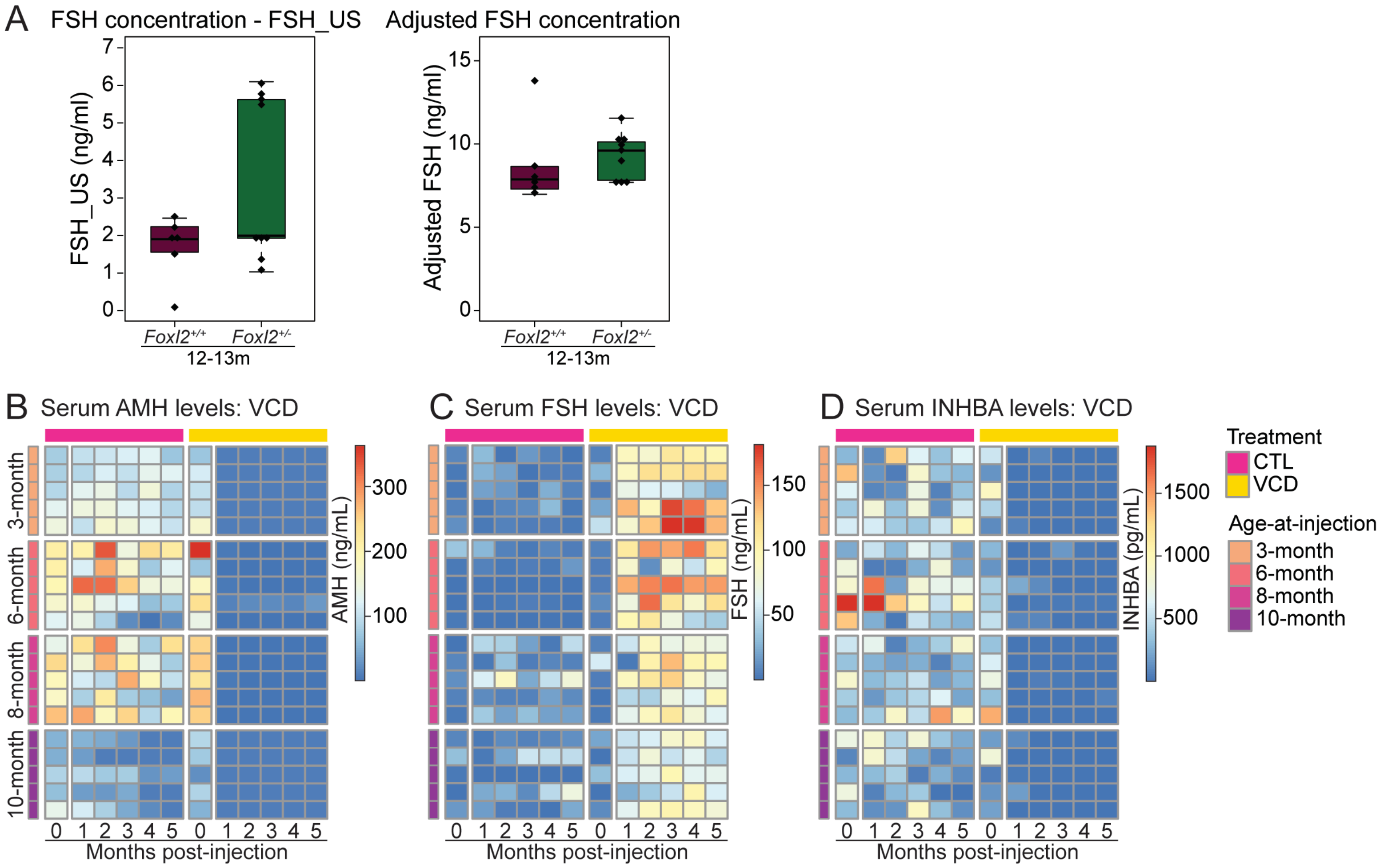
A
FSH concentration - FSH_US
Adjusted FSH concentration
FSH_US (ng/ml)
Adjusted FSH (ng/ml)
Foxl2+/+
Foxl2+/-
12-13m
B Serum AMH levels: VCD
C Serum FSH levels: VCD
D Serum INHBA levels: VCD
3-month
6-month
8-month
10-month
AMH (ng/mL)
FSH (ng/mL)
INHBA (pg/mL)
Months post-injection
Treatment
CTL
VCD
Age-at-injection
3-month
6-month
8-month
10-month

Fig. S4

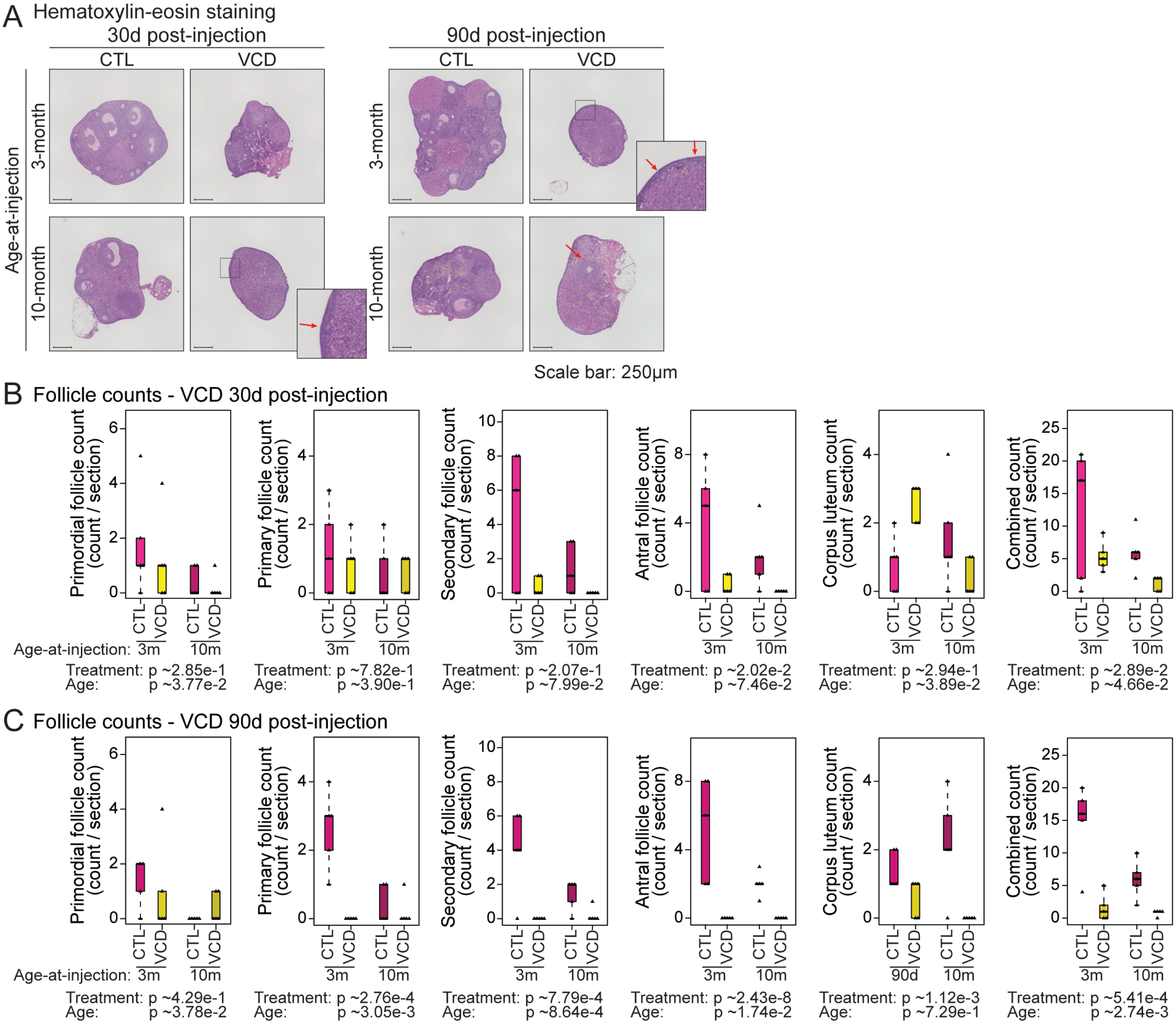

A Hematoxylin-eosin staining
30d post-injection
90d post-injection
CTL
VCD
Age-at-injection
3-month
10-month
Scale bar: 250μm
B Follicle counts - VCD 30d post-injection
Primordial follicle count (count / section)
Primary follicle count (count / section)
Secondary follicle count (count / section)
Antral follicle count (count / section)
Corpus luteum count (count / section)
Combined count (count / section)
Age-at-injection: 3m 10m
Treatment: p ~2.85e-1 Age: p ~3.77e-2
Treatment: p ~7.82e-1 Age: p ~3.90e-1
Treatment: p ~2.07e-1 Age: p ~7.99e-2
Treatment: p ~2.02e-2 Age: p ~7.46e-2
Treatment: p ~2.94e-1 Age: p ~3.89e-2
Treatment: p ~2.89e-2 Age: p ~4.66e-2
C Follicle counts - VCD 90d post-injection
Primordial follicle count (count / section)
Primary follicle count (count / section)
Secondary follicle count (count / section)
Antral follicle count (count / section)
Corpus luteum count (count / section)
Combined count (count / section)
Age-at-injection: 3m 10m
90d
Treatment: p ~4.29e-1 Age: p ~3.78e-2
Treatment: p ~2.76e-4 Age: p ~3.05e-3
Treatment: p ~7.79e-4 Age: p ~8.64e-4
Treatment: p ~2.43e-8 Age: p ~1.74e-2
Treatment: p ~1.12e-3 Age: p ~7.29e-1
Treatment: p ~5.41e-4 Age: p ~2.74e-3

Fig. S5

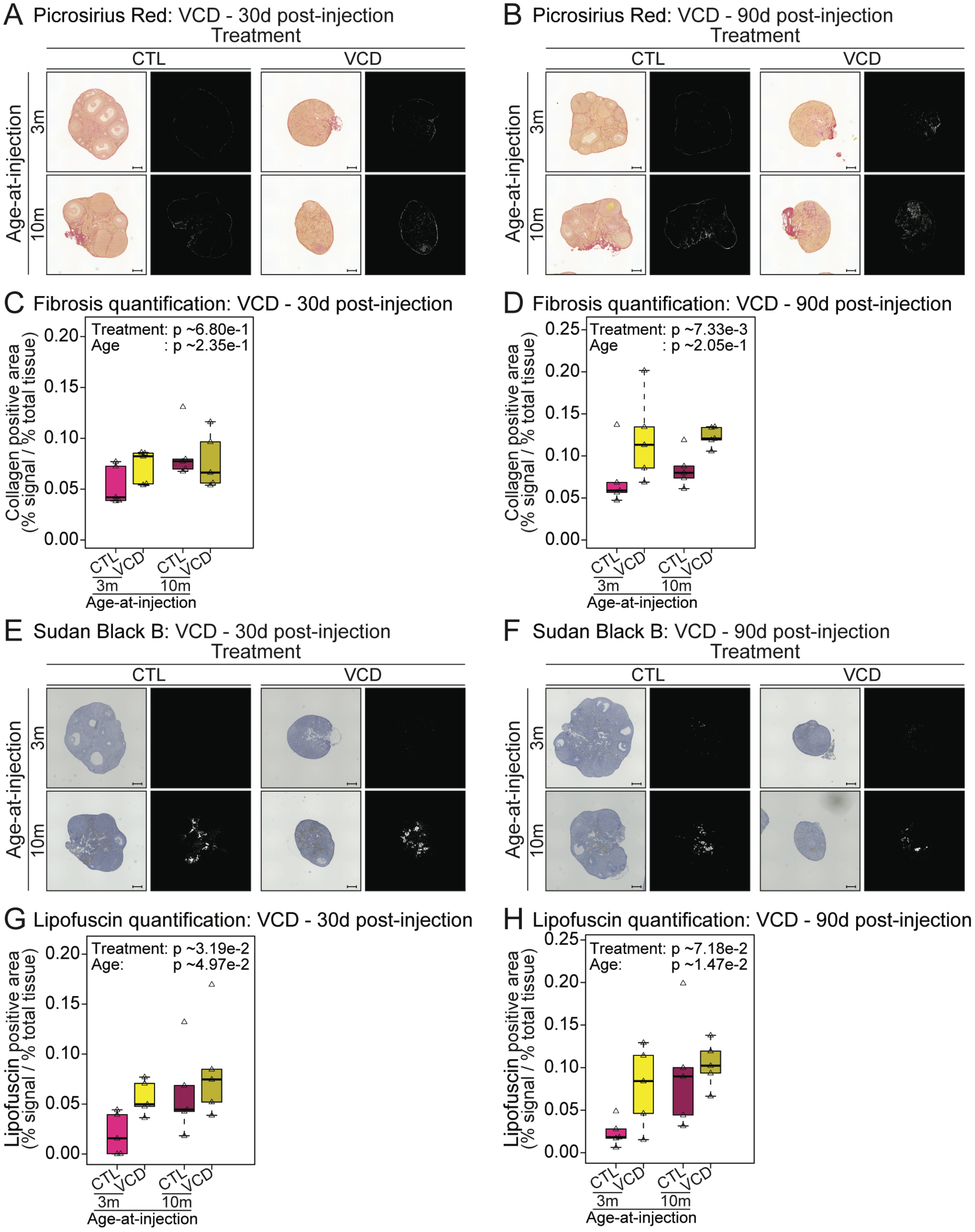
A Picrosirius Red: VCD - 30d post-injection
B Picrosirius Red: VCD - 90d post-injection
Treatment
CTL
VCD
Age-at-injection
3m
10m
C Fibrosis quantification: VCD - 30d post-injection
Treatment: p ~6.80e-1
Age : p ~2.35e-1
D Fibrosis quantification: VCD - 90d post-injection
Treatment: p ~7.33e-3
Age : p ~2.05e-1
Collagen positive area (% signal / % total tissue)
E Sudan Black B: VCD - 30d post-injection
F Sudan Black B: VCD - 90d post-injection
G Lipofuscin quantification: VCD - 30d post-injection
Treatment: p ~3.19e-2
Age: p ~4.97e-2
H Lipofuscin quantification: VCD - 90d post-injection
Treatment: p ~7.18e-2
Age: p ~1.47e-2
Lipofuscin positive area (% signal / % total tissue)
CTL
VCD
3m
10m
Age-at-injection

Fig. S6

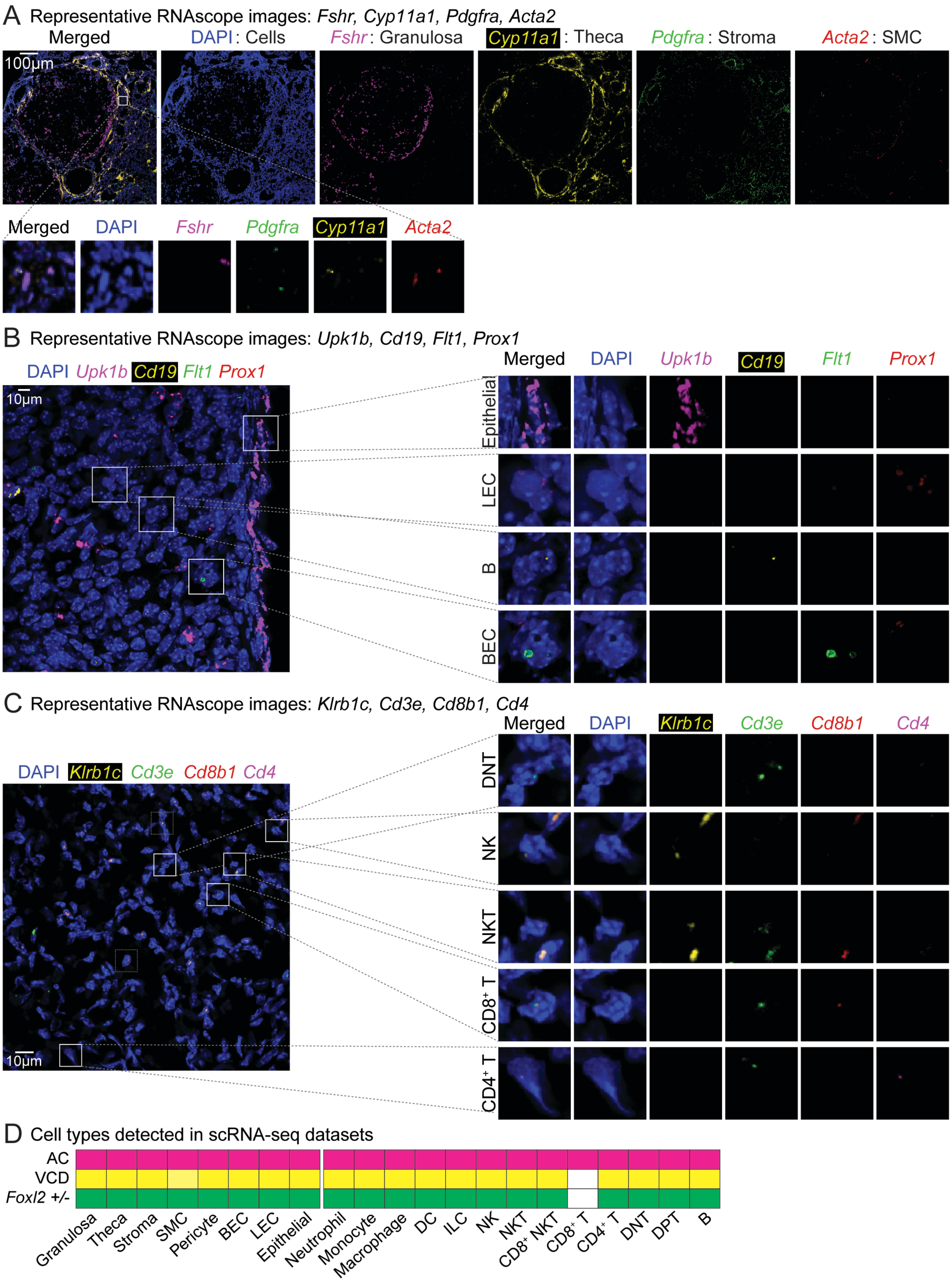
A Representative RNAscope images: Fshr, Cyp11a1, Pdgfra, Acta2
Merged
DAPI: Cells
Fshr: Granulosa
Cyp11a1: Theca
Pdgfra: Stroma
Acta2: SMC
100µm
Merged
DAPI
Fshr
Pdgfra
Cyp11a1
Acta2
B Representative RNAscope images: Upk1b, Cd19, Flt1, Prox1
DAPI Upk1b Cd19 Flt1 Prox1
10µm
Merged
DAPI
Upk1b
Cd19
Flt1
Prox1
Epithelial
LEC
B
BEC
C Representative RNAscope images: Klrb1c, Cd3e, Cd8b1, Cd4
DAPI Klrb1c Cd3e Cd8b1 Cd4
10µm
Merged
DAPI
Klrb1c
Cd3e
Cd8b1
Cd4
DNT
NK
NKT
CD8+ T
CD4+ T
D Cell types detected in scRNA-seq datasets
AC
VCD
Foxl2 +/-
Granulosa
Theca
Stroma
SMC
Pericyte
BEC
LEC
Epithelial
Neutrophil
Monocyte
Macrophage
DC
ILC
NK
NKT
CD8+ NKT
CD8+ T
CD4+ T
DNT
DPT
B

Fig. S7

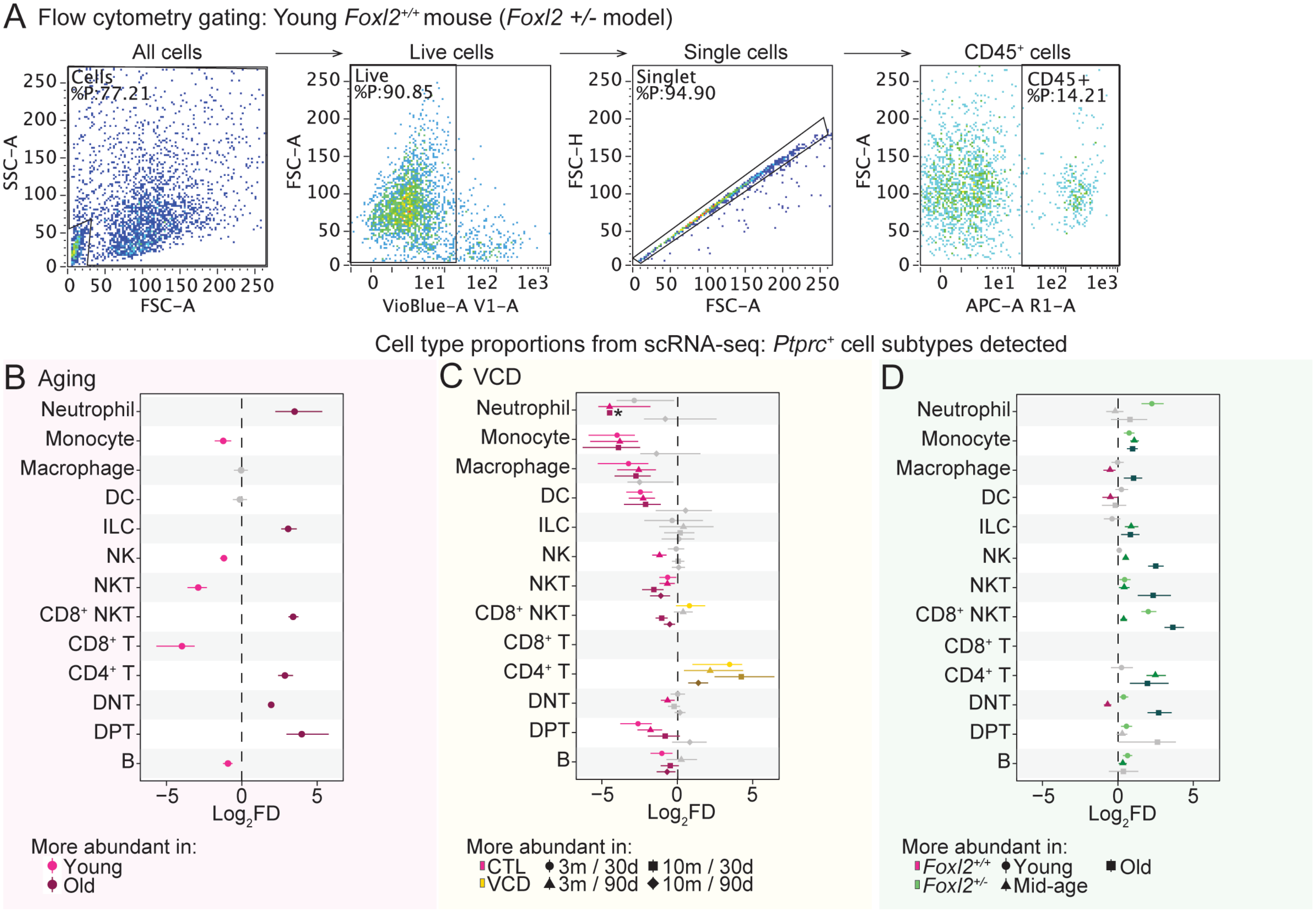

A Flow cytometry gating: Young Foxl2+/+ mouse (Foxl2 +/- model)
All cells
Live cells
Single cells
CD45+ cells
Cells %P:77.21
Live %P:90.85
Singlet %P:94.90
CD45+ %P:14.21
SSC−A
FSC−A
FSC−H
VioBlue−A V1−A
APC−A R1−A
Cell type proportions from scRNA-seq: Ptprc+ cell subtypes detected
B Aging
C VCD
D
Neutrophil
Monocyte
Macrophage
DC
ILC
NK
NKT
CD8+ NKT
CD8+ T
CD4+ T
DNT
DPT
B
Log2FD
More abundant in:
Young
Old
CTL
VCD
3m / 30d
3m / 90d
10m / 30d
10m / 90d
Foxl2+/+
Foxl2+/-
Young
Mid-age
Old

Fig. S8

RNAscope-based cell type abundance analysis: non-immune cells

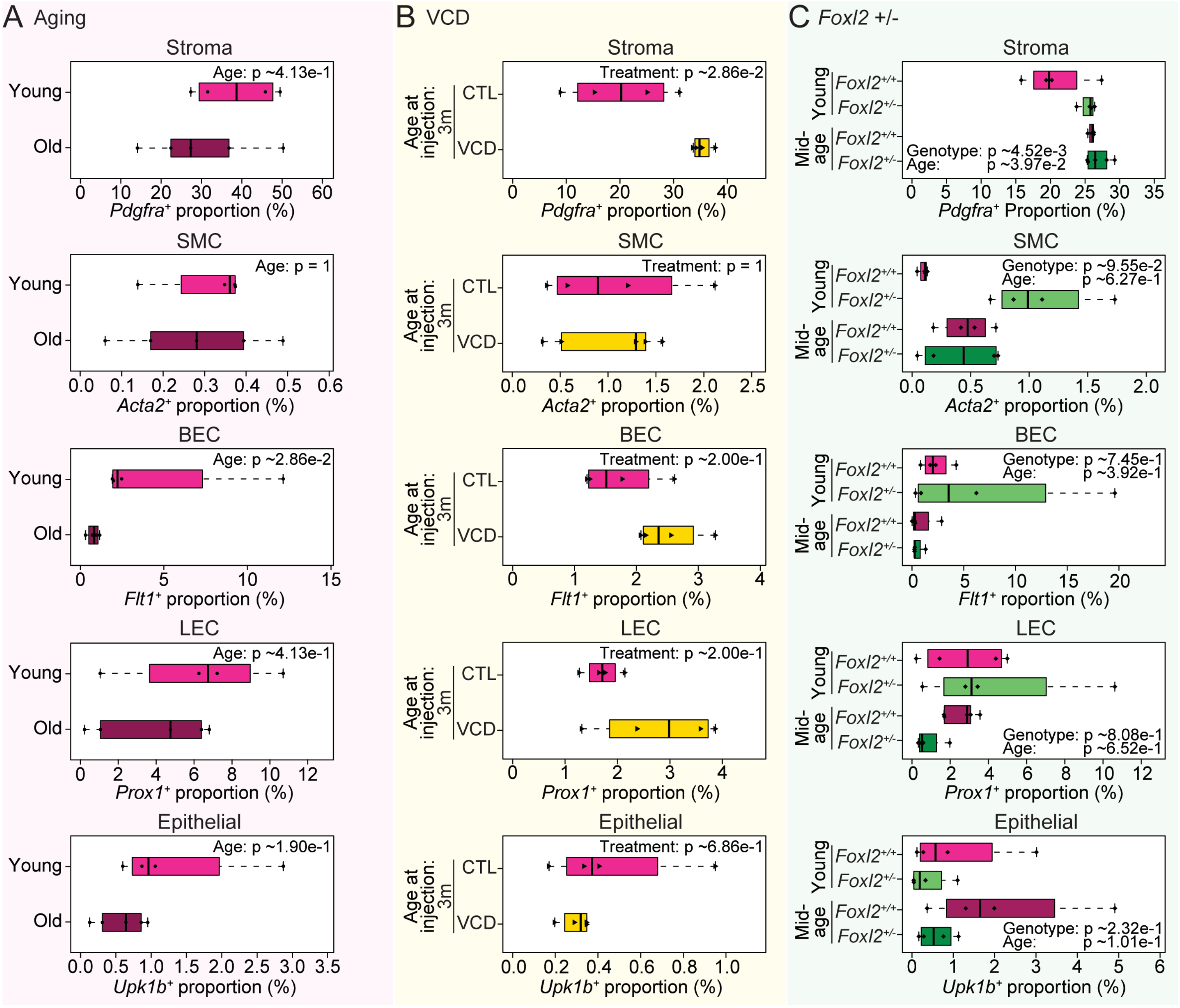

Fig. S9

RNAscope-based cell type abundance analysis: immune cells

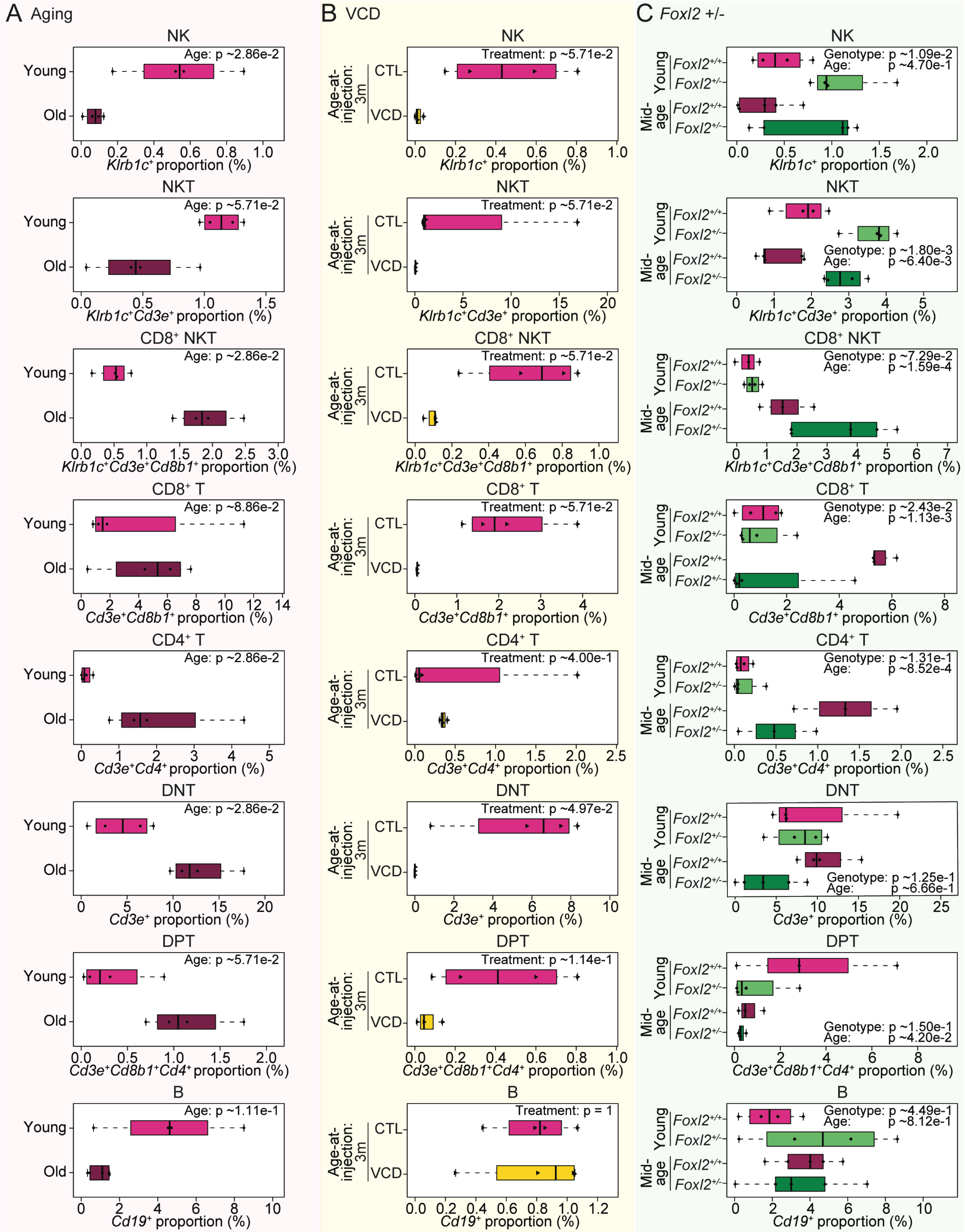

Fig. S10

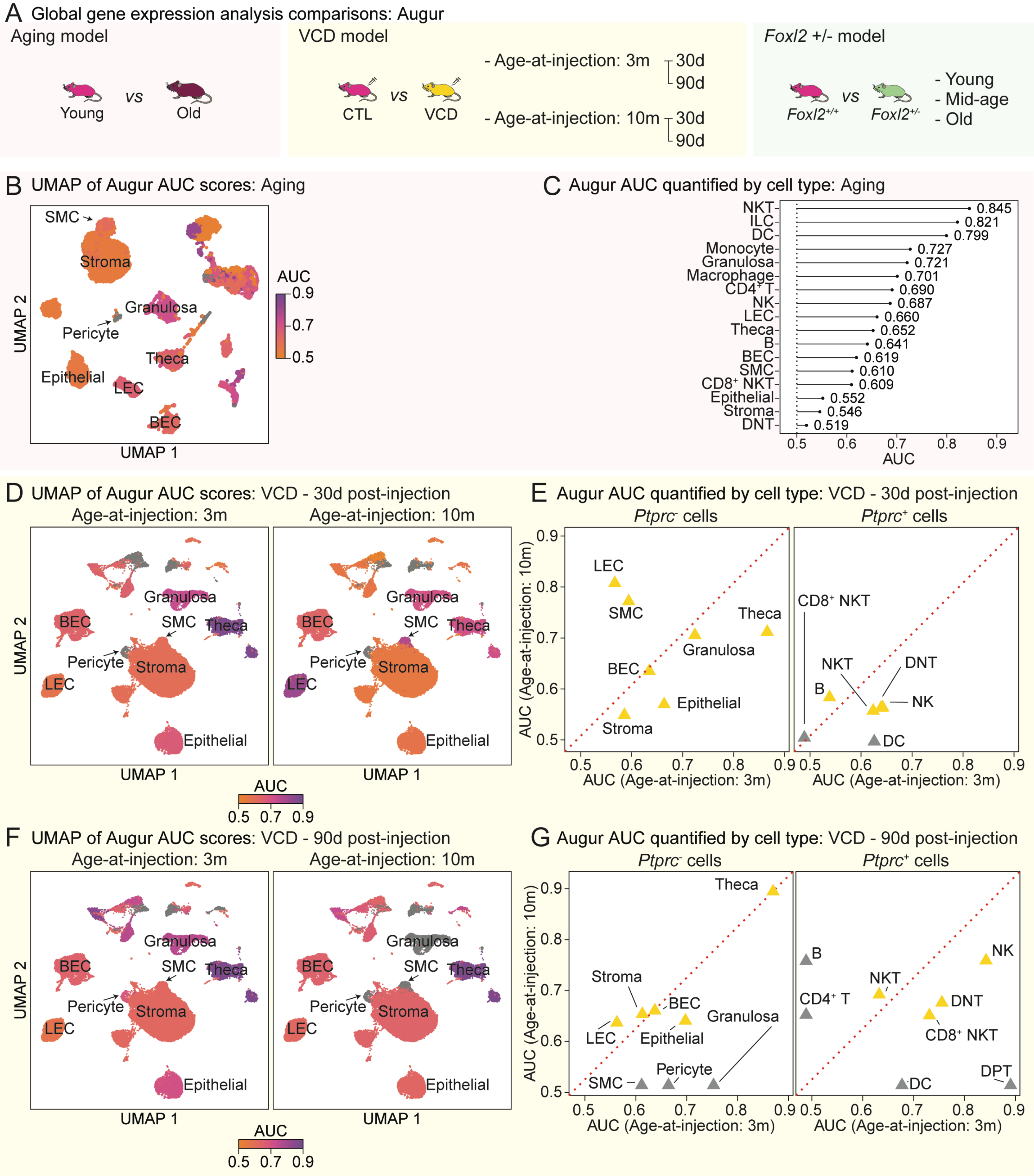

A Global gene expression analysis comparisons: Augur
Aging model
Young vs Old
VCD model
CTL vs VCD
- Age-at-injection: 3m 30d 90d
- Age-at-injection: 10m 30d 90d
Foxl2 +/- model
Foxl2+/+ vs Foxl2+/-
- Young
- Mid-age
- Old
B UMAP of Augur AUC scores: Aging
SMC
Stroma
Granulosa
Pericyte
Theca
Epithelial
LEC
BEC
UMAP 1
UMAP 2
AUC
0.9
0.7
0.5
C Augur AUC quantified by cell type: Aging
NKT 0.845
ILC 0.821
DC 0.799
Monocyte 0.727
Granulosa 0.721
Macrophage 0.701
CD4+ T 0.690
NK 0.687
LEC 0.660
Theca 0.652
B 0.641
BEC 0.619
SMC 0.610
CD8+ NKT 0.609
Epithelial 0.552
Stroma 0.546
DNT 0.519
0.5 0.6 0.7 0.8 0.9
AUC
D UMAP of Augur AUC scores: VCD - 30d post-injection
Age-at-injection: 3m
Age-at-injection: 10m
Granulosa
BEC
SMC
Theca
Pericyte
Stroma
LEC
Epithelial
UMAP 1
UMAP 2
AUC
0.5 0.7 0.9
E Augur AUC quantified by cell type: VCD - 30d post-injection
Ptprc- cells
Ptprc+ cells
LEC
SMC
Theca
Granulosa
BEC
Epithelial
Stroma
CD8+ NKT
NKT
DNT
B
NK
DC
AUC (Age-at-injection: 10m)
AUC (Age-at-injection: 3m)
0.5 0.6 0.7 0.8 0.9
F UMAP of Augur AUC scores: VCD - 90d post-injection
Age-at-injection: 3m
Age-at-injection: 10m
Granulosa
BEC
SMC
Theca
Pericyte
Stroma
LEC
Epithelial
UMAP 1
UMAP 2
AUC
0.5 0.7 0.9
G Augur AUC quantified by cell type: VCD - 90d post-injection
Ptprc- cells
Ptprc+ cells
Theca
Stroma
BEC
Granulosa
LEC
Epithelial
Pericyte
SMC
B
NK
NKT
CD4+ T
DNT
CD8+ NKT
DPT
DC
AUC (Age-at-injection: 10m)
AUC (Age-at-injection: 3m)
0.5 0.6 0.7 0.8 0.9

Fig. S11

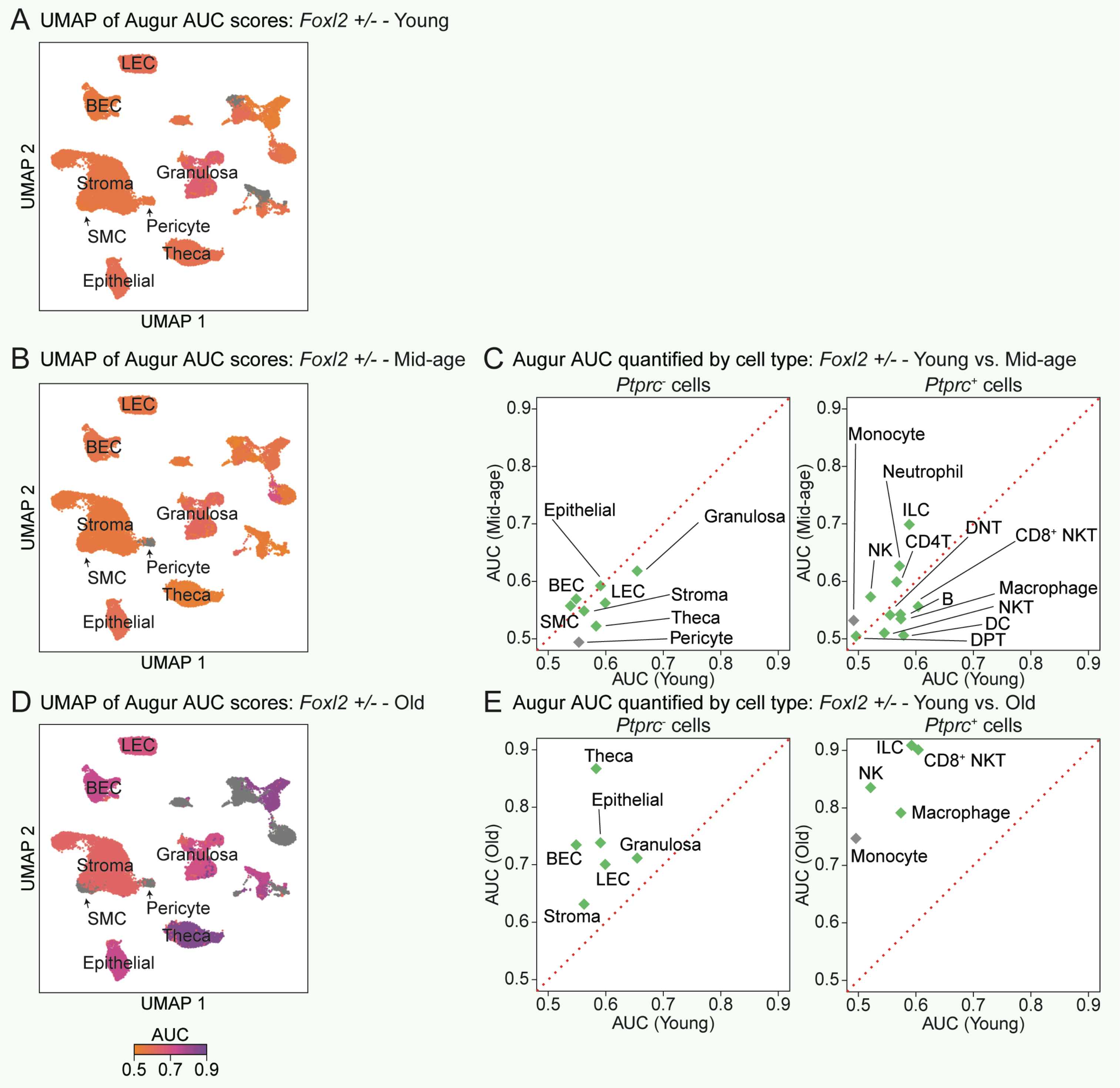

A UMAP of Augur AUC scores: *Foxl2 +/-* - Young
B UMAP of Augur AUC scores: *Foxl2 +/-* - Mid-age
C Augur AUC quantified by cell type: *Foxl2 +/-* - Young vs. Mid-age
D UMAP of Augur AUC scores: *Foxl2 +/-* - Old
E Augur AUC quantified by cell type: *Foxl2 +/-* - Young vs. Old
LEC
BEC
Stroma
Granulosa
SMC
Pericyte
Theca
Epithelial
UMAP 1
UMAP 2
AUC
0.5 0.7 0.9
*Ptprc*- cells
*Ptprc*+ cells
AUC (Young)
AUC (Mid-age)
AUC (Old)
Monocyte
Neutrophil
ILC
NK
CD4T
DNT
CD8+ NKT
Macrophage
B
NKT
DC
DPT
0.5 0.6 0.7 0.8 0.9

Fig. S12

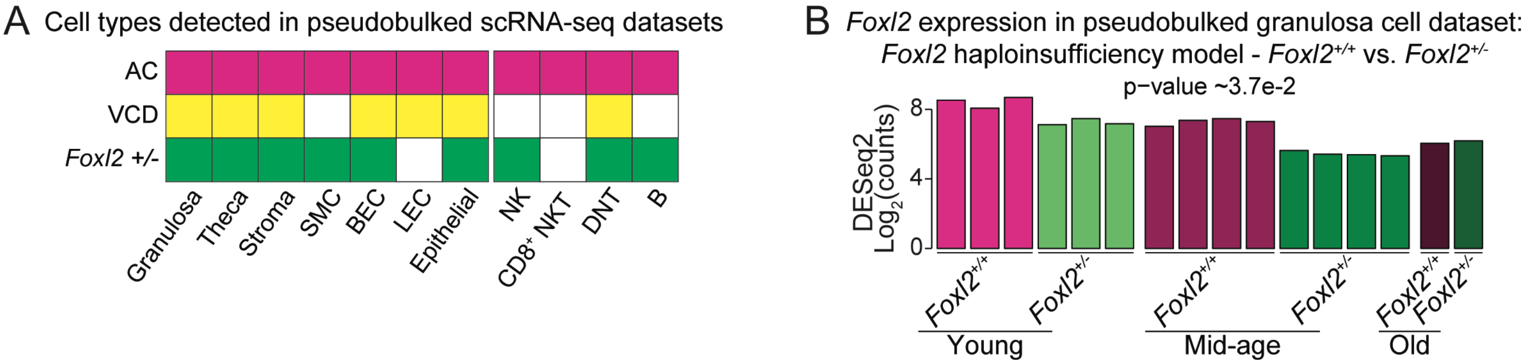
A
Cell types detected in pseudobulked scRNA-seq datasets
AC
VCD
Foxl2 +/-
Granulosa
Theca
Stroma
SMC
BEC
LEC
Epithelial
NK
CD8+ NKT
DNT
B
B
Foxl2 expression in pseudobulked granulosa cell dataset:
Foxl2 haploinsufficiency model - Foxl2+/+ vs. Foxl2+/-
p−value ~3.7e-2
DESeq2
Log2(counts)
8
4
0
Foxl2+/+
Foxl2+/-
Young
Foxl2+/+
Foxl2+/-
Mid-age
Foxl2+/+
Foxl2+/-
Old

Fig. S13

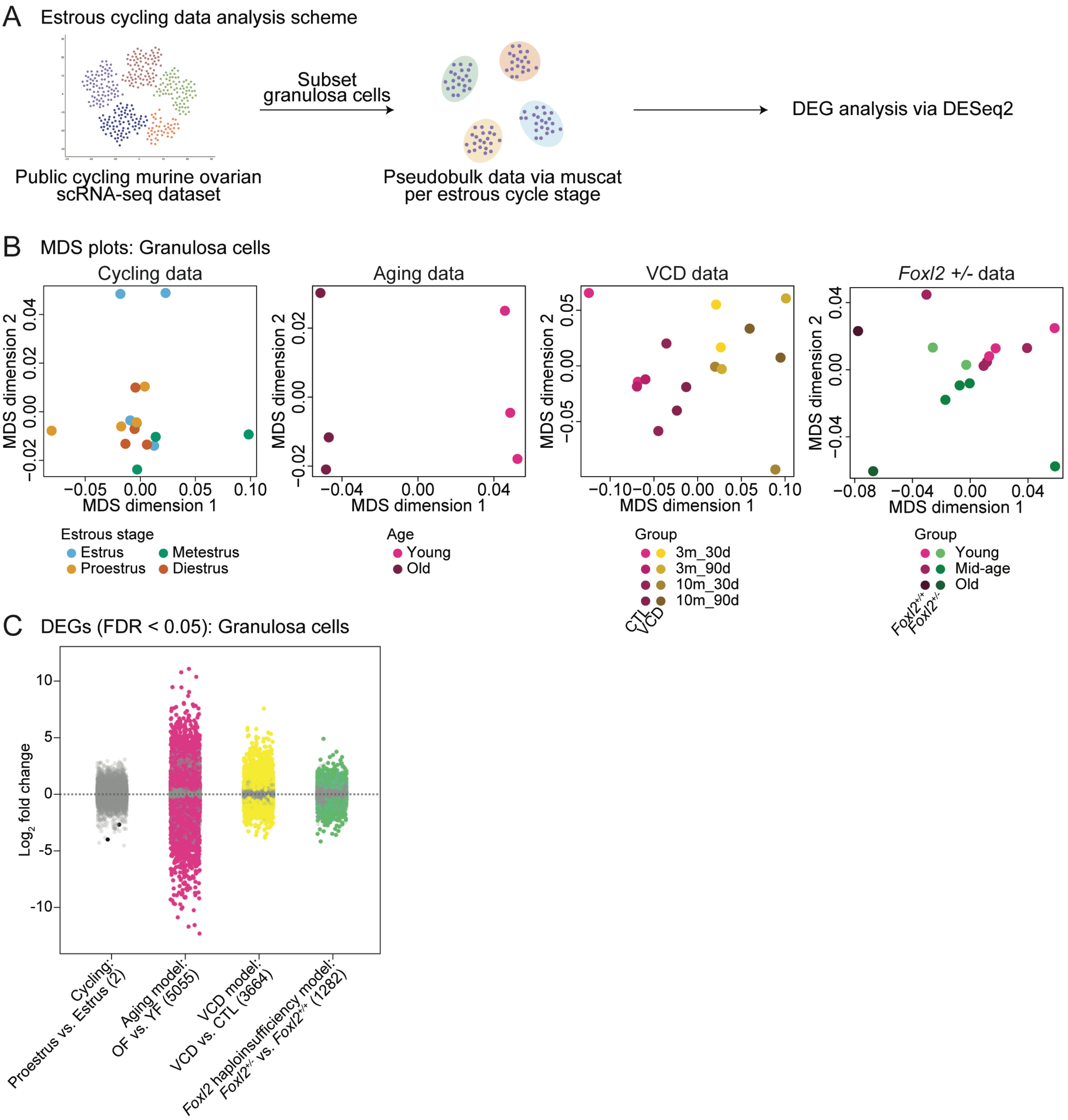
A Estrous cycling data analysis scheme
Public cycling murine ovarian scRNA-seq dataset
Subset granulosa cells
Pseudobulk data via muscat per estrous cycle stage
DEG analysis via DESeq2
B MDS plots: Granulosa cells
Cycling data
Aging data
VCD data
Foxl2 +/- data
MDS dimension 1
MDS dimension 2
Estrous stage
Estrus
Metestrus
Proestrus
Diestrus
Age
Young
Old
Group
3m_30d
3m_90d
10m_30d
10m_90d
CTL
VCD
Young
Mid-age
Old
Foxl2+/+
Foxl2+/-
C DEGs (FDR < 0.05): Granulosa cells
Log2 fold change
Cycling: Proestrus vs. Estrus (2)
Aging model: OF vs. YF (5055)
VCD model: VCD vs. CTL (3664)
Foxl2 haploinsufficiency model: Foxl2+/- vs. Foxl2+/+ (1282)

Fig. S14

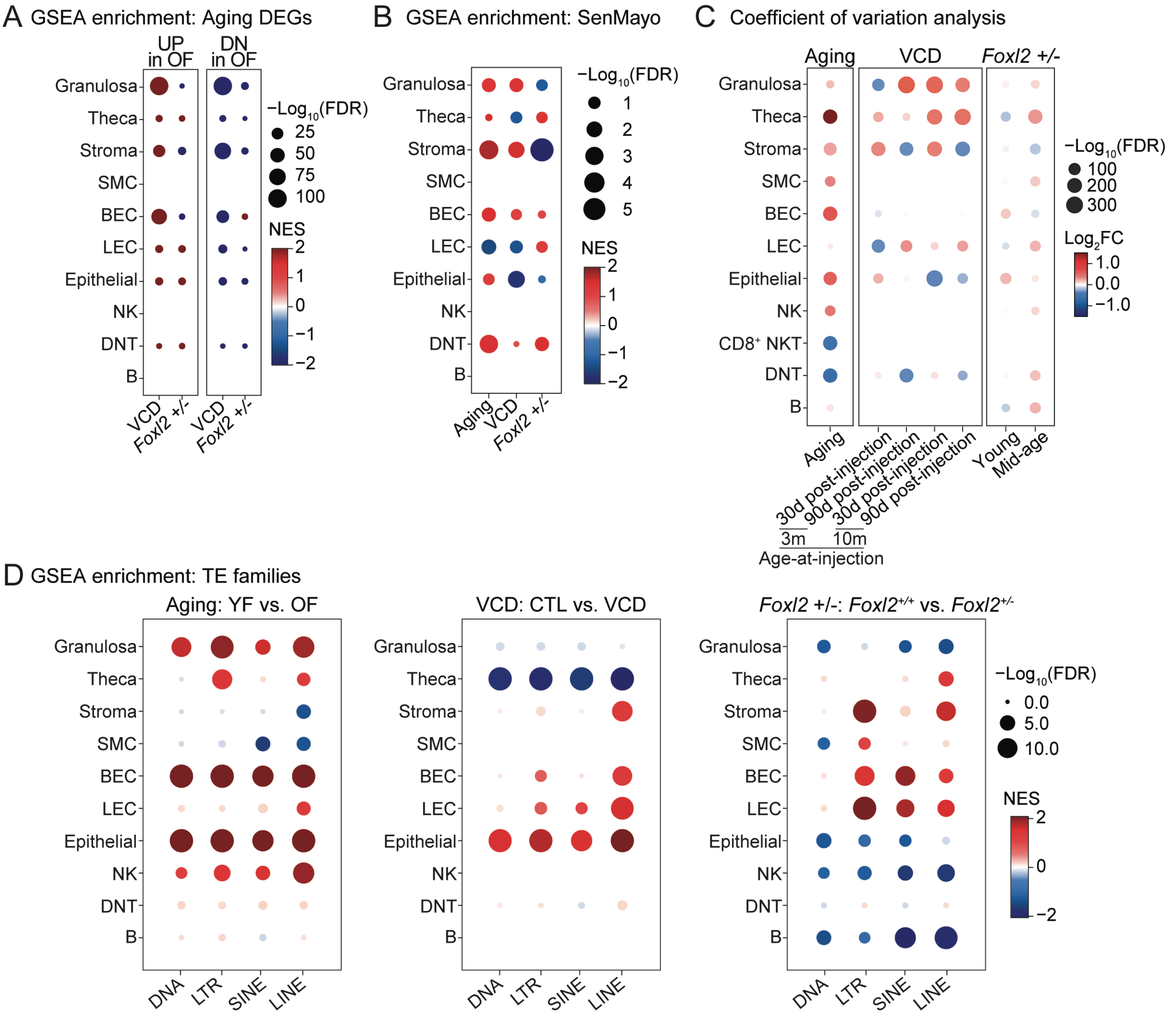
A GSEA enrichment: Aging DEGs
UP in OF
DN in OF
Granulosa
Theca
Stroma
SMC
BEC
LEC
Epithelial
NK
DNT
B
VCD
Foxl2 +/-
VCD
Foxl2 +/-
−Log10(FDR)
25
50
75
100
NES
2
1
0
−1
−2
B GSEA enrichment: SenMayo
Granulosa
Theca
Stroma
SMC
BEC
LEC
Epithelial
NK
DNT
B
Aging
VCD
Foxl2 +/-
−Log10(FDR)
1
2
3
4
5
NES
2
1
0
−1
−2
C Coefficient of variation analysis
Aging
VCD
Foxl2 +/-
Granulosa
Theca
Stroma
SMC
BEC
LEC
Epithelial
NK
CD8+ NKT
DNT
B
Aging
30d post-injection
90d post-injection
30d post-injection
90d post-injection
Young
Mid-age
3m
10m
Age-at-injection
−Log10(FDR)
100
200
300
Log2FC
1.0
0.0
−1.0
D GSEA enrichment: TE families
Aging: YF vs. OF
VCD: CTL vs. VCD
Foxl2 +/-: Foxl2+/+ vs. Foxl2+/-
Granulosa
Theca
Stroma
SMC
BEC
LEC
Epithelial
NK
DNT
B
DNA
LTR
SINE
LINE
−Log10(FDR)
0.0
5.0
10.0
NES
2
0
−2

Fig. S15

A Boxplots of ovarian age-regulated genes (FDR < 1e-5): VCD model, Granulosa cells

Top functional enrichment of ovarian age-regulated genes (GO BP)

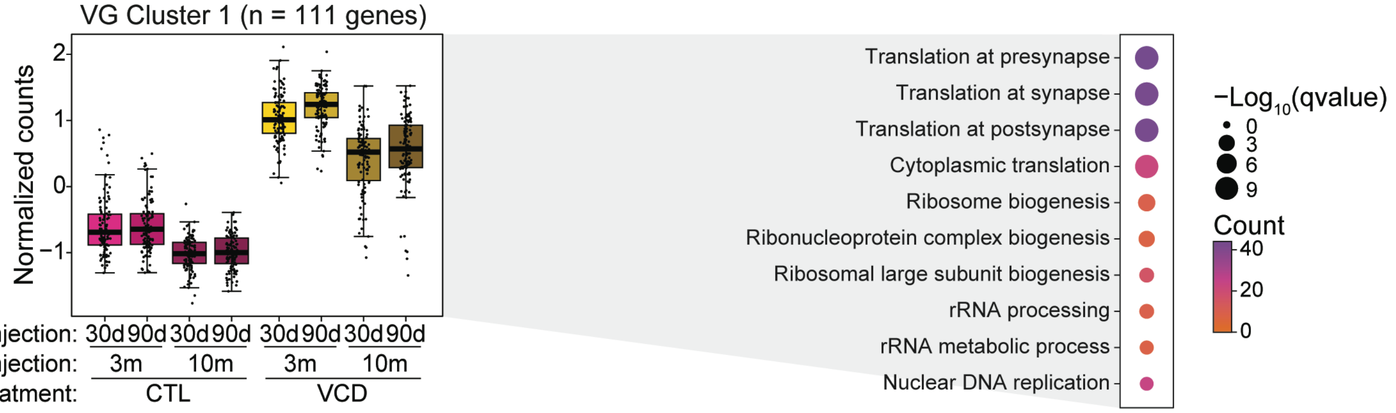


B Boxplots of ovarian age-regulated genes (FDR < 1e-5): *Foxl2* haploinsufficiency model, Granulosa cells

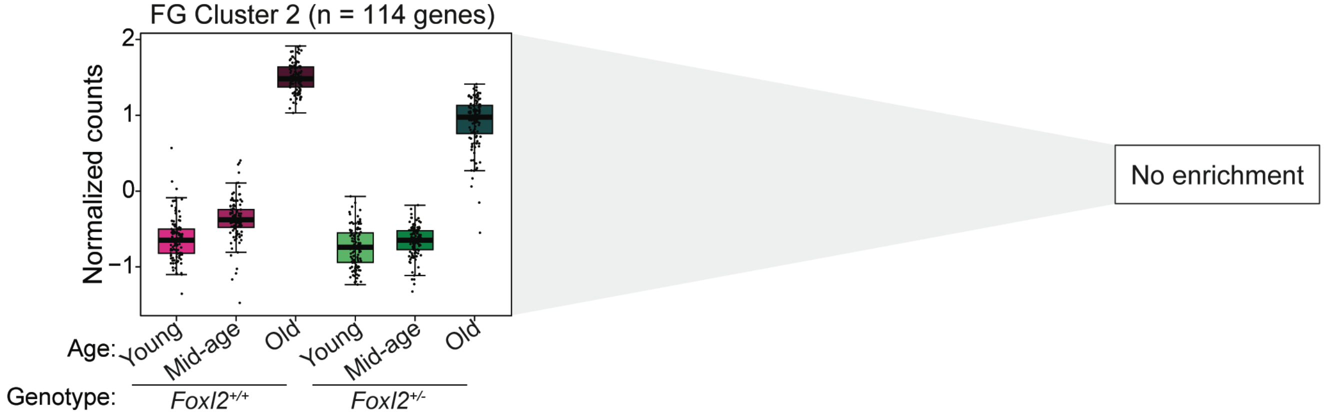

Fig. S16

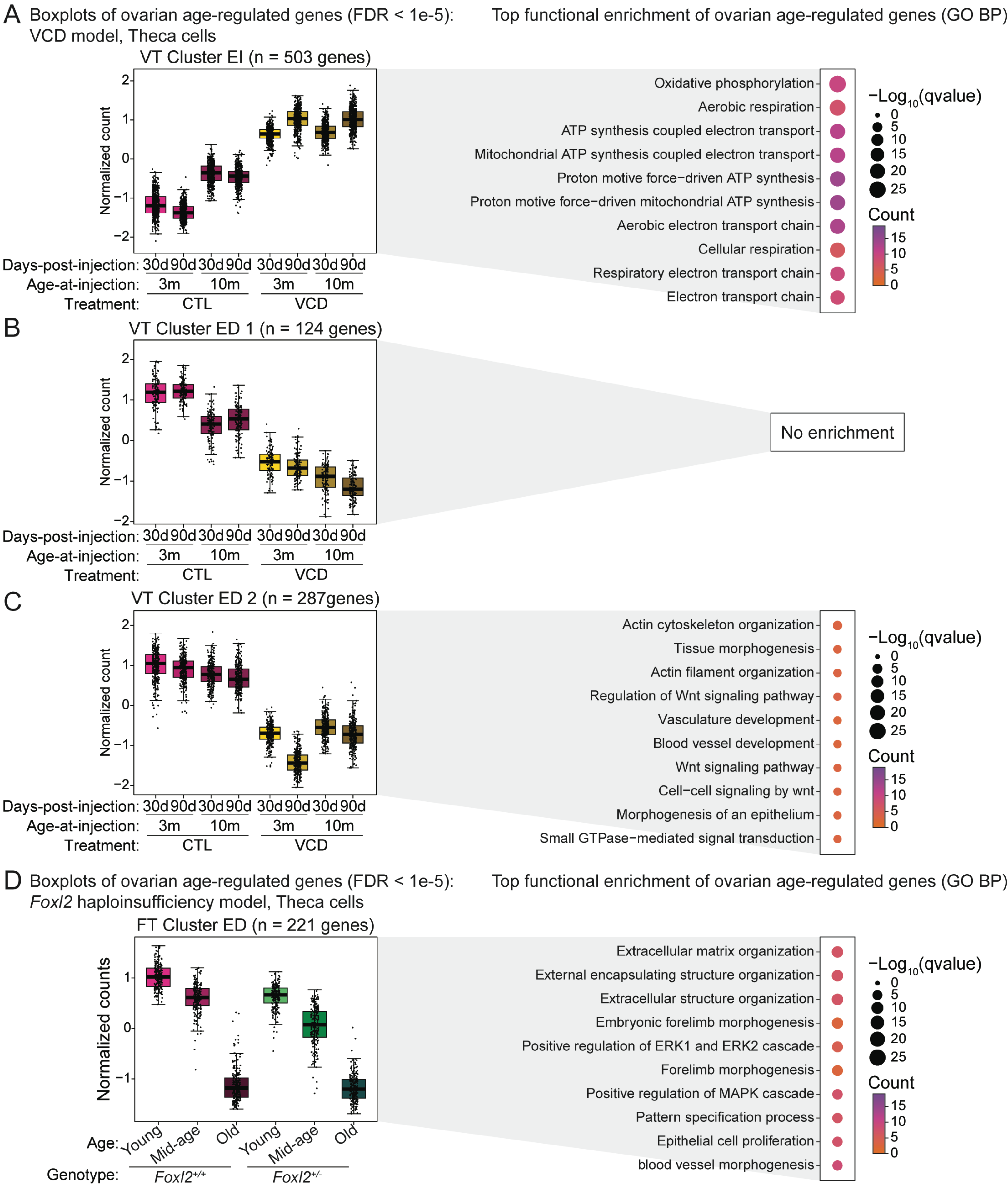
A Boxplots of ovarian age-regulated genes (FDR < 1e-5):
VCD model, Theca cells
Top functional enrichment of ovarian age-regulated genes (GO BP)
VT Cluster EI (n = 503 genes)
Normalized count
Days-post-injection: 30d 90d 30d 90d 30d 90d 30d 90d
Age-at-injection: 3m 10m 3m 10m
Treatment: CTL VCD
Oxidative phosphorylation
Aerobic respiration
ATP synthesis coupled electron transport
Mitochondrial ATP synthesis coupled electron transport
Proton motive force−driven ATP synthesis
Proton motive force−driven mitochondrial ATP synthesis
Aerobic electron transport chain
Cellular respiration
Respiratory electron transport chain
Electron transport chain
−Log10(qvalue)
Count
B
VT Cluster ED 1 (n = 124 genes)
No enrichment
C
VT Cluster ED 2 (n = 287genes)
Actin cytoskeleton organization
Tissue morphogenesis
Actin filament organization
Regulation of Wnt signaling pathway
Vasculature development
Blood vessel development
Wnt signaling pathway
Cell−cell signaling by wnt
Morphogenesis of an epithelium
Small GTPase−mediated signal transduction
D Boxplots of ovarian age-regulated genes (FDR < 1e-5):
Foxl2 haploinsufficiency model, Theca cells
Top functional enrichment of ovarian age-regulated genes (GO BP)
FT Cluster ED (n = 221 genes)
Normalized counts
Age: Young Mid-age Old Young Mid-age Old
Genotype: Foxl2+/+ Foxl2+/-
Extracellular matrix organization
External encapsulating structure organization
Extracellular structure organization
Embryonic forelimb morphogenesis
Positive regulation of ERK1 and ERK2 cascade
Forelimb morphogenesis
Positive regulation of MAPK cascade
Pattern specification process
Epithelial cell proliferation
blood vessel morphogenesis

Fig. S17

A Boxplots of ovarian age-regulated genes (FDR < 1e-5):
*Foxl2* haploinsufficiency model, Theca cells

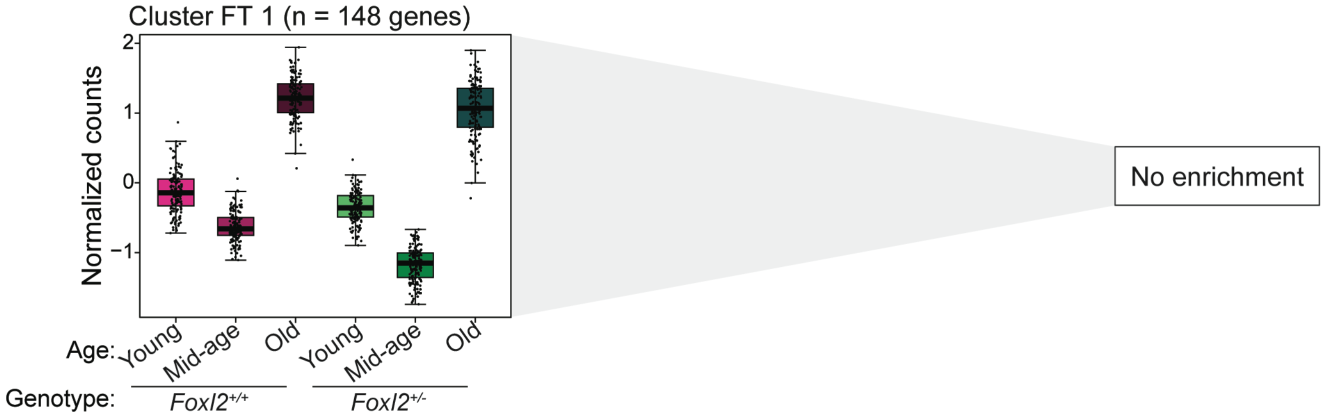


B

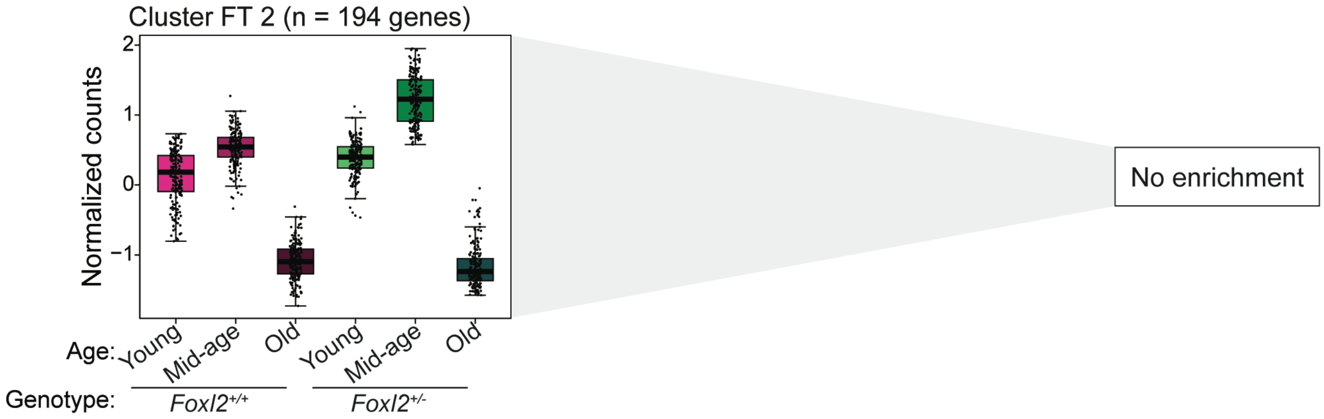

Fig. S18

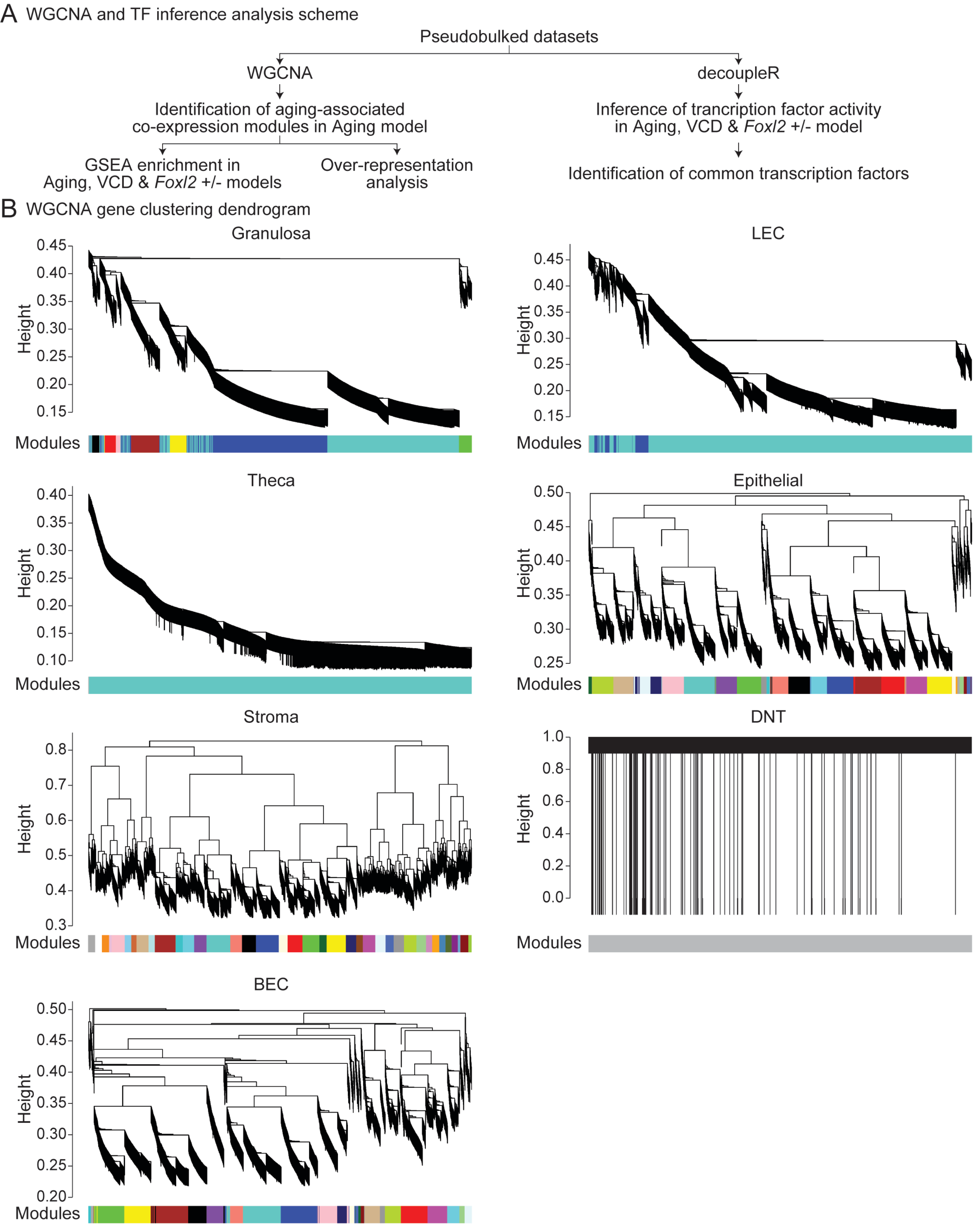
A WGCNA and TF inference analysis scheme
Pseudobulked datasets
WGCNA
decoupleR
Identification of aging-associated co-expression modules in Aging model
Inference of trancription factor activity in Aging, VCD & Foxl2 +/- model
GSEA enrichment in Aging, VCD & Foxl2 +/- models
Over-representation analysis
Identification of common transcription factors
B WGCNA gene clustering dendrogram
Granulosa
LEC
Theca
Epithelial
Stroma
DNT
BEC
Height
Modules

Fig. S19

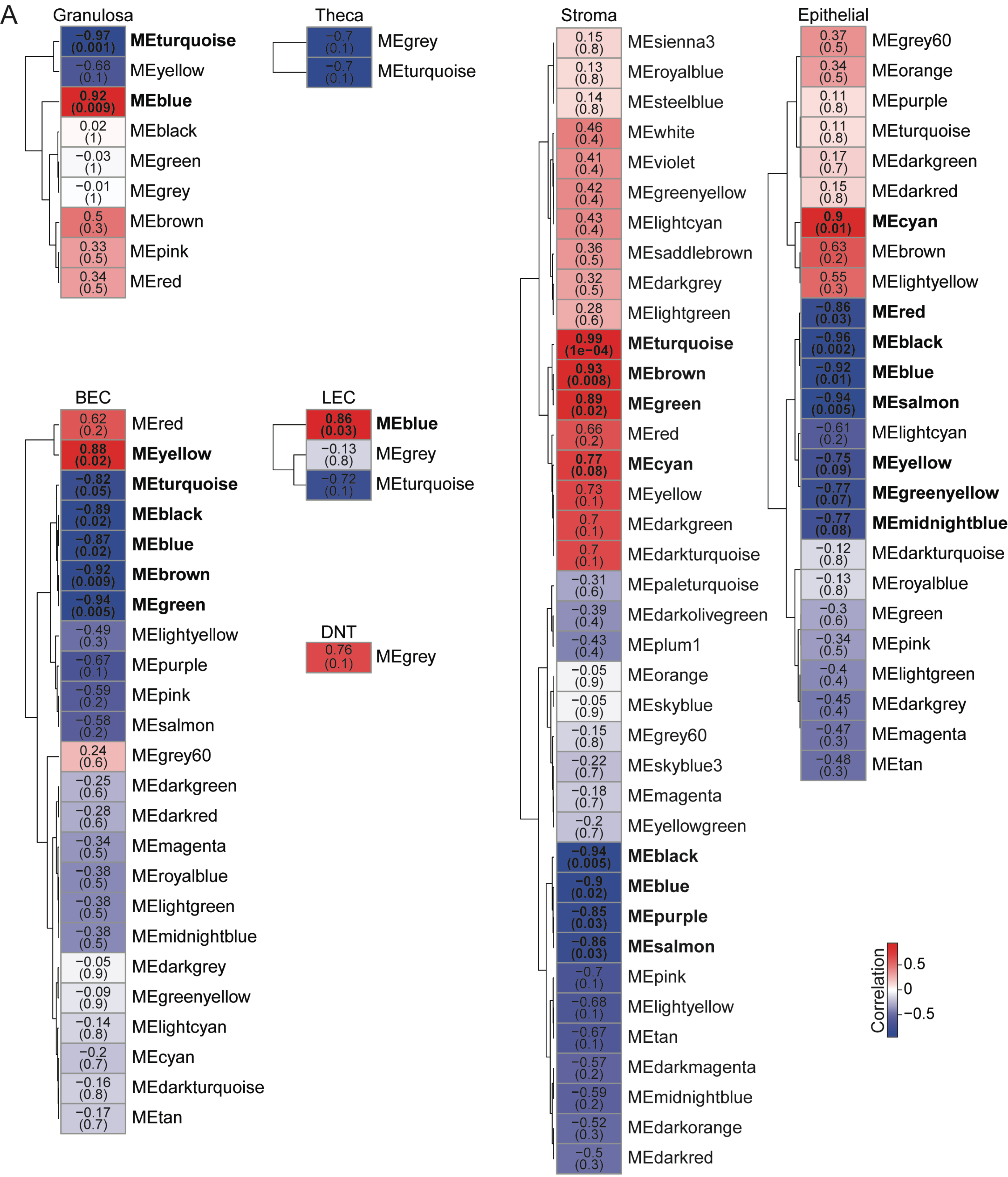
A
Granulosa
-0.97 (0.001) MEturquoise
-0.68 (0.1) MEyellow
0.92 (0.009) MEblue
0.02 (1) MEblack
-0.03 (1) MEgreen
-0.01 (1) MEgrey
0.5 (0.3) MEbrown
0.33 (0.5) MEpink
0.34 (0.5) MEred
Theca
-0.7 (0.1) MEgrey
-0.7 (0.1) MEturquoise
Stroma
0.15 (0.8) MEsienna3
0.13 (0.8) MEroyalblue
0.14 (0.8) MEsteelblue
0.46 (0.4) MEwhite
0.41 (0.4) MEviolet
0.42 (0.4) MEgreenyellow
0.43 (0.4) MElightcyan
0.36 (0.5) MEsaddlebrown
0.32 (0.5) MEdarkgrey
0.28 (0.6) MElightgreen
0.99 (1e-04) MEturquoise
0.93 (0.008) MEbrown
0.89 (0.02) MEgreen
0.66 (0.2) MEred
0.77 (0.08) MEcyan
0.73 (0.1) MEyellow
0.7 (0.1) MEdarkgreen
0.7 (0.1) MEdarkturquoise
-0.31 (0.6) MEpaleturquoise
-0.39 (0.4) MEdarkolivegreen
-0.43 (0.4) MEplum1
-0.05 (0.9) MEorange
-0.05 (0.9) MEskyblue
-0.15 (0.8) MEgrey60
-0.22 (0.7) MEskyblue3
-0.18 (0.7) MEmagenta
-0.2 (0.7) MEyellowgreen
-0.94 (0.005) MEblack
-0.9 (0.02) MEblue
-0.85 (0.03) MEpurple
-0.86 (0.03) MEsalmon
-0.7 (0.1) MEpink
-0.68 (0.1) MElightyellow
-0.67 (0.1) MEtan
-0.57 (0.2) MEdarkmagenta
-0.59 (0.2) MEmidnightblue
-0.52 (0.3) MEdarkorange
-0.5 (0.3) MEdarkred
Epithelial
0.37 (0.5) MEgrey60
0.34 (0.5) MEorange
0.11 (0.8) MEpurple
0.11 (0.8) MEturquoise
0.17 (0.7) MEdarkgreen
0.15 (0.8) MEdarkred
0.9 (0.01) MEcyan
0.63 (0.2) MEbrown
0.55 (0.3) MElightyellow
-0.86 (0.03) MEred
-0.96 (0.002) MEblack
-0.92 (0.01) MEblue
-0.94 (0.005) MEsalmon
-0.61 (0.2) MElightcyan
-0.75 (0.09) MEyellow
-0.77 (0.07) MEgreenyellow
-0.77 (0.08) MEmidnightblue
-0.12 (0.8) MEdarkturquoise
-0.13 (0.8) MEroyalblue
-0.3 (0.6) MEgreen
-0.34 (0.5) MEpink
-0.4 (0.4) MElightgreen
-0.45 (0.4) MEdarkgrey
-0.47 (0.3) MEmagenta
-0.48 (0.3) MEtan
BEC
0.62 (0.2) MEred
0.88 (0.02) MEyellow
-0.82 (0.05) MEturquoise
-0.89 (0.02) MEblack
-0.87 (0.02) MEblue
-0.92 (0.009) MEbrown
-0.94 (0.005) MEgreen
-0.49 (0.3) MElightyellow
-0.67 (0.1) MEpurple
-0.59 (0.2) MEpink
-0.58 (0.2) MEsalmon
0.24 (0.6) MEgrey60
-0.25 (0.6) MEdarkgreen
-0.28 (0.6) MEdarkred
-0.34 (0.5) MEmagenta
-0.38 (0.5) MEroyalblue
-0.38 (0.5) MElightgreen
-0.38 (0.5) MEmidnightblue
-0.05 (0.9) MEdarkgrey
-0.09 (0.9) MEgreenyellow
-0.14 (0.8) MElightcyan
-0.2 (0.7) MEcyan
-0.16 (0.8) MEdarkturquoise
-0.17 (0.7) MEtan
LEC
0.86 (0.03) MEblue
-0.13 (0.8) MEgrey
-0.72 (0.1) MEturquoise
DNT
0.76 (0.1) MEgrey
Correlation
0.5
0
-0.5

Fig. S20

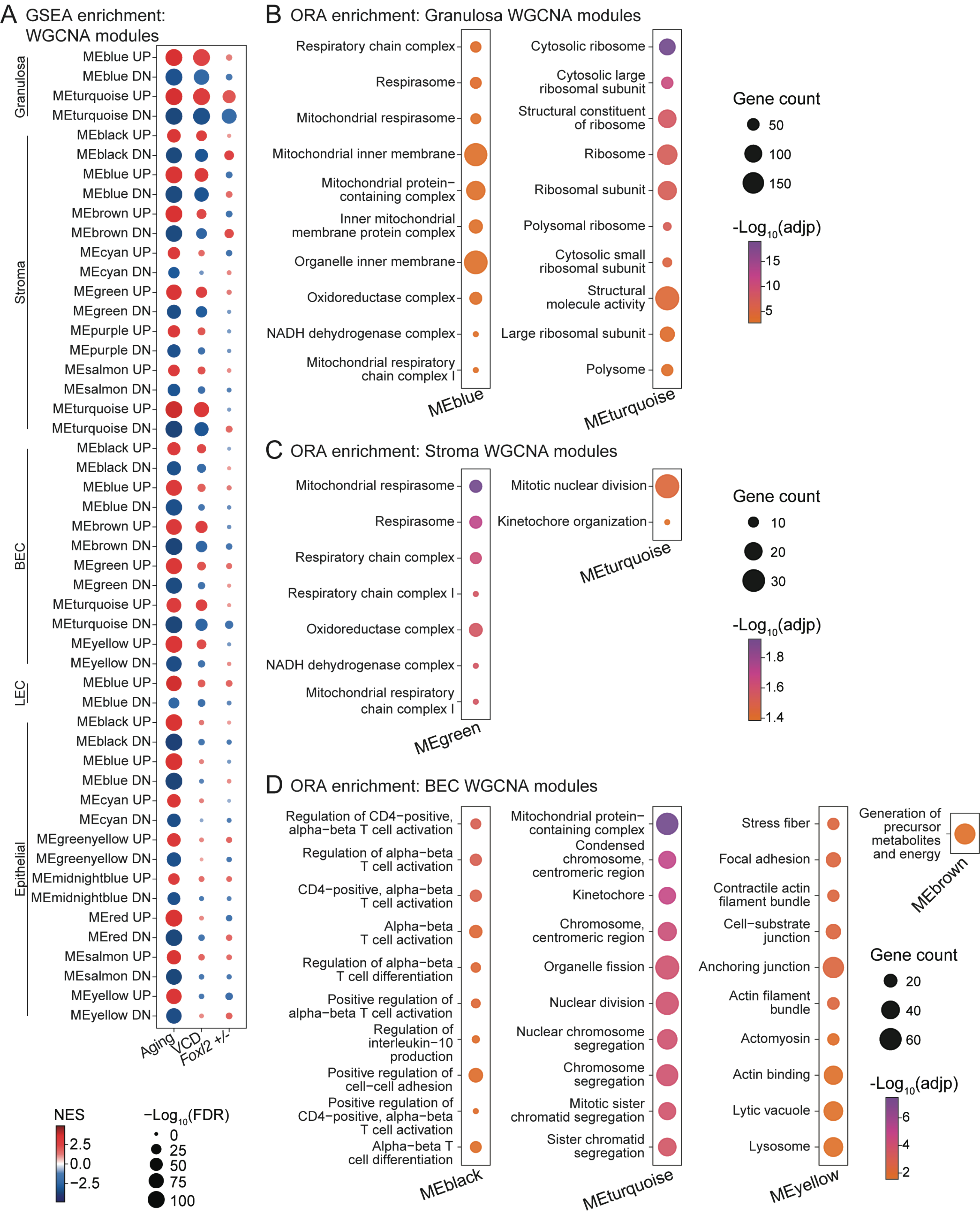

A GSEA enrichment: WGCNA modules
Granulosa
Stroma
BEC
LEC
Epithelial
MEblue UP
MEblue DN
MEturquoise UP
MEturquoise DN
MEblack UP
MEblack DN
MEbrown UP
MEbrown DN
MEcyan UP
MEcyan DN
MEgreen UP
MEgreen DN
MEpurple UP
MEpurple DN
MEsalmon UP
MEsalmon DN
MEyellow UP
MEyellow DN
MEgreenyellow UP
MEgreenyellow DN
MEmidnightblue UP
MEmidnightblue DN
MEred UP
MEred DN
Aging
VCD
Foxl2 +/-
NES
2.5
0.0
−2.5
−Log10(FDR)
0
25
50
75
100
B ORA enrichment: Granulosa WGCNA modules
Respiratory chain complex
Respirasome
Mitochondrial respirasome
Mitochondrial inner membrane
Mitochondrial protein−containing complex
Inner mitochondrial membrane protein complex
Organelle inner membrane
Oxidoreductase complex
NADH dehydrogenase complex
Mitochondrial respiratory chain complex I
MEblue
Cytosolic ribosome
Cytosolic large ribosomal subunit
Structural constituent of ribosome
Ribosome
Ribosomal subunit
Polysomal ribosome
Cytosolic small ribosomal subunit
Structural molecule activity
Large ribosomal subunit
Polysome
MEturquoise
Gene count
50
100
150
-Log10(adjp)
15
10
5
C ORA enrichment: Stroma WGCNA modules
Mitochondrial respirasome
Respirasome
Respiratory chain complex
Respiratory chain complex I
Oxidoreductase complex
NADH dehydrogenase complex
Mitochondrial respiratory chain complex I
MEgreen
Mitotic nuclear division
Kinetochore organization
MEturquoise
Gene count
10
20
30
-Log10(adjp)
1.8
1.6
1.4
D ORA enrichment: BEC WGCNA modules
Regulation of CD4−positive, alpha−beta T cell activation
Regulation of alpha−beta T cell activation
CD4−positive, alpha−beta T cell activation
Alpha−beta T cell activation
Regulation of alpha−beta T cell differentiation
Positive regulation of alpha−beta T cell activation
Regulation of interleukin−10 production
Positive regulation of cell−cell adhesion
Positive regulation of CD4−positive, alpha−beta T cell activation
Alpha−beta T cell differentiation
MEblack
Mitochondrial protein−containing complex
Condensed chromosome, centromeric region
Kinetochore
Chromosome, centromeric region
Organelle fission
Nuclear division
Nuclear chromosome segregation
Chromosome segregation
Mitotic sister chromatid segregation
Sister chromatid segregation
MEturquoise
Stress fiber
Focal adhesion
Contractile actin filament bundle
Cell−substrate junction
Anchoring junction
Actin filament bundle
Actomyosin
Actin binding
Lytic vacuole
Lysosome
MEyellow
Generation of precursor metabolites and energy
MEbrown
Gene count
20
40
60
-Log10(adjp)
6
4
2

Fig. S21

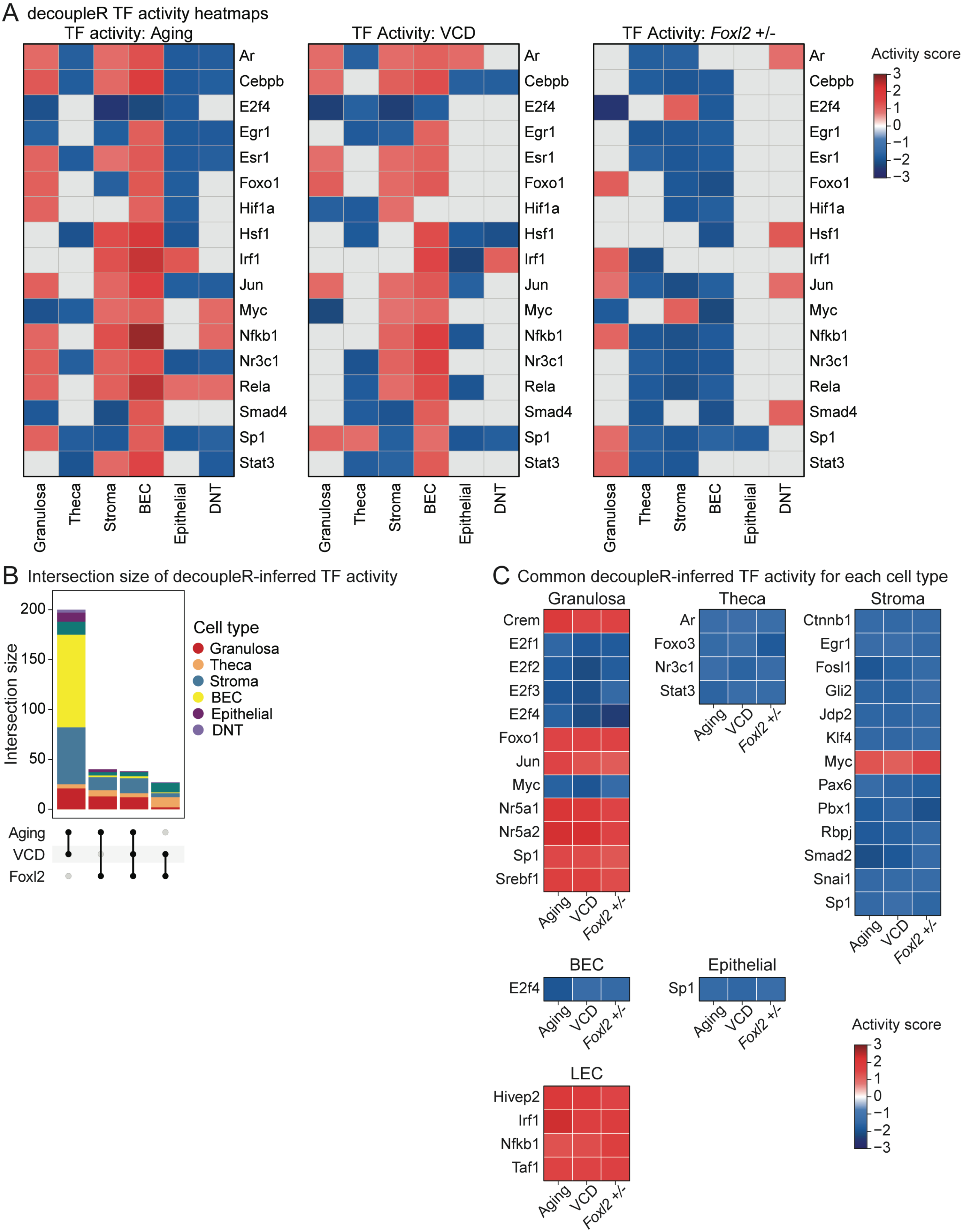
A decoupleR TF activity heatmaps
TF activity: Aging
TF Activity: VCD
TF Activity: *Foxl2* +/-
Ar
Cebpb
E2f4
Egr1
Esr1
Foxo1
Hif1a
Hsf1
Irf1
Jun
Myc
Nfkb1
Nr3c1
Rela
Smad4
Sp1
Stat3
Granulosa
Theca
Stroma
BEC
Epithelial
DNT
Activity score
3
2
1
0
−1
−2
−3
B Intersection size of decoupleR-inferred TF activity
Intersection size
200
150
100
50
0
Cell type
Granulosa
Theca
Stroma
BEC
Epithelial
DNT
Aging
VCD
Foxl2
C Common decoupleR-inferred TF activity for each cell type
Granulosa
Crem
E2f1
E2f2
E2f3
E2f4
Foxo1
Jun
Myc
Nr5a1
Nr5a2
Sp1
Srebf1
Theca
Ar
Foxo3
Nr3c1
Stat3
Stroma
Ctnnb1
Egr1
Fosl1
Gli2
Jdp2
Klf4
Myc
Pax6
Pbx1
Rbpj
Smad2
Snai1
Sp1
BEC
E2f4
Epithelial
Sp1
LEC
Hivep2
Irf1
Nfkb1
Taf1
Aging
VCD
*Foxl2* +/-
Activity score
3
2
1
0
−1
−2
−3

Fig. S22

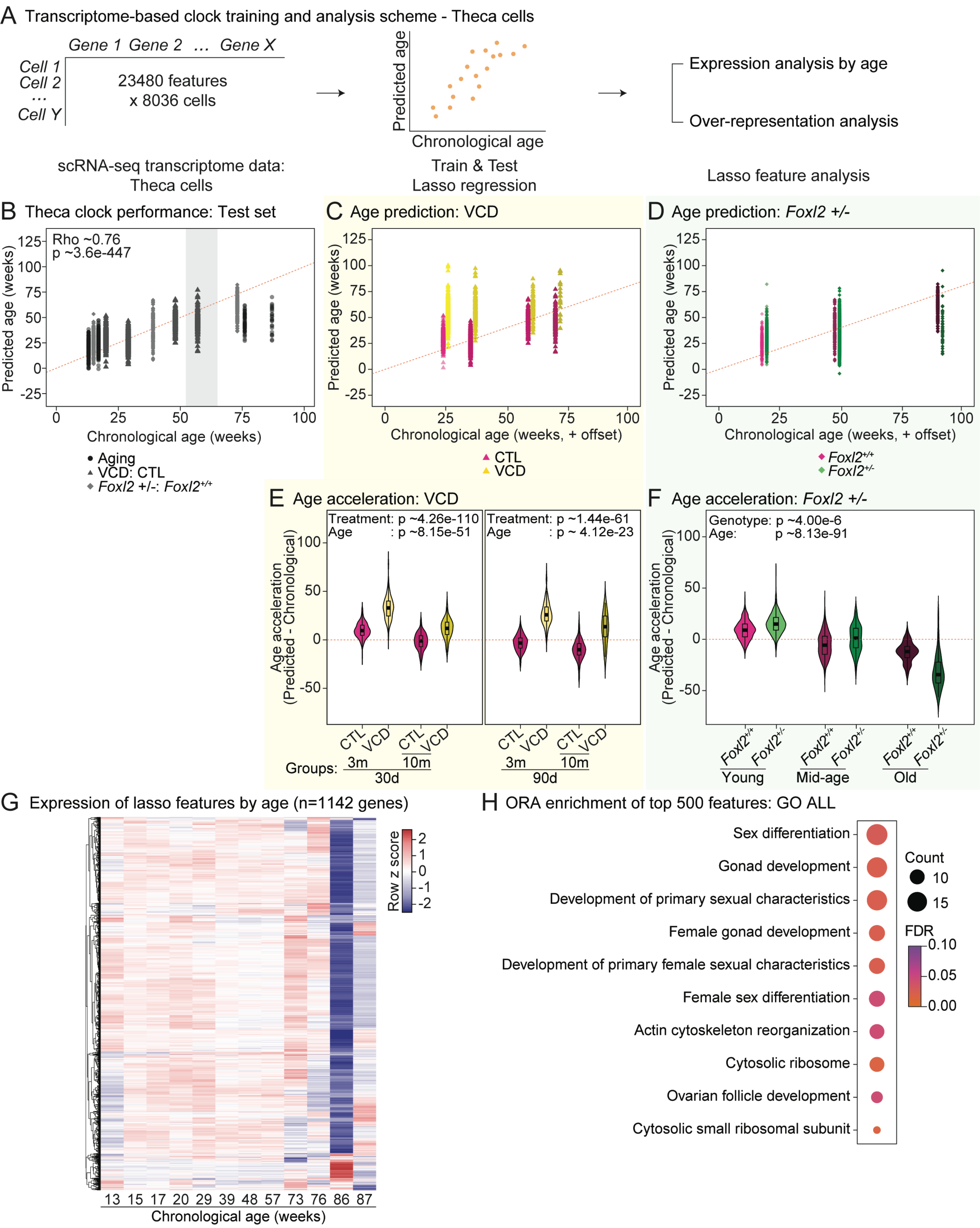
A Transcriptome-based clock training and analysis scheme - Theca cells
Gene 1 Gene 2 … Gene X
Cell 1
Cell 2
…
Cell Y
23480 features x 8036 cells
scRNA-seq transcriptome data: Theca cells
Predicted age
Chronological age
Train & Test Lasso regression
Expression analysis by age
Over-representation analysis
Lasso feature analysis
B Theca clock performance: Test set
Rho ~0.76
p ~3.6e-447
Predicted age (weeks)
Chronological age (weeks)
Aging
VCD: CTL
Foxl2 +/-: Foxl2+/+
C Age prediction: VCD
Chronological age (weeks, + offset)
CTL
VCD
D Age prediction: Foxl2 +/-
Foxl2+/+
Foxl2+/-
E Age acceleration: VCD
Treatment: p ~4.26e-110
Age : p ~8.15e-51
Treatment: p ~1.44e-61
Age : p ~ 4.12e-23
Age acceleration (Predicted - Chronological)
Groups:
CTL VCD CTL VCD
3m 10m
30d
90d
F Age acceleration: Foxl2 +/-
Genotype: p ~4.00e-6
Age: p ~8.13e-91
Young
Mid-age
Old
G Expression of lasso features by age (n=1142 genes)
Row z score
13 15 17 20 29 39 48 57 73 76 86 87
Chronological age (weeks)
H ORA enrichment of top 500 features: GO ALL
Sex differentiation
Gonad development
Development of primary sexual characteristics
Female gonad development
Development of primary female sexual characteristics
Female sex differentiation
Actin cytoskeleton reorganization
Cytosolic ribosome
Ovarian follicle development
Cytosolic small ribosomal subunit
Count
FDR
